# Predictive Cellular Signatures from Live Human Motor Neurons Distinguish TDP-43 ALS and Enable ALS Subtype Stratification

**DOI:** 10.64898/2026.04.22.719920

**Authors:** Julia Kaye, Naufa Amirani, Úna Chan, Noura Al Bistami, Zohreh Faghihmonzavi, Monika Ahirwar, Reuben Thomas, Wesley Robinson, Edward Vertudes, Krishna Raja, Mariya Barch, Drew Linsley, Ana Jovicic, Steven Finkbeiner

## Abstract

Amyotrophic lateral sclerosis (ALS) is a fatal neurodegenerative disorder characterized by the progressive, rapid deterioration of motor neurons (MNs). Rare mutations in a handful of genes are sufficient to cause ALS; however, 90% of ALS cases are not linked to these genes and their underlying cause remains unknown. Abnormal subcellular distribution, structure or aggregation of the TDP-43 protein are nearly universal hallmarks of the disease, suggesting a shared molecular mechanism across both genetic and sporadic ALS (sALS). However, the heterogeneity of the ALS clinical syndrome suggests that the underlying mechanisms culminating in ALS and TDP-43 pathology may partly differ among individuals and may need to be understood to develop successful therapies that target subgroups of patients. Here, we harnessed the power of machine learning (ML) to begin to decode, in a systematic and unbiased fashion, the cellular signatures of ALS. We used high-content imaging of live, human iPSC-derived motor neurons (iMNs) from ALS patients or gene-edited and gene-corrected TDP-43 mutant lines to train shallow connected ML algorithms (SMLs) and deep convolutional neural networks (DNNs). Our models identified and distinguished mutant and control iMNs with moderately high accuracy. We then used explainability methods to uncover the discriminating cellular signals and found that the strongest ones mapped to the nuclear area, suggesting underlying alterations within the nucleus. We validated this finding by revealing that TDP-43 mutant iMNs display alterations in nucleocytoplasmic shuttling and cellular integrity. Further, a time-interaction ML model uncovered dynamic morphological transitions preceding degeneration, offering a window into early pathogenic events as well as neurodevelopmental changes. Extending our ML pipeline to iMNs with mutations in the ALS gene *C9orf72* or derived from sALS revealed both overlapping and distinguishable signatures, suggesting shared yet distinct mechanistic pathways. Together, these findings establish ML-driven phenotypic profiling as a powerful approach to stratify people with ALS, help disentangle the molecular heterogeneity of ALS and produce a more holistic phenotypic definition in cell-based models, and ultimately find causes and treatments. This strategy offers a scalable and innovative paradigm for uncovering early disease mechanisms not only in ALS but potentially across a spectrum of neurodegenerative and sporadic disorders.

## Introduction

ALS is characterized by progressive degeneration of upper and lower motor neurons (MNs) that innervate skeletal muscle. Loss of motor unit function^1–3^ causes atrophy of skeletal muscle and loss of muscle contraction, which eventually leads to paralysis and death. Only ∼10-13% of ALS cases have an identified genetic basis^4^, but 40-60% of the risk of ALS is believed to be inheritable^5^. Genes whose mutation causes ALS include *SOD*1^6–8^, which encodes superoxide dismutase 1, and *TARDBP*^9–11^, which encodes the RNA/DNA-binding protein TDP-43^12–14^. Another gene associated with ALS, *C9ORF72*^15–17^, contains a hexanucleotide repeat whose expansion leads to the production of toxic RNA and/or protein products. The remaining cases are known as sporadic ALS (sALS) and have, so far, no known genetic cause.

Except for cases linked to *SOD1* mutations, all ALS cases (including sporadic and those with *C9OR72* mutations^15, 18–20^) display TDP-43 pathology^21–23^. TDP-43 pathology also occurs in other neurodegenerative diseases, in particular frontotemporal dementia (FTD), but less consistently so. For instance FTD-linked mutations in *C9ORF72* and *GRN* (which encodes progranulin) reliably lead to TDP-43 abnormalities^15, 18–20, 24–27^ whereas mutations in *MAPT* (which encodes tau) do not^28, 29^. At the molecular level, TDP-43 pathology is characterized by a loss of nuclear TDP-43 accumulation^30–33^ and the build-up of cytoplasmic TDP-43^30^ that can be hyperphosphorylated^34–38^, ubiquitinated^21, 33, 39^ and aberrantly cleaved^40–44^, all of which result in downstream molecular events such as activation of cryptic exon splicing (CES)^45–49^, dysregulation of gene expression and protein aggregation^50^. The fact that mutations in TDP-43 itself can directly cause ALS^12, 51, 52^ (and occasionally FTD^21, 22, 53^), demonstrates that TDP-43 dysfunction can directly drive neurodegeneration. Yet in the majority of ALS cases, TDP-43 is not mutated^6, 9, 12, 54^ leaving a fundamental unresolved question: is TDP-43 pathology a primary driver of neurodegeneration, or a downstream consequence of cellular stress? Further, why is TDP-43 pathology present in some disease contexts, but absent in others, and what does this reveal about underlying mechanisms? Given the well-established heterogeneity of ALS, both in terms of genetic origin and disease course, it is important to determine which cellular signatures are shared across ALS subtypes and which are unique to specific forms of the disease. Distinguishing between these possibilities is essential for defining the role of TDP-43 in ALS pathogenesis. We hypothesize that mutations in *TARDBP* mutations (collectively referred to as TDP-43^muts^) and *C9ORF72* (collectively referred to as C9orf72^muts^) confer distinct yet partially overlapping cellular mechanisms that differ from each other and from sporadic ALS. Our objective was therefore to define both the shared and mutation-specific cellular signatures underlying ALS pathogenesis.

Elucidating these signatures may provide insight into underlying disease mechanisms. Ultimately, our goal is to identify features shared across ALS subtypes, as these may enable the development of more broadly effective therapies than those targeting mutation-specific pathways. The recent success of the groundbreaking gene-based therapy, Toferson, that targets SOD1-mediated ALS^55, 56^, underscores that a deeper understanding of genetic drivers and underlying mechanisms is critical for developing effective ALS therapies. Further, if the disease mechanisms underlying ALS are heterogeneous as opposed to singular, then identifying the molecular signatures that define biologically distinct subtypes becomes essential.

To identify shared or subtype-specific cellular signatures of ALS, we imaged live cells from ALS patients differentiated into motor neurons (iMNs). Live cell imaging captures morphology on a cell-by-cell basis which reveals as an integrated readout of the cell’s molecular state. Each pixel, together with its spatial context, can encode information arising from interacting genomic, epigenomic, proteomic, and metabolic processes. In fact, numerous studies have demonstrated that cellular images contain rich information not only within individual pixels, but also in the spatial relationships among them that can reflect emergent properties of the cell and its underlying molecular landscape^57–60^. Moreover, we have shown that images of unlabeled cells contain sufficient information to accurately predict fluorescent labels of cellular structures, as well as cell type and state, underscoring that these images encode far more information than is accessible to the human eye^61^. To image live cells, we use a live-cell imaging platform called robotic microscopy (RM). RM collects epifluorescence images from individual neurons in a high-throughput fashion over the life of the cells^62–69^. We have used RM extensively in the context of ALS. For example, RM has allowed us to elucidate TDP-43 disease mechanisms in ALS^70^, investigate cell non-autonomous roles of human glia in neurodegeneration in fALS^71^, discover genetic modifiers of neurodegeneration in models of ALS^72, 73^, develop small molecule autophagy inducers that mitigate disease phenotypes^74, 75^ and develop new artificial intelligence analyses^61^.

We then subjected images of patient-derived and gene-edited iMNs to machine learning (ML) approaches, which can uniquely harness this high-dimensional information to classify ALS and its subtypes with superhuman precision. ML has an extraordinary ability to detect complex patterns and signals that are difficult or impossible for human observers to discern. ML has been applied in diagnostics in AD^76, 77^, stratification for various dementias^78^ or elucidating disease progression in ALS, PD and other neurodegenerative diseases^79–81^ using clinical and image data or both. For example, we have used convolutional neural networks (CNNs) to correlate amyloid β pathology in human brain samples with AD clinical severity^82^.

However, ML has not been widely applied to human iPSC-derived neuronal models to uncover disease-specific signatures or to elucidate underlying mechanisms. Notably, a recent study employed DNNs to classify control and VCP-mutant motor neuron lines using images from fixed and stained cells, highlighting the promise of this ML-based approach for disease phenotyping^83^. However, classification based purely on live human motor neuron morphology has not, to our knowledge, been reported. We developed classifiers capable of distinguishing images of ***live*** iMNs carrying TDP-43^muts^ from images of normal controls and to identify the cellular features they used to make this distinction. We then asked whether these classifiers could also distinguish iMNs carrying C9orf72 mutations or derived from sALS patients from control iMNs.

Here, we present classifiers spanning DNNs and shallow connected machine learning networks (SML) models that discriminate, with moderately high accuracy, TDP-43^mut^ iMNs from control cells, as well as moderately classifying ALS from patients with expansions in the *C9ORF72* gene or from patients with sporadic sALS. We also implemented a novel feature-based model we previously developed that allows us to capture various morphological properties of live cells in a temporal fashion^84^. While cross-sectional classifiers can detect disease-associated morphology in ALS, the disease is a rapidly progressive disorder. Therefore, we examined if modeling longitudinal transitions would improve our models’ ability to capture disease-associated phenotypes. Because we generated images of cells with different genetic backgrounds of ALS and tracked the same groups of cells over time, we captured not only rich spatial information within each image, but also additional information in the temporal domain which provides us with an extra axis along which differences can be resolved. Using linear mixed-effects models to estimate time-by-disease interactions, we quantified a comprehensive panel of size, shape, complexity, texture, and brightness features. We found evidence of alterations that may reflect developmental changes, which is a relatively overlooked aspect of the disease until recently^85, 86^. This analysis revealed coherent trajectories that were shared across ALS groups or specific to particular subtypes. Our findings suggest that ML models may be capable of identifying unique patterns associated with different underlying mechanisms of ALS. This approach may offer a novel and innovative avenue for elucidating disease mechanisms, particularly in the context of complex disease disorders.

## Methods

### iPSC acquisition and culturing

Control, sALS and C9orf72 iPSCs were obtained from Answer ALS^87^ (**Supplementary Table 1**). We acquired a patient-derived line carrying the *TARDBP* A382T mutation (TDP-43^A382T^) (NN0005320/ NH50306)) and its gene-corrected control (GC-CTRL^1^) (NN0005319/ NH50305) from the NINDS Coriell collection^88^. In addition, we generated (via gene editing) lines containing TDP-43^M337V^ and TDP-43^Q331K^ mutations in the background of a healthy control from the Personal Genome Project Canada (PGPC)^89^ (PGP-CTRL^1,2^) to create an isogenic series (**Supplementary Table 1 and Figure 1**).

**Figure 1.**
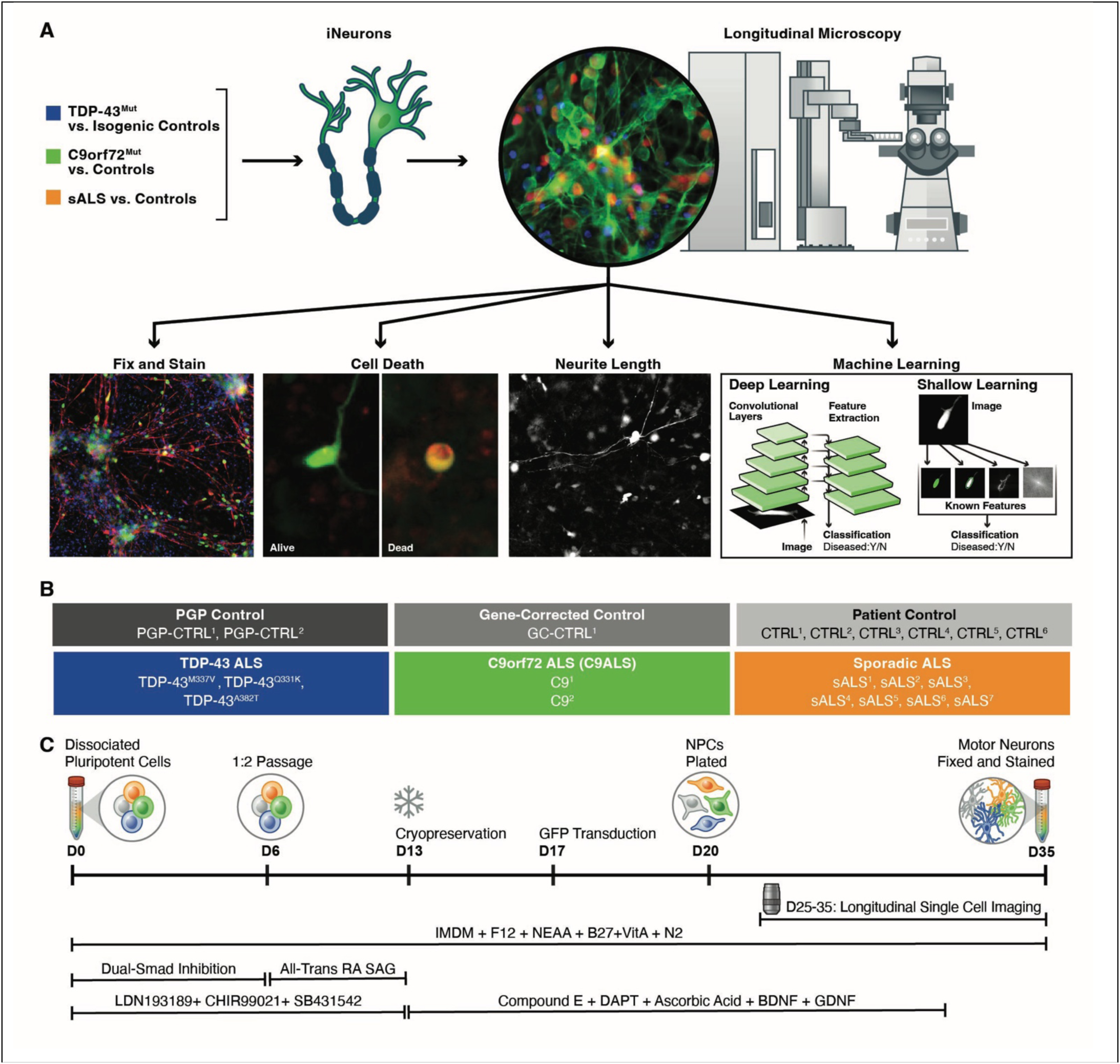
Overview of deep phenotyping and high-content analyses of ALS iMNs compared to controls. **A**) Workflow and platform for capturing TDP-43^muts^, C9orf72^muts^, sALS and control cellular signatures using several phenotyping metrics and both shallow and deep learning. **B**) Lines used in the study **C**) Motor neuron differentiation paradigm and timeline.

### CRISPR gene editing

The parental PGP1 line was engineered by Synthego Corporation (Redwood City, CA), to introduce M337V (*c*.1009A>G) or Q331K(*c*.991C>A) mutations in the coding sequence of *TARDBP* gene. Editing was achieved using *Sp*Cas9 nuclease and sgRNAs targeting exon 6 of *TARDBP*, co-delivered with single-stranded DNA template. SgRNAs were selected based on high on-target activity and minimal predicted off-target effects. Following transfection and clonal expansion, genomic DNA was extracted and the *TARDBP* locus was amplified via PCR using the following primers: Forward primer (5′–3′): GCGCAGTCTCTTTGTGGAGA, Reverse primer (5′–3′): GCTGCACCAGAATTAGAGCC. The introduction of intended SNPs was confirmed through Sanger Sequencing.

### Genotyping and karyotyping of iPSCs

We evaluated the TDP-43^M337V^, TDP-43^Q331K^, TDP-43^A382T^ lines (collectively TDP-43^muts^), two controls (PGP-CTRL^1-2^) and one gene-corrected control GC-CTRL^1^ (collectively CTRLs^All^). Cell pellets were sent to Laragen^90^ for Sanger sequencing to evaluate the presence of the point mutations introduced by CRISPR gene editing or in the parental lines. All variants were confirmed as follows: TDP-43^M337V^ contained the M337V:c.1009A>G mutation, TDP-43^Q331K^ the Q331Kc.991C>A variant, and TDP-43^A382T^ the A382T:c.1144G>A mutation. All controls (PGP-CTRL^1-2^ and GC-CTRL^1^) were confirmed to be WT at these locations. iPSCs from AALS were previously evaluated using whole-genome sequence data (WGS)^87^. While variants in a subset of ALS genes were found in sporadic ALS (sALS) or control lines, no variants known to be associated with ALS were found in the sALS lines^87^ (**data not shown**). Repeat expansions at the *C9ORF72* locus were evaluated in all AALS lines using the ExpansionHunter tool^91^ by the New York Genome Center^87^ or by non-CLIAA certified repeat-primed PCR data as previously described ^92^. C9^1^ and C9^2^ (collectively C9orf72^muts^) displayed expansions in the *C9ORF72* gene (**Table 1**).

We also examined the karyotype of all lines in this study (**Figure 1 and Supplementary Table 1)**. Frozen cell pellets were sent to Cell Line Genetics^93^ and were subjected to high-resolution array comparative genomic hybridization (aCGH array) using an Agilent 180K Standard aCGH +SNP. No gross abnormalities were found across all sALS, C9orf72^muts^, TDP-43mut engineered and gene-edited lines. However, the PGP-CTRL^1^, TDP-43^M337V^ and TDP-43^Q331K^ lines contained a small amplification (∼670 KB) of the long arm of chromosome 20, which is commonly observed in iPSCs^94^.

### iPSC culturing and motor neuron differentiation

All iPSCs were thawed into media with ROCK inhibitor (ROCKi; Selleck Chemicals, #S1049) and mTESR Plus media (StemCell Technologies, # 00-1130). iPSCs were cultured in mTESR Plus and grown on Growth Factor Reduced Matrigel GFR (Corning #356231). iPSCs were passage ∼ 1:5 using ReLeSR™ or Versene (Thermo Fisher Scientific, #15040066). All lines were expanded, banked and stored in LN2. To start the motor neuron differentiation, iPSCs were grown to ∼90% confluency, washed once with DBPS 1X (Thermo Fisher, #21-031-CV) and disassociated with Accutase for ∼10 minutes (Thermo Fisher #NC9839010). iPSCs were washed in mTESR and plated at 4 million cells per T25 flask (Omnilab, #5430429) in mTESR plus ROCKi.

When the iPSCs reached ∼90% confluency (usually within 1-2 days after plating), they were cultured in Stage 1 media similar to previous reports^87^. Stage 1 media consists of the following: Base media, corresponding to 1X IMDM (Gibco, #12440061), 1X F12 (Gibco, #11765062), 50X NEAA (Gibco, #11140-50), 100X N2 Supplement (Life Technologies, #17502048), and 50X B27+VitA (Gibco, #7504044), in addition to 10 mM LDN 193189 (R@D/ Tocris, #6053), 10 mM CHIR 99021(R@D/ Tocris, #4423), and 10 mM SB 431542 (R@D/ Tocris, #1614). The switch to Stage 1 media corresponds to day 0, during which the cells are fed daily with 10 mL of Stage 1 media until Day 6. On day 5, T-25 flasks were coated with 2.5 mL of a 1:4 dilution of 100 µg/mL poly-L-ornithine solution (Sigma, #RNBK5888) to 25 µg/mL using sterile cell culture water and left at room temperature overnight. The next day, the flasks were washed thrice with cold cell culture water and 3 mL of a 1:50 dilution of mouse laminin (Sigma, #L2020) diluted in cell culture water was added. The flasks were then stored in the incubator at 37°C for at least 30 min. Day 6 corresponds to the switch from Stage 1 to Stage 2 media, whereby the iPSCs are being patterned to motor neuron progenitors (MNPCs). The MNPCs were split 1:2 by use of a cell scraper (Perkin Elmer, #6057500). The MNPCs were pipetted up and down thrice using a 10 mL pipette before being distributed into the new flasks and cultured in Stage 2 media. Stage 2 media consisted of Stage 1 media plus the addition of 10 mM All Trans Retinoic Acid (AT-RA) (Stemgent, #04-0021) and 100 mM SAG (R@D/ Tocris, #4366). During this passage, Stage 1 media is replaced by Stage 2 in addition to ROCK inhibitor. On day 7, the media is switched to only Stage 2, fed every other day until day 13, at which point the MNPCs were cryopreserved. To freeze, the MNPCs were washed with DBPS and disassociated with Accutase for 20 minutes at room temperature before washing with Stage 2 media with ROCK inhibitor. The MNPCs were frozen at 8 million cells per cryovial in freezing media, which consists of 4 µg of FGF (R@D, #233-FB) for every 1 mL of Stem-Cell Banker GMP Grade (CedarLane Labs, #11890).

The day prior to thawing the cryopreserved MNPCs, a 12-well plate was coated with 1 mL of a 1:4 dilution of poly-l-ornithine solution to cell culture water and left at room temperature overnight. The day of thaw (which corresponds to day 13 of patterning), the wells were washed thrice with cold cell culture water and plated with 1 mL of a 1:50 dilution of mouse laminin to cell culture water and left in the incubator at 37°C for at least 30 minutes. The frozen MNPCs were thawed in the water bath until a small pellet is visible before being transferred to Stage 3 media with ROCK inhibitor. Stage 3 media^87^ consists of base media with the addition of 10 mM AT-RA, 100 mM SAG, 10 mM db-CAMP (Millipore/ Sigma, #D0627), 1 mM Compound E (Millipore/Sigma, #565790), 10 mM DAPT (Tocris/ R@D, #2634), 500 µg/mL Ascorbic Acid (Sigma, #A4403), 25 µg/mL BDNF (R@D, #248-BD), and 10 µg/mL GDNF (R@D, #212-GD). MNPCs were plated at 2.34 million cells per well. The media was changed the next day to only Stage 3, and the MNPCs were fed every other day for 6 days, until Day 20. On ∼day 17, the cells were transduced with the lentivirus containing the GEDI^95^ LV-Synapsin-GFP (Signagen # SL100271) or the 2Gi2R^96^ biosensor on at an MOI of 5–10 and washed the next day.

On differentiation day 18, plates were prepared for the final passage. CellCarrier-384 Ultra PDL-coated Microplates (Perkin Elmer, #6057500) were coated with 40 µL of a 1:4 dilution of 100 µg/mL poly-L-ornithine solution to 25 µg/mL using sterile grade cell culture water and left overnight at room temperature. The following day, the plates were washed thrice with cold cell culture water and 40 µL of a solution consisting of 5mg/mL human fibronectin (Corning, #CB40008A) and 1 mg/mL mouse laminin to 5 µg/mL with the use of cold sterile grade cell culture water. The plates were stored overnight in the incubator at 37°C. On day 20, the cells begin to mature into neurons and we call these MoNOs. The day-20 MoNOs were disassociated with Accutase for 20 minutes at room temperature and washed with Stage 3 media with ROCK inhibitor. The cells were further dissociated with pipetting several times and pushed through a 40 µM cell strainer lodged inside a 50 ml falcon tube three times. The MoNOs were then plated at a density of 30,000 cells per well, and the media was changed twice a week. After day 25 they are called motor neurons (iMNs). iMNs and were subjected to RM from day ∼27 to 38 until fixation for immunocytochemistry, which is typically on differentiation day ∼40.

### Immunocytochemistry

iMNs were fixed at day ∼40 with 4% PFA + 4% sucrose at 30 µL per well for 15 minutes at room temperature. The cells were then washed 1x with PBS at 60 µL per well before adding 80 µL of PBS and stored at 4°C. The iMNs were then permeabilized with PBS + 0.1% Triton-X at 80 µL per well and incubated at room temperature for 20 minutes. Following this, the cells were treated with 1 M Glycine in PBS for 20 minutes at room temperature at 80 µL per well. Block solution, consisting of 0.1% Triton-X and PBS + 2% FBS + 3% BSA was added for 1-2 hours at room temperature. After blocking, primary antibodies were added at 50 µL per well at different dilutions in the block solution as listed in the **Supplementary Table 2**, and the cells were incubated at 4°C overnight. The following day, the iMNs were washed thrice with PBS + 0.1% Triton-X (5 minutes/wash). Secondary antibodies were then added at 50 µL per well. Prior to being added, the antibodies were spun at 800 RPM for 5 minutes and diluted in the block solution. The cells were incubated for 2 hours at room temperature and covered in foil. Following the incubation, the cells were washed once with PBS for 10 minutes before washing once more with PBS and DAPI at a 1:1000 dilution for 5 minutes at 80 µL per well. The plate was imaged after washing once more with PBS.

### Quantification of stained images

CellProfiler was used to quantify differentiation propensity across all the iMNs. After fixing and staining with various antibodies as described above, cultures were imaged using RM. Images were subjected to a modified CellProfiler98 pipeline as previously described^84, 97^ and adapted to measure the percentage of cells stained with specific antibodies. Following the approach detailed in the CellProfiler examples (https://cellprofiler.org/examples), images were pooled and analyzed on a per-image basis. Images were first subjected to illumination correction using the Fit Polynomial smoothing method and the divide illumination function to ensure homogeneous brightness and illumination across all images. Following the corrections, nuclei objects were identified and segmented from DAPI images using a diameter minimum and maximum for size and the minimum cross-entropy intensity thresholding method^98^ to capture nuclear objects within the desired size range. This was followed by a feature enhancement step that increases the signal-to-noise ratio. These enhanced images were then segmented to identify all objects meeting the criteria for size and signal intensity. Intensity values for each object were analyzed to exclude non-nuclear artifacts or dead cells with abnormally bright nuclei, and valid objects were relabelled as nuclear segments.

After the nuclei object identification step, images were subjected to speckle feature enhancement, which improved signal clarity, before object identification similar to the method mentioned above. The “RelateObjects” module links nuclei and antibody objects by assigning which antibody signals are associated with each nucleus, allowing for the number of antibody objects per nucleus to be counted. The resulting objects were designated as “positively labelled” segments. These segments were then correlated with the nuclear segments to determine the proportion of nuclei expressing the respective antibodies. We calculated the fraction of positively labelled cells by dividing the total number of positively labelled cells with the total number of nuclei. Segmentation overlays and numerical results were exported and plotted using Prism10^99^ for visualization.

### Neurite length analysis

Neurite length and branching for all lines was evaluated by quantifying the length and branches of neurons as previously described^84, 97, 100^. Briefly, image tiles in the GFP channel of iMNs transduced with GEDI were picked at differentiation day ∼31 (Image acquisition timepoint four “T4”) to run through a CellProfiler pipeline identifying neurites. First, neuronal somas are identified as a primary object and then refined to include clear, bright objects outside of any clumps. Clumped objects are eliminated due to the high probability of overdrawn neurites from overlapping processes. Next, secondary objects are identified as projections coming from the seed object, the soma. Projections are assigned to a seed object and merged in the next step to create a single object. Neurite trace markers are identified as trunks, branches, non-trunk branches, and total skeleton length. These values are plotted using GraphPad Prism version 10.5.0 (673) for Mac, GraphPad Software, Boston, Massachusetts USA^101^.

### Longitudinal cell imaging and analysis

iMNs containing biosensors were imaged longitudinally via robotic microscopy (RM), which has been extensively described elsewhere^62–69^. iMNs were imaged every day for ∼8–10 days as previously described^62–69, 87^. The plates were kept in an incubator at 37°C during imaging (Liconic Automated STX44-ICBT-CO2 incubator). Plates were loaded from the incubator with a plate loading robot (PreciseFlex PF400), which is a robotic arm capable of transporting plates to and from the microscope from the incubator. The plates were imaged on a Molecular Devices Image Express Micro Confocal (IXM) system using a 20X objective. Images were collected over the course of the experiment and to identify and track individual iMNs. Images of different microscope fields from the same well were stitched together into montages, and montages of the same well collected at different time points were organized into composite files in temporal order in a computational pipeline constructed within the open-source program Galaxy. ImageXpress Micro Confocal filter sets included DAPI (excitation: 377/54 nm, emission: 447/60 nm, dichroic: 409 nm), FITC (excitation: 475/34 nm, emission: 536/40 nm, dichroic: 506 nm), Cy3 (TRITC) (excitation: 531/40 nm or 543/22 nm, emission: 593/40 nm, dichroic: 562 nm), Texas Red (excitation: 560/32 nm, emission: 624/40 nm, dichroic: 593 nm), and Cy5 (excitation: 631/28 nm, emission: 692/40 nm, dichroic: 660 nm). Images acquired were background-subtracted and stitched together by our custom-built image processing pipeline built in Galaxy as previously described^84, 87^.

### Cumulative risk of death using COX proportional hazards models

iMNs were transduced with the Synapsin:EGP biosensor and imaged daily using RM as previously described^62–69, 84, 87, 100, 102^ on an ImageXpress Micro Confocal High-Content Imaging System from Molecular Devices for 7-9 days. Images of different microscope fields from the same well were stitched together into montages using a custom-built image processing pipeline in Galaxy^84, 87^, and montages of the same well collected at different time points were organized into composite files in temporal order. Images were hand curated for survival using a custom Matlab script previously described^68, 74, 84, 87, 100, 103^ that records the last timepoint a human curator finds the neuron alive. The live dead status for each neuron was evaluated using Kaplan-Meier (KM) analysis and Cox Proportional Hazards analysis were calculated using custom scripts written in R, and survival functions were fit to these curves to derive cumulative survival and risk-of-death curves that describe the instantaneous risk of death for individual neurons as previously described^68, 74, 84, 87, 100, 103^. We performed survival analysis by tracking single cells over time and using the Cox Mixed Effects model (COX ME)^104^ to estimate the cumulative risk of death as previously described^87^. The COX ME captures experiment-to-experiment variability and the image-to-image variability within each experiment; the individual cell lines themselves were modelled as random effects. The hazard ratio of neuron survival of disease lines versus control lines was estimated as a fixed effect. The design of the experiment was such that in no situation was the experiment effect entirely confounded by the cell-line effect.

### Analysis of GEDI and calculating the odds ratio of cell death

GEDI^95^-expressing control, TDP-43^muts^, C9orf72^muts^ and sALS iMNs were imaged using RM over a 7–10-day period. Image segmentation was performed with a custom-built pipeline on the Galaxy platform^105^. The GEDI ratio for each cell was calculated, and thresholds distinguishing “alive” from “dead” cells were determined as previously described^95, 106^. iMNs from each cell line were assayed for viability across multiple plates, experiments, and time points.

The change in odds ratio of cell death at any given time (OR-CD) between cell lines was estimated using a Generalized Linear Mixed Effects Model (GLMM)^107^ or a Linear Mixed Effects Model (LMM)^108^ implemented via the glm function in R^109^ with a binomial family. Model covariates included plate (random effect), time (continuous), cell line (fixed), and the interaction between cell line and time^97, 106, 110^. The coefficient β represents the additional change in the log odds of cell death among ALS cells from baseline (0 hr) to a given time point, relative to the corresponding change in control cells. The odds ratio (OR) thus quantifies the odds of cell death in sALS, TDP-43^muts^ and C9orf72^muts^ iMNs at a given time relative to control cells. Statistical significance was defined as p < 0.05 under the null hypothesis β = 0, using the glmer function in R^109^.

The final model incorporated experiment ID as a random effect, and fixed effects for cell line, time, and their interaction. Corresponding odds ratios were derived from model fits. A similar GLMM was applied to compare disease lines (e.g., multiple ALS lines) against controls, with cell line and experiment ID as random effects, and disease class (0 = control, 1 = disease), time, and their interaction as fixed effects. Time was modeled either as a continuous variable (linear model) or categorical variable (non-linear model). Model selection was based on the lowest Akaike Information Criterion (AIC)^111^. Non-linear models best described TDP-43^muts^, C9orf72^muts^, sALS versus control comparisons. All analysis code is available on GitHub^112^.

### Image processing and feature extraction

6144 x 6144 montages consisting of 3 x 3 well panels from control, sALS, C9orf72^muts^ and TDP-43^muts^ iMNs expressing the GFP from the GEDI biosensor^95^ were obtained by RM and processed with our custom-built image-processing pipeline using Galaxy software^113^. In Galaxy, the panels first went under background subtraction to obtain a high signal-to-noise ratio. This involves the generation of a background image that is the median of all panels in each well. Montages are generated by stitching these panels together. The montages are subjected to a custom-built pipeline written in Python^114^ called “CropCellPipeline” (available at https://github.com/naamirani/IXMCropPipeline) that takes as input the montages of each well at image-acquisition timepoints 1 and 6 (differentiation days ∼27 and 33 respectively for the gene-edited control and TDP-43^muts^ images, and days 25 and 31 for images of other controls and of sALS and C9orf72^muts^).

The CropCellPipeline has different modules that perform different functions. The S1 module generates crops (115 x 115 pixels) that capture a single cell’s body and branching neurites. Image crops were filtered so that the objects (iMNs) contained a minimum eccentricity value of 0.3, and an area size of 250–10,000 pixels, and mean intensity above 80. To ensure that each soma was not truncated, we checked if the contour of the soma could be detected on the borders of the crop via pixel coordinates by the S2 module. The cell crops are then sorted by disease or control categories, as determined by the origin of the cell line, in the S3 module and subjected to feature calculation algorithms described below as part of the S4 module.

### Model training

We trained both SML and DNNs on the same exact set of images (**Supplementary Table 3**). First, each model was fit exclusively on the training set, with internal cross-validation performed only within this subset to generate out-of-fold predictions for stacking. The validation set was held out during this stage and used solely for external performance assessment and hyperparameter tuning. After finalizing the model configuration, a full retraining step was performed using the combined train and validation sets to maximize data used for parameter estimation. Final performance was evaluated on an independent test set, via prediction accuracies on images based on the test filename lists that had never been seen by the model. As a negative control, class labels were randomly permuted within each data split (training, validation, and test) while keeping the feature matrices fixed. While inference rests primarily on the training/validation permutation (a model trained on scrambled labels would be expected to perform at chance regardless), labels were permuted across all splits for procedural consistency. This procedure preserves class balance, but removes the true correspondence between samples and labels. Models were trained and evaluated using the scrambled labels under otherwise identical conditions. In this experiment, a collapse of performance to chance levels supports the interpretation that classification performance in the primary analyses reflects meaningful signal in the cellular features rather than data leakage or spurious correlations.

### Shallow connected machine learning networks

The feature detection algorithms are written in Python and contained within Jupyter notebooks^115^. We used various feature detection algorithms using the image processing packages OpenCV ^116^, Pillow^117^, and Scikit-Image^118^. We applied a hybridization of Otsu and binary thresholding^119^ as a preprocessing step for multiple features to isolate the cell for feature calculation. The features that were used were: contour perimeter^120^, contour area^120^, contour eccentricity with ellipse^121^, solidity^122^, radial distance^123^, local peak maxima^124^, luminance gradient^125^, Fast Fourier Transform Shift Magnitude Median (FFT-MED)^126^, Inverse FFT Shift Magnitude (I-FFT-MEA)^126^, HOG feature^127^, Minkowski-Bouligand Fractal Dimension Score^128^ with Sobel transformation^129^ applied in both y- and x-axis directions, Sholl Intersections Analysis^130^ with Skeletonization transformation^131^, Voronoi Points^132^ evaluated as Clusters with DBSCAN^133^ evaluation (VP-C), Shi-Tomasi^134^ + FAST^135^ weighted corner detection, and Gray-Level Co-occurrence Matrix Homogeneity^136^.

These features capture different properties from each cell. The shape and size of the soma were measured by edge detection, which uses canny processes and simple tree approximation. From this we calculated the contour perimeter^120^, contour area^120^, contour eccentricity with ellipse^121^ represented by the largest elliptical contour. Solidity^122^ was obtained by drawing the convex hull area of the soma and dividing by contour area. The radial distance^123^ feature captures the largest object moment in the image, which in our images is the soma. The median distance from the centroid to each of the four extrema (upper, lower, right, left) was calculated. We also used the Voronoi algorithm^132^ to generate points of interest of the cell as divided by specific Voronoi regions. We then applied the DBSCAN^133^ algorithm which dynamically calculates the optimum number of clusters from these points, as represented by VP-C.

For capturing complexity, Minkowski-Bouligand Fractal Dimension Score^128^ is a box-counting fractal dimension that measures how complexity of detail changes with the scale at which one views the fractal dimension of a set S in a Euclidean space Rⁿ. Shi-Tomasi + Fast Detection is similar to Harris^137^ corner detector except for the way the score (R) is calculated, which yields a better result especially if combined as a weighted average with FAST calculated corner counts. Gray-Level Co-occurrence Matrix (GLCM)^136^ Homogeneity measures the closeness of the distribution of elements in the GLCM to the diagonal of the GLCM. The GLCM itself characterizes image texture by counting how often pixels with a certain intensity value occur in a specific spatial relationship to pixels with other intensity values^138^. Fast Fourier Transform converts spatial or temporal data into the frequency domain data for exploring the power spectrum of a signal. Of interest for our features is FFT-MED, the median of magnitude spectrum, which indicates the central frequency content, and I-FFT-MEA, the mean of magnitude spectrum skew for inverse FFT shifted image. This translates to the overall brightness of the reconstructed image and therefore together provide an assessment of the dominant frequencies within each cropped image. Sholl Intersections Analysis^130^ is conducted by first generating a skeletonized mask of the cell, then counting the total number of intersections from the Sholl algorithm.

The HOG feature extracts the gradient, orientation/magnitude, and direction of the edges. The image is visualized as a grid and from there broken down into smaller regions, and for each region the means of the gradients and orientation are calculated. A histogram for each of these regions is generated separately and the overall slope mean was obtained. To evaluate brightness, we converted the image to the HSV format and extract the ‘V’ channel as the luminance value. Our images are grayscale, therefore the conversion to HSV is not strictly necessary, however the code is designed to also be usable by RGB images. Then, we compute the image gradients using Sobel filters, combine them into a gradient magnitude map, and then collect only the gradient values within the masked cell. This allowed us to measure how quickly brightness changes across the image. Evaluating brightness in this way allows the analysis to be more robust to uneven illumination or non-biological variations in brightness across cells.

Each object crop from their respective disease or control category was assigned a numerical score for each feature. These scores were output into a CSV file for further downstream analysis, after undergoing preprocessing for blank, non-numeric, or duplicate entries. Single-timepoint data represents cells as separated by day of imaging, while multiple-timepoint data concatenates both days.

In the S5 module of CropCellPipeline, we constructed training, validation, and test split file lists from the final datasets for TDP-43^mutatnt^, sALS, and C9orf72^mutatnt^ versus controls (**Supplementary Table 3**) according to a fixed 80/10/10 ratio to create training sets for fitting the model, validation sets for tuning hyperparameters, and test sets for assessing the final model. Each split is strictly balanced by class label (1 for ALS type, 0 for controls) by down sampling the larger class to match the smaller, guaranteeing that there are equal numbers of class 0 and 1 sample per file list. To further ensure fairness, the data were first organized per cell line within each class, capped so no cell line contributed disproportionately, and then split, guaranteeing that each experiment was represented in all splits. This enabled a distribution of data that is not only equal among ALS type and control, but also across the underlying cell lines and experiments. By balancing the data across experiments and cell lines, we can perform experiment- and cell line–specific hold-out tests to evaluate how excluding individual experiments or cell lines affects model performance.

Once the data was processed as previously described in the S1-S5 modules of CropCellPipeline and generated the feature CSV files and their train/test/validation split file lists, we enter the S6 module for SML. The balanced dataset was obtained by cross-referencing the file lists with the feature CSV files, then underwent a standard scaling that removes the mean and scales to unit variance as the last step before being introduced to our shallow machine learning models.

The ML models we used for classification between controls and disease are: Logistic Regression^139^, Support Vector Machine (SVM)^140^, MLP Classifier^141^ with a single hidden layer, Random Forests^142^, and XGBoost^143^. The final model that we used to generate results is a Stacked Ensemble Classifier^144^ incorporating MLP, SVM, and XGBoost predictions as its base models, with a Logistic Regression meta-model as the final estimator. In this Stacked Ensemble Classifier, the base models contribute predictions that are used as inputs to the meta-model. MLP captures importance through the average absolute weights connecting input features to the first hidden layer, representing how strongly each feature influences the network’s internal representation. SVM (with a linear kernel) provides direct coefficients for each feature in the decision function, which can be interpreted as the directional influence (positive or negative) of each feature on the decision boundary. XGBoost, which is a gradient-boosted tree model, reports feature importance using gain, which quantifies the average improvement in model performance when a feature is used for splitting, therefore offering a measure of true contribution to the model. All importance values were normalized from [0,1] to enable fair comparison across models. The outputs/predictions of these base models were used as inputs to a logistic regression meta-model, which learns how to optimally weight each base model’s prediction when producing the final classification. The logistic regression coefficients thus reflect the influence of each base model, not individual features, on the final stacked prediction.

### Convolutional neural nets using ResNet50

We employed SimCLR^145^, a contrastive self-supervised learning framework, to learn visual representations from the image crops described above for TDP-43^muts^, C9orf72^muts^, sALS versus controls in the exact same train/test/validation dataset splits for the SML models (**Supplementary Table 3**), described above such that we would be able to directly compare the results. The SimCLR model is designed to maximize agreement between differently augmented views of the same image in a latent embedding space and to learn the important features in the crop. The pretrained model uses a ResNet-50^146^ backbone. This pretrained model is then trained and saved to be used for further fine-tuning with labels. Each input crop was randomly augmented twice to create a positive pair. The augmentation pipeline included horizontal flipping, and Gaussian blur augmentations known to encourage the model to learn invariant representations. These augmentations ensure that the network learns robust features that generalize across different image transformations. The model was pre-trained using the normalized temperature-scaled cross-entropy loss (NT-Xent). For a batch of *N* crops, this creates 2*N* augmented views, and the loss is computed over all *N* positive pairs, with the remaining 2*N*−2 examples in the batch serving as negatives. The objective encourages the model to pull together embeddings of positive pairs while pushing apart all other negative pairs. The model was pretrained and subsequently fine-tuned using non-linear layers, with separate classifier heads for each group “TDP-43^muts^”, “C9orf72^muts^”, “sALS”) versus their respective controls. versus their controls. The model was tested on a hold-out dataset (**Supplementary Table 3**) that was not seen by the model at all during training.

Let *sim*(*u*, *v*) = (*u^T^v*)/( ||*u*|| ||*v*||) denoting the dot product between 2 normalized u and v (i.e., cosine similarity).

Then the NT-Xent loss function for a positive pair of examples (*i*, *j*) is defined as:

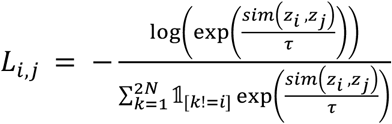

where 1_[*k!=i*]_∈ {0, 1} is an indicator function evaluating to 1 iff *k*! = *i* and τ denotes a temperature parameter. The final loss is computed across all positive pairs, both(*i*, *j*) and (*j*, *i*), in a mini-batch.

### Model Testing

After each model was trained on each group (TDP-43^muts^, C9orf72^muts^, sALS versus controls), we evaluated how well it performed on unseen data - hold-out data that was not used during training. This process, known as model testing, helped determine whether the model has truly learned the underlying patterns or has merely memorized the training examples. We compared the model’s predictions with the ground truth and compute a matrix that summarizes how many positive cases were correctly identified, how many were misclassified as negative, and vice versa. Then, we calculated the precision and recall metrics - precision indicates how many of the predicted positive cases are correct, while recall represents how many of the actual positive cases are successfully detected. Together, they reflect the model’s effectiveness in identifying true positives without producing excessive false negatives.

Train/test/validation image folders were then regenerated based on randomization of control and ALS images within each data split, such that amount of 0 and 1 classified images remained the same per split. These folders were then used to finetune the model, after pretraining with SimCLR as previously described^145^, and predict upon unseen data. The code for this pipeline can be found here: https://github.com/finkbeinerlab/ALS_classification_microscopic_data

### Convolutional neural nets using ResNet18

To classify TDP-43^mutant^ versus control iMNs using deep learning, we used ResNet 18d^147^ a variant of the ResNet18 architecture featuring a deeper stem with three 3×3 convolutions instead of a single 7×7 convolution, which improves feature extraction for small objects. The model was initialized with weights pre-trained on ImageNet^148^ and fine-tuned on our dataset to leverage transfer learning from natural images to cellular morphology.

Image crops (115 × 115 pixels) generated by CropCellPipeline were used as input. To match the expected input dimensions of ResNet18d, crops were resized to 224 × 224 pixels using bilinear interpolation. Single-channel grayscale images were converted to three-channel inputs by replicating the intensity values across all channels. Pixel intensities were normalized using ImageNet statistics (mean = [0.485, 0.456, 0.406], standard deviation = [0.229, 0.224, 0.225]) to maintain compatibility with the pre-trained weights.

Data augmentation was applied during training to improve generalization and reduce overfitting. The augmentation pipeline included random horizontal and vertical flips, random rotations (±15°), and random affine transformations with slight scaling (0.9–1.1×) and translation (±10% of image dimensions). These augmentations were applied stochastically to each image during training while validation and test images were processed without augmentation.

The network architecture retained the original ResNet18d backbone, consisting of an initial convolutional stem followed by four residual stages with [2, 2, 2, 2] blocks, respectively. Skip connections within each residual block enable gradient flow during backpropagation and facilitate learning of identity mappings. The final fully connected layer was replaced with a binary classification head outputting two classes (TDP-43^mutant^ versus control).

Training was performed using the same data splits as the SML models, with an 80/10/10 ratio for train/validation/test sets, stratified by class label and balanced across cell lines. The model was optimized using the AdamW optimizer with an initial learning rate of 1×10^−4^ and weight decay of 1×10^−2^. A cosine annealing learning rate schedule was applied over the training period. Cross-entropy loss was used as the objective function. Training was conducted for 50 epochs with early stopping based on validation loss to prevent overfitting, using a patience of 10 epochs. A batch size of 32 was used throughout training.

Model performance was evaluated on the held-out test set using area under the receiver operating characteristic curve (AUC-ROC), precision-recall curves, average precision (AP), and confusion matrices. As a negative control, labels were randomly permuted while preserving the feature matrices and class balance. Models trained on scrambled labels served to verify that classification performance reflected true biological signal rather than artifacts or data leakage. Separate models were trained for timepoints T1 and T6 to assess whether discriminative features emerged early or late in the imaging period. All code is available at: https://github.com/drewlinsley/genentech^149^ and: https://github.com/naamirani/IXMCropPipeline.

### Gradient-based saliency maps (SmoothGrad)

We measured model decision making strategies using SmoothGrad, a technique that yields a map showing the extent to which every image pixel contributed to the model’s ultimate decision. This is done by averaging gradients of the image decision with respect to multiple noise-perturbed inputs, to yield robust decision attribution maps. For each image, 50 noise samples were generated by adding Gaussian noise (σ = 0.1 × input dynamic range) to the FITC image and the resulting gradients of the target class logits with respect to the input pixels were computed, averaged together, then normalized to [0, 1] for visualization and quantitative comparison.

### Gradient–stain correlation analysis

To quantify the similarity between model-attributed features and biological markers, pixel-wise Spearman correlations were computed between each stain intensity map and the corresponding SmoothGrad saliency map. For each stain: Both stain intensity and saliency maps were flattened into vectors. Spearman’s ρ was computed across all image pixels. Correlations were averaged across biological replicates. Negative controls ensured that correlations were not driven by image background.

### AUC-Overlap (AUC-o) quantification

AUC-overlap was used to measure spatial co-localization between gradients and each stain. For each image, SmoothGrad maps and stain pixels were thresholded at 25%, 50%, and 75% intensities, then IOU was computed and ultimately integrated across these thresholds to produce an AUCHigher AUC-o indicates that gradient hotspots preferentially align with the brightest regions of a staining channel.

### Partial R² analysis and computation

To estimate the unique contribution of each stain to the variance explained in gradient distributions, partial R² values were computed using a multiple linear regression framework. For each image, the SmoothGrad map served as the dependent variable. Stain intensity channels (DAPI, NEUN, LaminB1, SUN2, GM130) were used as predictors. All images were spatially flattened before regression. For each stain: A full regression model including all stains was fit. A reduced model was generated by removing the stain of interest. Partial R² was computed as the change in explained variance between the full and reduced model: Partial R² values were averaged per sample and compared between control and TDP-43 mutant groups using a two-sided t-test with Welch’s correction. P-values were corrected for multiple comparisons using the Benjamini-Hochberg FDR procedure.

### Clustering and heatmap visualization

To visualize the structure of stain contributions across samples, Partial R² values were z-scored per stain. Hierarchical clustering was performed using Euclidean distance and Ward’s linkage. Heatmaps were generated using standardized color scales, with warmer colors indicating stronger unique contributions to the gradient signal. All statistical analyses were performed in Python using NumPy, SciPy, scikit-learn, and seaborn. Data are presented as mean ± SEM unless otherwise specified. A significance threshold of α = 0.05 was used.

### 2Gi2R biosensor analysis

Day 18 MNPCs were transduced with the nucleocytoplasmic shuttling biosensor 2GiR at an MOI of 2.5. The following day, the media was changed on the MNPCs and subjected to the differentiation protocol to generate iMNs as described above. 2Gi2R-containing iMNs were imaged in the RFP (150 ms) and GFP (150 ms) channels daily for 7 days starting on differentiation day 27. Montages of 9 tiles were imaged in RFP (150 ms) and GFP (150 ms) channels at 20x-40x magnification per well. Images were processed using a custom workflow in Galaxy^87^ that generated single cell crops from differentiation day 31. Cropped images were hand curated and filtered to eliminate crops that contain multiple cells, debris, or did not display neuronal morphology. The single cell crops from both the GFP and RFP channels were loaded into a custom-built Cell Profiler pipeline to identify RFP signal and GFP signal individually based on occupation of either fluorescent molecule. The GFP masks produced by Cell Profiler are refined by hand to identify signal from neuronal looking objects only. RFP masks are hand refined to eliminate objects too close together to eliminate any measurements from overlapping cells. Objects that have overlapping RFP and GFP masks are taken forward to calculate the area occupied by RFP to the area occupied by GFP on a per object basis. Object ratios were then plotted in Prism10.

### Time interaction analysis

We developed a custom-built pipeline to evaluate how cellular features change over time^84^. Images from iMNs from control, sALS, TDP-43^muts^ and C9orf72^muts^ iMNs were processed at both T1 and six days later at a later timepoint “T6” separately to generate feature CSV files. The feature CSV files are processed with the RMeDPower2^110^ tool kit that allows a user to examine the normality assumptions for the linear mixed model^150^ and adjust if necessary. We removed outliers via Rosner’s test^151^ which evaluates both ends of the data distribution. In addition to the features described for developing our SML model, we added two more features for RMedPower. The first is Variance, where we calculate the variance of the largest object/cell in the crop, and the second is Gaussian Distance Map^152^, where Gaussian blur is applied to the image, then a distance map is generated to detect and count objects in the image that remain after blurring. All of the feature values except for Sholl Total Intersections, Voronoi Points + DBSCAN Clusters are considered continuous features that are not normally distributed so they were log transformed as described^150^. This procedure is integrated into RMedPower2 and runs during data transformation to remove outliers on a per-feature basis. These outliers are marked as NaN and afterward, rows containing NaN will be removed from the transformed dataframe in preparation for downstream processing. Dataset and outlier information for both single-timepoint and multiple-timepoint data are shown in **Supplementary Table 3**. RMeDPower2^110^ for controlling for experiment and cell line as random effects are utilized to determine significance of difference in feature scores for “ALS” vs. “CTR” categories. We removed outliers and at initial timepoint T1 and the later timepoint T6 separately (single-timepoint analysis), as well as T1 and T6 in conjunction to investigate possible effects of time interaction (multi-timepoint analysis).

Once the data was preprocessed, the single-timepoint data is run through a two-class LMM in a custom-built R script^109^. All code is available on Github at: https://github.com/naamirani/IXMCropPipeline. LMMs^154^ are used to estimate associations of interest in situations where the responses are clustered or correlated by design. In our case, responses are typically clustered by the experiment and cell line in which these cellular responses are assayed. In LMMs the clustering variable is typically modeled as a random effect whose influence on the mean response is assumed to be drawn from a normal probability distribution with zero mean and given variance. This model outputs a table of associations including estimate, standard error, degrees of freedom, t-value, raw p-value, corrected p-value, Benjamini-Hochberg (BH) and Sidak SD multiple corrections methods, as well as effect size. For the purposes of remaining stringent with our research parameters, BH was selected as our reference metric, as it uses the p-values while also controlling for false discovery rate.

The multi-timepoint dataset is analyzed separately from the single-timepoint version because it explicitly accounts for the effect of timepoint differences. This LMM produces a timepoint-interaction CSV that includes the estimated effect, standard error, degrees of freedom, and corrected p-values for the comparison of sALS versus CTR as modulated by timepoint (denoted C1–T7 or C2–T7, respectively). A significant interaction indicates that changes in classification and changes in timepoint act synergistically to influence the corresponding phenotype. The direction of this effect can be inferred from the reported estimates.

To estimate this shift modulated by disease status we use a Linear Mixed Effects Model (LMM)^109, 155^ called RMedPower^150^ to capture the mean feature values in TDP-43^muts^, C9orf72^muts^ and sALS and control iMNs at each distinct timepoint and the additional shift across the two time-points due to a change in disease status via an interaction term. Hence, the interaction term is between the time and the disease status variables. The mathematical formulation of this LMM is described next and has been previously described^84^. Here we let *Y^t^_ijk_* denote the feature value in the natural or the logarithm scale of the *k^th^ (k ∈ {1,2,…,N^c^_ij_})* iMN measured at time-point *t* (*t* ∈ {*T*1, *T*7}), derived from the *j^th^* (*j* ∈ {1,2, ⋯, *N*^c^}) cell-line assayed in the *i^th^* (*i* ∈ *Exp* = {1,2, ⋯, *N*^e^}) experimental batch. *N*^&^ is the number of experimental batches. *N*^c^ is the number of cell-lines. *N^c^_ij_* is the number of iMNs from each line *j* assayed in experimental batch *i*. The notation *A* and *R* refers to the sets of all alternate and reference cell-lines respectively that are used in each experiment. The LMM states,

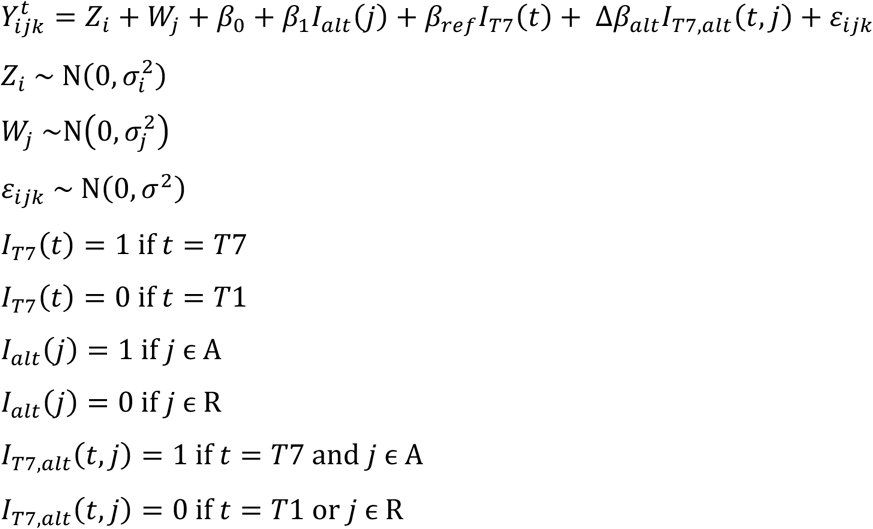

Where *Z_i_* denotes the random effect associated with the *i^th^* experimental batch, *W*_j_ denotes the random effect associated with the *j^th^* cell-line. Both these random effects are assumed to be drawn from normal distributions with zero means and variances σ^2^_i_ and σ^2^_j_ capturing the inter-experimental batch variances and inter-cell-line variances, respectively. The ε_ijk_ term captures the residual error of the model. Let β_ref_ refer to the change in the marginal expectation of the feature value over time for the reference cell-lines (e.g., CTRL), i.e.,

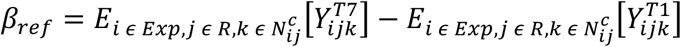

The β_0_ term captures the mean feature value at time T1 for the reference cell-lines, i.e.,

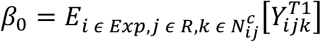

The β_1_ term captures the change in mean feature value at time T1 between the alternate and the reference cell-lines, i.e.,

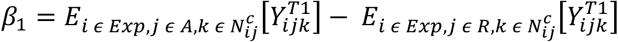

The subscripts for the expectation function, *E* indicate that the mean is computed by averaging out effects over all experimental batches, all reference cell-lines and all cells in each of these reference cell-lines. *I*_T7_(*t*) and *I*_T7,alt_(*t*, *j*) are indicator functions. The coefficient β_*alt*_ can be defined analogously for the alternate cell-line though is not directly estimated as one of the coefficients in the above model. Δβ_*alt*_ estimated by the coefficient of the interaction term in the LMM described above, captures the additional change over time of the mean feature value for the alternate cell-line. This interaction term corresponds to the “<u>M</u>orphology <u>C</u>hange <u>O</u>ver <u>T</u>ime” or “MCOT” (the time by disease status interaction) which is then scaled by the change over time in the TDP-43mut, C9orf72^mutant^ and sALS groups vs. controls expressed as percentages as previously described^84^. Specifically, MCOT is defined as,

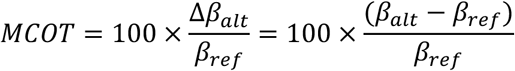

Significant differences in the rate of change of these features were identified by the significance of the interaction term. The LMMs were fitted using the lmerTest^109, 155^. Use of RM allows us to acquire very large sample sizes and therefore resulted in many observations and as such we observed very low p-values, but in some cases this change was very small therefore, we considered the effect sizes in highlighting significant associations^156^ and only considered a change of 8% or greater to be considered a *bona fide* difference across all groups.

## Results

### TDP-43 mutations do not impair the patterning of iPSCs into motor neurons

To identify molecular signatures of mutations in TDP-43 ALS, we performed deep phenotyping and high-content analyses of iMNs using robotic microscopy (RM) coupled with standard phenotyping assays and unbiased machine learning approaches (**Figure 1A**). We focused on three ALS-linked *TARDBP* variants A382T (TDP-43^A382T^), M337V (TDP-43^M337V^) and Q331K (TDP-43^Q331K^) which have been previously studied in the context of ALS. *TARDBP* variants disrupt liquid-liquid phase separation (LLPS)^157, 158^, and cause TDP-43’s misfolding, aggregation^159–161^ and toxicity^162^. These mutations all reside within the C-terminal glycine-rich domain of TDP-43 and, while their pathological features are overlapping but not identical, they likely converge on shared mechanisms^11, 156, 162–179^. Our goal was to find distinct TDP-43 signatures of pathology in lines that contain *TARDBP* mutations (TDP-43^muts^) compared to control lines (**Figure 1B, left and middle grey panels**). We obtained well-characterized iPSC lines from a healthy control from the Personal Genome Project Canada (PGPC)^89^ (PGP-CTRL^1^, PGP-CTRL^2^) (**Figure 1B, left grey panel**) and a patient-derived iPSC line containing the A382T mutation in *TARDBP* (TDP-43^A382T^) as well as its gene-corrected control (GC-CTRL) (**Figure 1B, middle grey panel**). We also created an isogenic series of TDP-43 variant-containing lines by introducing the Q331K and M337V mutations (TDP-43^Q331K^, TDP-43^M337V^) in the PGP-CTRL^1^ background via CRISPR editing (**Figure 1B, blue panel and Supplementary Table 1**).

To generate motor neurons (iMNs), we subjected the TDP-43^muts^ and control lines to a modified differentiation protocol (**Figure 1C**) based on previous reports^87, 106^. We fixed the cells after ∼5 weeks in culture (differentiation day ∼DIV35) and stained them with various neuronal and motor neuron markers to assess each line’s differentiation propensity. We observed robust MAP-2 staining indicating high numbers of mature neurons, robust expression of neurofilament H (NFH), and low numbers of proliferating cells as stained by KI67 (**Figure 2A and Supplementary Figures 1-4)**. We also observed typical expression of TDP-43 with no gross changes in localization of TDP-43 in the TDP-43^mut^ iMNs (**Supplementary Figure 5 and data not shown)**.

**Figure 2.**
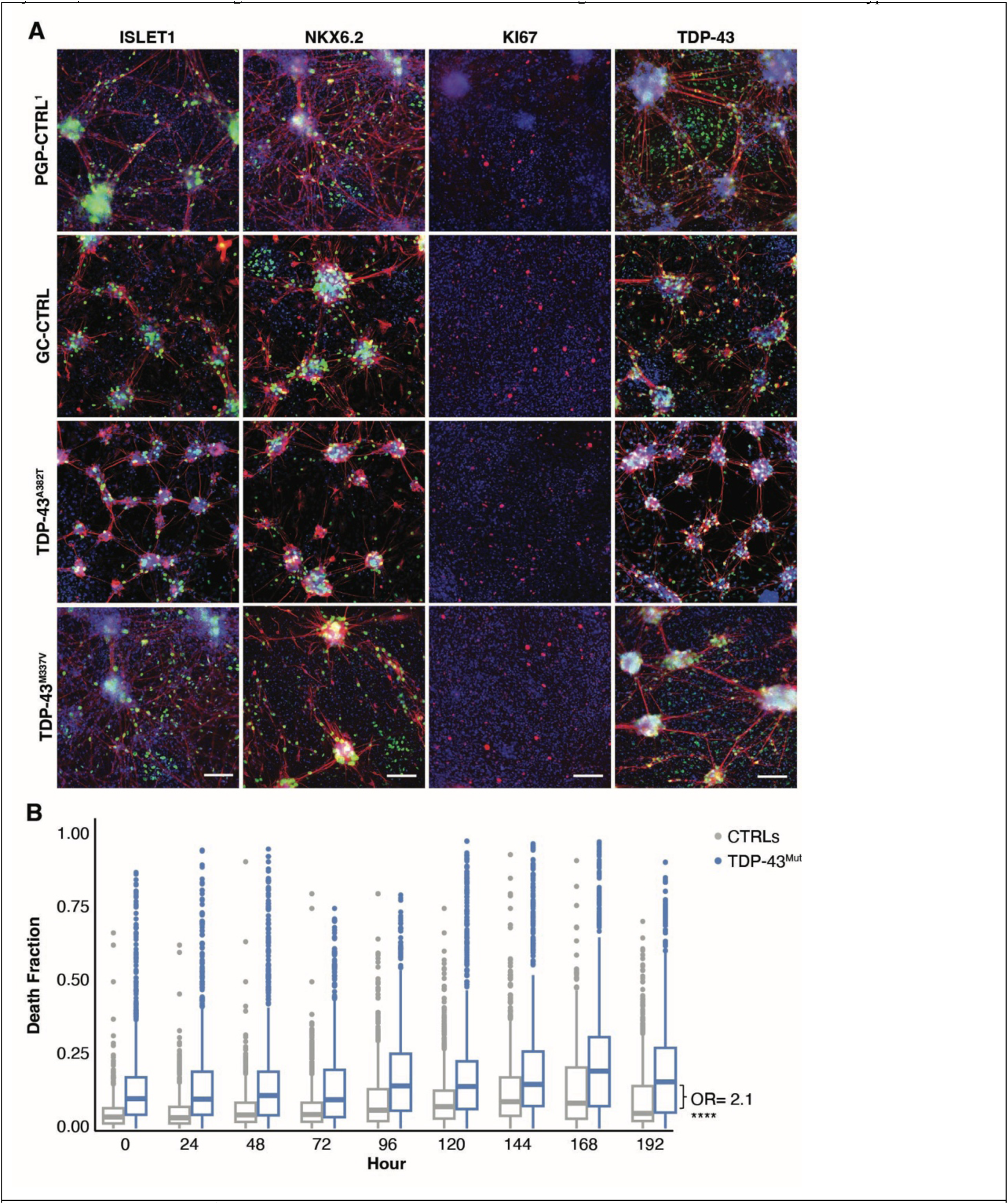
TDP-43^mut^ iMNs display typical patterning but degenerate more than controls. **A**) Representative images of iMNs used in the study. At differentiation day ∼DIV 35, iMNs were fixed and stained using antibodies against MAP-2 (all panels except KI67), ISLET1 (green, left), NKX6.2 (green, middle left), KI67(red, middle right), TDP-43 (right) and DAPI (blue-all) scale bar= ∼100 µm. **B**) Cultures of TDP-43^mut^ iMNs display a higher death fraction than cultures of healthy control iMNs, OR = 2.1, p < 0.00001. n = 9 experiments, live neurons = 1,447,764 cells, dead neurons = 198,904 cells. Boxplots display the median and interquartile range of the dead cell fraction per well per time point; whiskers represent data range. Individual dots show outlier wells for the TDP-43^mut^ (blue) and control (grey) iMN cultures and dots represent outliers by montage. Individual death plots are shown in **Supplementary** Figure 9. We evaluated the OR-CD at each time point using a generalized linear model (LM)108 with the glm function in R109.

The expression of motor neuron markers ISLET1 and NKX6.2 were moderate and variable across control and TDP-43^muts^ iMNs (**Supplementary Figure 6).** To determine whether this variability reflected an impact of the *TARDBP* mutations on motor neuron differentiation, we employed a linear mixed model (LLM)-based tool called RMeDPower^110^ that can account for variation across experiments, cell lines and batches. We found no significant differences in the proportion of ISLET1-positive or NKX6.2-positive cells between all controls and TDP-43^muts^, with an average of ∼33% ISLET positive and 38% NKX6.2-positive cells per image tile (**Supplementary Figure 6A, C)**. However, there were subtle differences across the gene-edited clones and gene-corrected lines compared to parental controls (**Supplementary Figure 6B, D)**. Similarly, we did not observe differences in the number of KI67-positive cells in the TDP-43^mut^ lines compared to the controls (**Supplementary Figure 6E, F).** We also did not detect significant differences in TDP-43–expressing cells (**Supplementary Figure 6 G, H)**. Together, this data suggests that overall, the cultures have patterned to a motor neuron-like fate with a low number of proliferating cells and display differentiation propensity similar to previous reports^87^.

### TDP-43 mutant iMNs degenerate in the absence of stressors

To examine the impact of *TARDBP* mutations on cell morphology and cell death, we transduced the iMNs with the RGEDI biosensor (RGEDI-P2a-EGFP)^95^, which combines an EGFP (green) marker of cell morphology with a Ca^++^-sensitive red fluorescent marker that only emits when cells are dying^95^. Combined with RM, RGEDI allows us to monitor tens to hundreds of thousands of cells at a time. iMNs were transduced with RGEDI on differentiation day ∼DIV20 and subjected ∼ 1 week later (∼DIV27) to daily time-lapse imaging using RM^62–69, 87, 102^ in both the red and green channels^95, 97^ for 9 days (**Supplementary Figure 7**). In the green channel, iMNs displayed typical neuronal morphology with dendrites and branching processes (**Supplementary Figure 8A**), and quantification of neuron length and structure on ∼DIV31 using previously described methods^100, 180^ revealed no significant differences in neurite length, number of processes or branch ends in the TDP-43^mut^ lines compared to their isogenic controls (**Supplementary Figure 8 B-G**). These findings suggest that live TDP-43^muts^ and control iMNs in culture achieved overall normal dendritic and axonal branching and morphology.

Previous reports found that patient-derived iMNs with *TARDBP* mutations degenerated and died at a faster rate than controls^74, 87^. To determine whether this was true in our TDP-43^mut^ iMNs as well, we evaluatedthe RGEDI ratio of red-fluorescing (dying) cells to the total number of green-fluorescing (live) cells in mutant and control cells over time (**Supplementary Figure 7B**). We calculated the odds ratio of cell death (OR-CD) over time as previously described^97, 106^. Briefly, the OR-CD was modelled using a generalized linear model (LM)^108^ to which we incorporated additional mixed effects to account for changes including the plate on which the cell was assayed, the time (as a continuous variable) at which the cell status was ascertained, the cell line from which the cell was derived and the interaction between the cell line and time^97, 106^. All TDP-43^mut^

iMNs displayed a higher proportion of dead and dying cells than their isogenic controls (OR-CD of 2.4) (**Figure 2B, Supplementary Figure 9**), even though two of the control lines displayed different death rates (**Supplementary Figure 9A**). The TDP-43^Q331K^ iMNs displayed the highest death rate compared to controls (OR-CD = 2.6), followed by TDP-43^M337V^ iMNs (OR-CD= 1.7) (**Supplementary Figure 9 B C**). The iMNs from the patient with the TDP-43^A382T^ mutation displayed an OR-CD of 1.4 compared to its gene-corrected control (GC-CTRL^1^), which was similar to our previous reports^106^ (**Supplementary Figure 9D**). While we did not observe significant changes collectively in ISLET1or NXK6.1 iMNs across the controls and all TDP-43^mut^ iMNs, there were significantly fewer NKX6.2-positive cells among the TDP-43^Q331K^ and TDP-43^M337V^ iMNs compared to their parental controls (**Supplementary Figure 6B**). Given that these lines also display the highest OR-CD, this finding is consistent with the hypothesis that NKX6.2-positive cells are the most vulnerable cells. Together these results suggest that patient-derived lines or genetically engineered lines with *TARDBP* ALS-causing mutations display evidence of neurodegeneration in culture, consistent with previous studies^74, 87^.

### Live cell morphology distinguishes TDP-43 mutants from control cell lines

Having established that the increased death rate of TDP-43^mut^ lines is readily observed in culture, we sought to identify cellular abnormalities that lead to neurodegeneration. To uncover these features, we subjected images acquired from RM to shallow fully connected ML algorithms (SML) in a custom-built pipeline we developed (**Supplementary Figure 10**). Images from TDP-43^mut^ and control cell lines in the EGFP channel were processed in our Galaxy software^113^ and cropped so that only the cell body and immediate processes were captured for each cell at two timepoints: T1 (∼ DIV27) and T6 (∼ DIV33) (**Figure 3A, Supplementary Figure 9**). We used a feature extraction that captures cellular properties such as cell size, shape, complexity and texture (**Supplementary Figures 11 A-P)**. We subjected crops from six independent experiments (**Supplementary Table 3**) that included TDP-43^mut^ iMNs and their isogenic controls to the Stacked Ensemble Classifier (defined in the SML section of Methods) at each timepoint (T1 and T6). After developing the model with the training and validation sets, we used it to predict classes (i.e. TDP-43^mut^ versus control cell) on the test set for each timepoint. We evaluated prediction accuracy using the area under the curve (AUC) and visualized the results using a receiver operator characteristic (ROC) curve, which shows the model’s ability to distinguish between classes across the two groups^181^. We observed an AUC score of 68% (**Figure 3B)** at T1 and 75% at T6 (**Figure 3C**). This may suggest that more mature neurons exhibit morphological characteristics that better reflect disease pathology than do earlier, more immature neurons.

**Figure 3.**
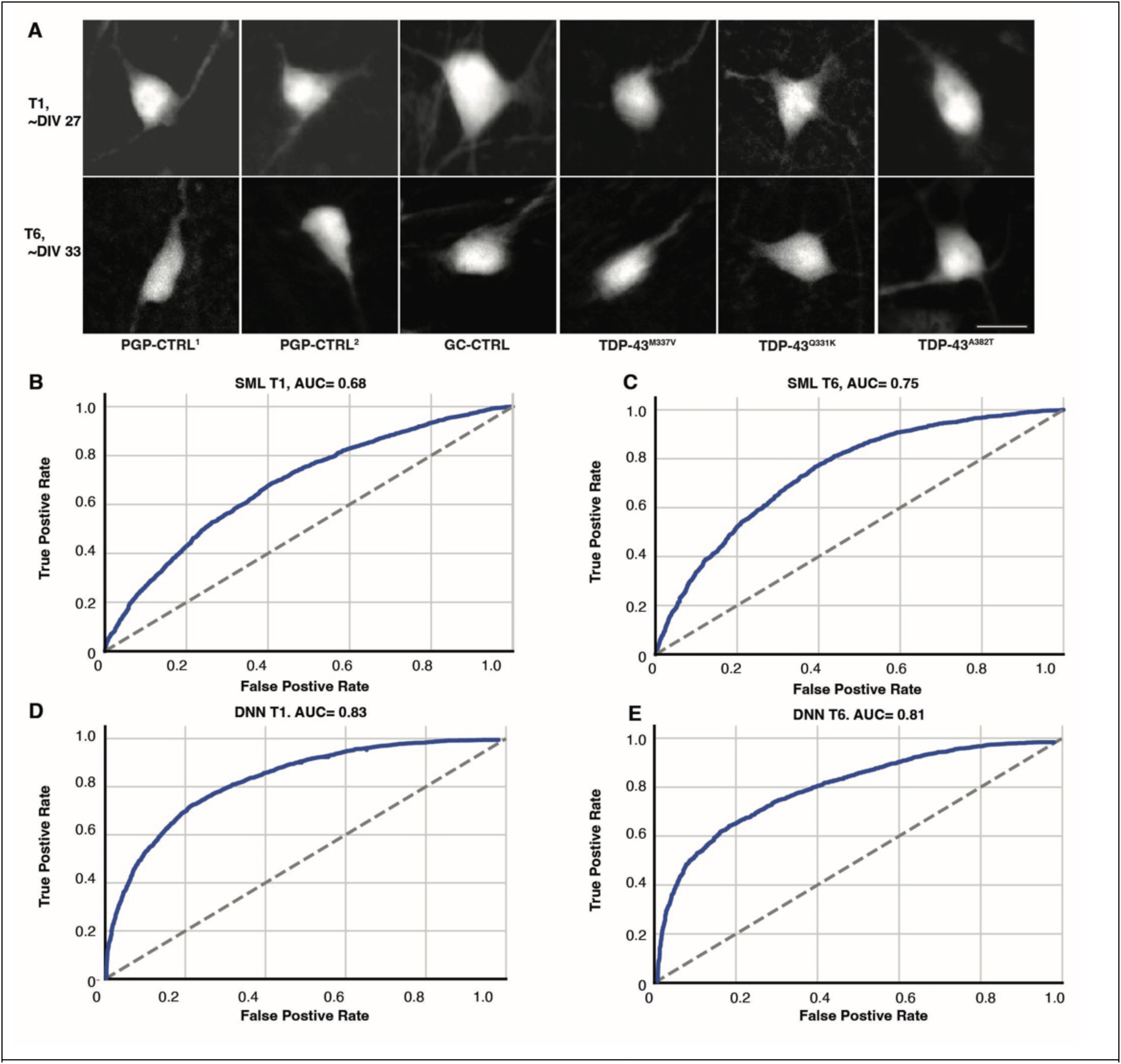
SML and ResNet DNNs distinguish TDP-43^mut^ iMNs from controls with moderate to high accuracy. **A**) Example of cropped images from time points T1 (DIV ∼27; top) and T6 (DIV ∼ 33; bottom) generated as input for the ML models. Scale bar= ∼11 µM. **B–C**) ROC curves (blue lines) showing accuracy of the indicated models at the validation step for the SML models at T1 and T6 respectively (See **Supplementary Table 3** for training/testing/validation splits of the datasets). **D-E**) ROC curves showing accuracy of ResNet18 at T1 and T6. Grey dashed lines indicate chance-level performance. Higher AUC values reflect improved discrimination between positive and negative classes.

Confusion matrices (CMs) revealed strong but imperfect separation of classes at both timepoints and confirmed that the predictive accuracy was higher at T6 than T1. The True Positive (TP) and True Negative (TN) rates at T1 were twice as high as False Positive (FP) and False Negative (FN) rates at T6, but slightly less than that at T1 (**Supplementary Figures 12A, D**). This may suggest that at the later timepoint, the model’s ability to identify either TDP-43^muts^ or controls iMNs is similar with no strong bias at either timepoint. However, there were fewer misclassification counts and the identification of TDP-43^muts^ cells improved subtlety at the later time point (**Supplementary Figures 12 A, D**). To confirm that the SML models use signals from the iMNs that reflect mutation class and are not just classifying based on artifacts, leakage, or structure in the data, we randomized the labels and found no true relationship between features and the labels exists (**Supplementary Figures 12 B, C, E, F**). Upon scrambling the labels, we find that the ratio of correctly classified crops, regardless of if they are control or TDP-43, is comparable to incorrectly classified crops, show that the models perform no better than chance (**Supplementary Figures 12 B, C, E, F**). This finding confirms that the discriminatory signal observed in the true labels is attributable to *TARDBP* mutation status.

While the results of SML were encouraging, we wondered whether we could achieve better discrimination of TDP-43^muts^ vs. control iMNs using deep neural networks (DNNs)^182^. Traditional SML algorithms rely on features engineered into the classifier (in our case, cell size, shape, complexity and texture). By contrast, DNNs^182^ can automatically extract hundreds of features, potentially offering greater predictive power. Typically, DNNs outperform SML algorithms in classification and pattern recognition tasks^183–185^.

We focused on four different models, ranging from older standards like VGG^186^ and ResNet18d^146, 147^ to the newer “transformer”-based ViT^187^ and MaxViT^188^, which are near the state-of-the-art on computer vision tasks like object recognition. Predictions from all DNNs except the ViT were significantly above chance using permutation testing (p < 0.001) (**data not shown**). The ResNet18d classifier^146, 147^ demonstrated the most robust performance, with high prediction accuracy at T1 (AUC = 0.83) and T6 (AUC = 0.81) (**Figure 3D, E**) and high discriminative abilities at both timepoints (**Supplementary Figures 12 G, H, K, L**). The model on T1 validation images achieved an AP of 0.83 and at T6 AP of 0.82. Confusion matrices for both T1 and T6 showed that TN and TP rates were much higher than the FP and FN rates (**Supplementary Figures 12 G, K**). We scrambled the labels across both timepoints and there was a complete loss of classification (**Supplementary Figures 12 I, J, M, N**). Collectively, these results indicate that the ResNet-based classifier performs better than our SML model at both time points, and detects features associated with TDP-43–related pathology more reliably than features characteristic of control samples.

We also evaluated SimCLR^145^, a contrastive self-supervised learning framework, to learn visual representations from TDP-43^mut^ and controls. The pretrained SimCLR model uses a more powerful ResNet with 50 instead of 18 layers (Res-Net50^146, 147^) as backbone. We again used the exact same dataset splits as for the above models (**Supplementary Table 3)**. The Res-Net50 DNN achieved higher precision across a wide range of recall values than did the SML models at both T1 (AUC = 0.79) and T6 (AUC = 0.81), but not as high as the Res-Net18 DNN (**Supplementary Figures 12 P, U)**. The ROC curves were 0.79 for both T1 and T6 models (**Supplementary Figures 12 Q, V**), indicating a robust ability to identify true positive TDP-43^mut^ samples while minimizing false positives, though not as reliably as the ResNet18 model. The confusion matrices at both T1 and T6 showed that the ResNet50 model exhibited slightly less sensitivity and specificity than both the ResNet18 the SML models however was still robust (**Supplementary Figures 12 O, T**). Furthermore, the predictive models again lost all predictive capacity when labels were scrambled, confirming that the learned signal reflects genuine biological structure rather than technical artifact. (**Supplementary Figures 12 R, S, W, X**).

Overall, these findings demonstrate that multiple ML algorithms and model architectures can robustly distinguish TDP-43^muts^ from controls, achieving high predictive accuracy and strong discriminatory performance. Label-scrambling analyses confirm that the discriminative signal arises from biologically meaningful differences within the cell types themselves, rather than from artifacts or model overfitting.

### The TDP-43 mutant–discriminating signal is nuclear and perinuclear

To understand the nature of the signal that distinguished TDP-43^muts^ from control iMNs, we employed an interpretability algorithm, SmoothGrad^189^, which allows one to identify which parts of the input (in this case, cropped images of iMNs from TDP-43^mut^ and controls) most influenced the model’s prediction. SmoothGrad produced an attribution map highlighting the regions in an image that are important for predicting each class, which we then refined with SmoothGrad^189^. In CTRL images (see 2 representative samples in **Figure 4A**, top row), SmoothGrad activation was weak, diffuse, and spatially inconsistent. Although faint hotspots could be detected, they were broad and variable in both intensity and localization. In contrast, in the TDP-43^mut^ iMNs (representative samples in bottom two rows of **Figure 4A**), SmoothGrad highlighted compact, high-intensity hotspots centrally located within the soma area that appeared to be the nucleus. These regions are markedly smaller and more sharply bounded than those of CTRL samples, mirroring the tight soma-centered activation pattern observed across all TDP-43^mut^ images in the dataset. The observed visual consistency across the 245 TDP-43^mut^ examples further supports that the model is capturing a robust, recurring ALS-specific morphological signature. This technique suggested that most of the signal that distinguishes TDP-43^mut^ iMNs from isogenic controls is in the perinuclear and nuclear area (**Figure 4A**), suggesting alterations in the nucleus, chromatin or nuclear pore.

**Figure 4.**
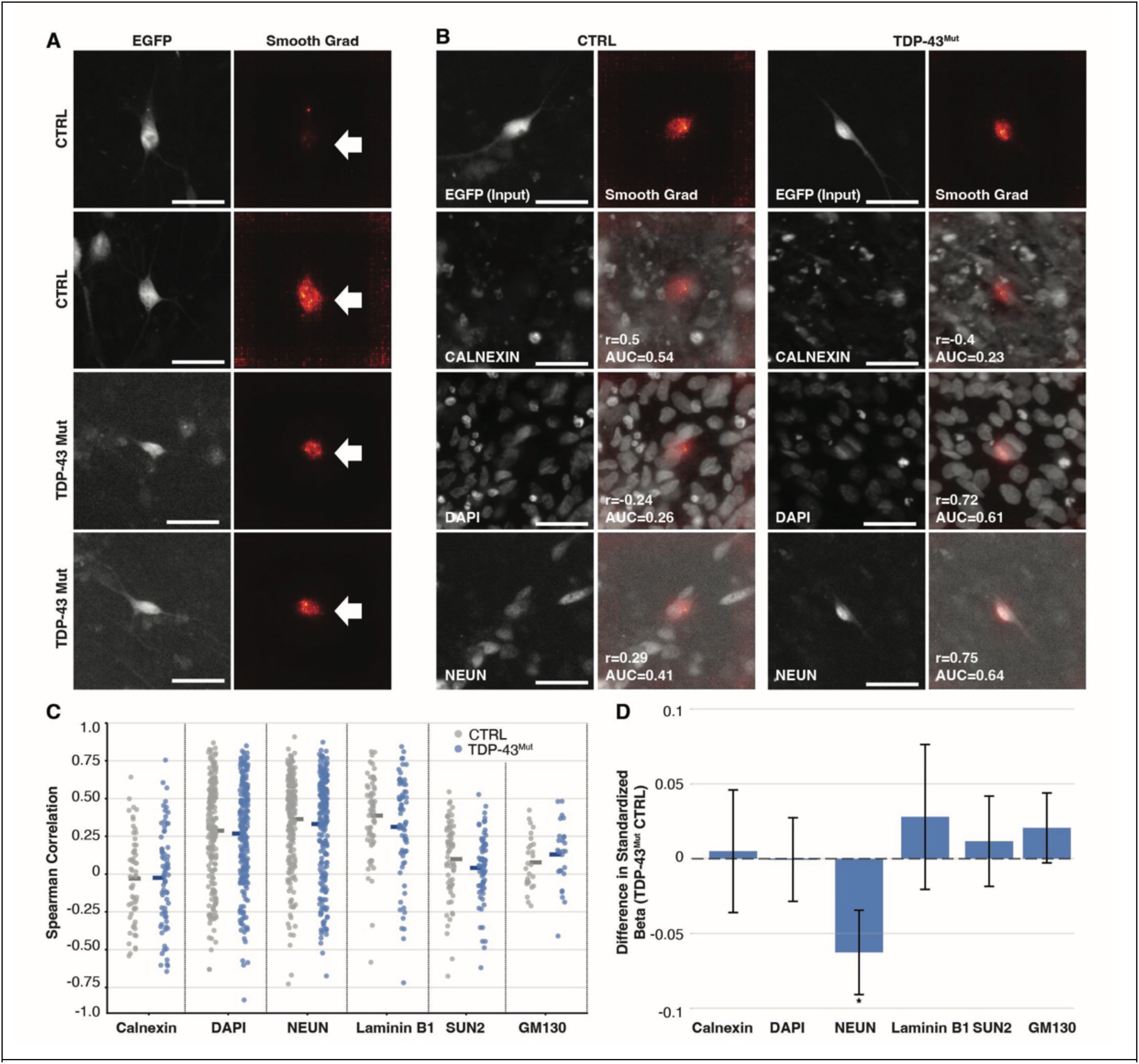
Feature attribution maps reveal that the classifying signal of TDP-43mut iMNs from controls is perinuclear. **A**) ***Left*** Representative fluorescence images showing the morphology (EGFP portion of GEDI) of control and TDP-43^mut^ iMNs. ***Right*** Model-derived SmoothGrad saliency maps of the same cells highlighting regions contributing most to classifier decisions (arrows). Scale bar = ∼11uM. **B**) Overlap between saliency maps (right panels) and CALNEXIN, DAPI or NEUN staining (left panels) of control and TDP-43^mut^ iMNs. For each marker, Spearman correlation (π) and AUC indicate the degree to which the salient regions overlap with specific subcellular features. **C**) Partial R² for each stained compartment of the cell for control (blue) and TDP-43^mut^ (red) iMNs (n=463 cells; Control n=218, TDP-43^mut^ n=245). Each dot represents a single cell. NeuN and DAPI showed the highest partial R² values in both groups, indicating that nuclear markers best explained the model’s attribution signal. **D**) Differences in standardized regression coefficients (β) between TDP-43^mut^ and CTRL cells for each cellular marker. Positive values indicate markers with stronger contributions to predicting the mutant condition, whereas negative values indicate features more associated with CTRL cells. Error bars represent standard error. Only NEUN’s value is statistically significant (p = 0.014, permutation test)

To better pinpoint the origin of the discriminative signal, we carried out a series of ICC stains using antibodies that label different cellular compartments and performed a partial correlation between the model’s feature attribution map and each stain^190^. The antibodies recognized NEUN (nucleus and perinucleus^191, 192^), GM130 (Golgi^193^), Calnexin (ER^194^), SUN-2 (inner nuclear membrane (INM^195^), Lamin B1 (nuclear envelope^196, 197^) and DAPI (nucleus^198^) (**Supplementary Figure 13 A, B, C, D**). We then generated single cell cropped images of cell bodies including the EGFP signal (from GEDI) and the signal from each antibody (**Supplementary Figure 14 A, B, C, D).** Across multiple fields of view, the EGFP morphology channel showed variable degrees of alignment with nuclear and neuronal stains, and weaker relationships with markers of cytoplasmic or organellar compartments (**Figure 4B and Supplementary Figure 15 A**).

To quantify the differences between the model’s feature-attribution map and TDP-43^mut^ signal in iMNs and control cells, we computed semi-partial correlations between the attribution heatmaps and the fluorescence signals from stained cells, controlling for background and other confounding channels (**Figure 4 C, D and Supplementary Figure 15 A, B, C**). We observed differences in the cellular features driving predictions for TDP-43^mut^ vs control iMNs (**Figure 4 C)**. TDP-43^mut^ iMNs (blue dots) showed lower contributions of DAPI, NeuN, and Lamin B1 than did controls (grey dots). Spearman correlations between stain intensity and gradient magnitude were modest overall, but consistently positive for nuclear markers (**Supplementary Figure 15 D**). Likewise, mean AUC-of-overlap (AUC-o) values indicated that the spatial distribution of SmoothGrad saliency was best captured by nuclear and neuronal stains (**Supplementary Figure 15 E**). By contrast, GM130, Calnexin and SUN2, showed the lowest gradient–stain correspondence across metrics.

To examine group differences, we computed partial R² values and estimated each stain’s unique contribution to explaining gradient variance. Nuclear and neuronal markers accounted for the largest proportion of explained gradient variability, with NeuN and DAPI showing the strongest contributions. Comparisons between groups revealed significantly elevated partial R² values in TDP-43^mut^ iMNs relative to controls for several markers, most prominently NEUN and Lamin B1 (**Supplementary Figure 15 F**).

These patterns were corroborated by hierarchical clustering of partial R² values, which segregated samples by condition and emphasized a distinct contribution of nuclear morphology and neuronal identity to classifier decision-making. We observed that there was an increased reliance on the nuclear label NEUN and to a lesser degree DAPI and Calnexin and Lamin B1 (**Supplementary Figure 15 F**). Together, these results suggest that nuclear and neuronal features disproportionately influence the model’s predictions and that these features are differentially leveraged in TDP-43^mut^ iMNs.

Next, we performed standardized regression coefficients to examine which stains were most correlated with each class. TDP-43^muts^ showed increased contributions of LaminB1 and DAPI, although these differences did not reach statistical significance (**Figure 4D**). In contrast, NeuN exhibited a significantly weaker association with the attribution map in TDP-43^muts^ compared to controls (p = 0.014, via a permutation test). This pattern indicates that cellular alterations that occur in the TDP-43^muts^ selectively disrupt the relationship between the attribution map gradient and NEUN. Overall, our findings highlight the nucleus and perinuclear regions as the most discriminative sources of signal for distinguishing TDP-43 mutants from controls.

We also confirmed the relative contribution of cellular and subcellular markers to variance across control and TDP-43 mutant conditions, by visualizing the computed partial R² values as a clustered heatmap (**Supplementary Figure 15 G**). Across all samples, NEUN consistently exhibited the largest absolute partial R² values, indicating that nucleus–associated features accounted for a substantial proportion of explained variance. DAPI also showed moderate contributions across multiple samples, corroborating that the nuclear content was an additional driver of variability.

### TDP-43 mutant iMNs display evidence of nucleocytoplasmic dysfunction

The identification of a discriminating signal in the nuclear and perinuclear regions suggested the presence of pathological alterations in or near the nucleus. Since nuclear pore dysfunction is likely an underlying cause of TDP-43 pathology in ALS^199–202^, we therefore examined nucleocytoplasmic transport (NTC) using a fluorescent nucleocytoplasmic shuttling biosensor called 2Gi2R^96^ (**Figure 5A**). This construct contains a nuclear localization motif tethered to the RFP and a nuclear export signal tethered to GFP (**Figure 5A**). The two signals are encoded on the same transcript and therefore expressed at a 1:1 stoichiometry, thus allowing us to evaluate the fidelity of NCT by measuring the RFP/GFP ratio in each cell. We transduced the 2GiR biosensor into ∼DIV 34 iMNs and imaged the cells using RM. We quantified the spatial occupancy of each fluorophore within GFP- and RFP-positive iMNs (**Figure 5B and Supplementary 16A**). The RFP/GFP ratio was higher in TDP-43^mut^ than control iMNs (**Figure 5C and Supplementary 16B**). Since the RFP/GFP ratio compares the size of the areas fluorescing in red versus green, this finding could reflect an enlargement of the nucleus of the mutant iMNs. To eliminate this possibility, we measured the size of the nuclei, as revealed by DAPI signal, in the same cell lines. While we observed some difference in nuclear size between the TDP-43^Q331K/M337V^ iMNs and their controls, and between the patient derived TDP-43^A382T^ iMNs and their gene-corrected control (**Supplementary 16C)**, those differences did not reach significance when cells from the 3 mutant lines combined were compared to cells from the 3 control lines (**Figure 5D**). These differences were also not found across gene-corrected iMNs compared to the TDP-43^A382T^ nor in the gene-edited lines compared to the parental control (**Supplementary 16C)**. The increased RFP signal size in the TDP-43^muts^ iMNs could be due to leakage of the nuclear reporter, presumably as the consequence of a dysfunctional NCT. However, as this reporter is constanly shuttling between the nucleus and the cytoplasm this result could also be due to reduced nuclear import, which will lead to a new steady state and redistribution over time. Alternaltivley, both scenarios could be happenieng. Neverthelss the fact that we narrowed the mutated TDP-43 predictive signal to the nucleus and that this led us to identify NCT defects in the TDP-43^muts^ illustrates the promise of ML to detect pathological signals that can help us better understand the mechanisms of ALS.

**Figure 5.**
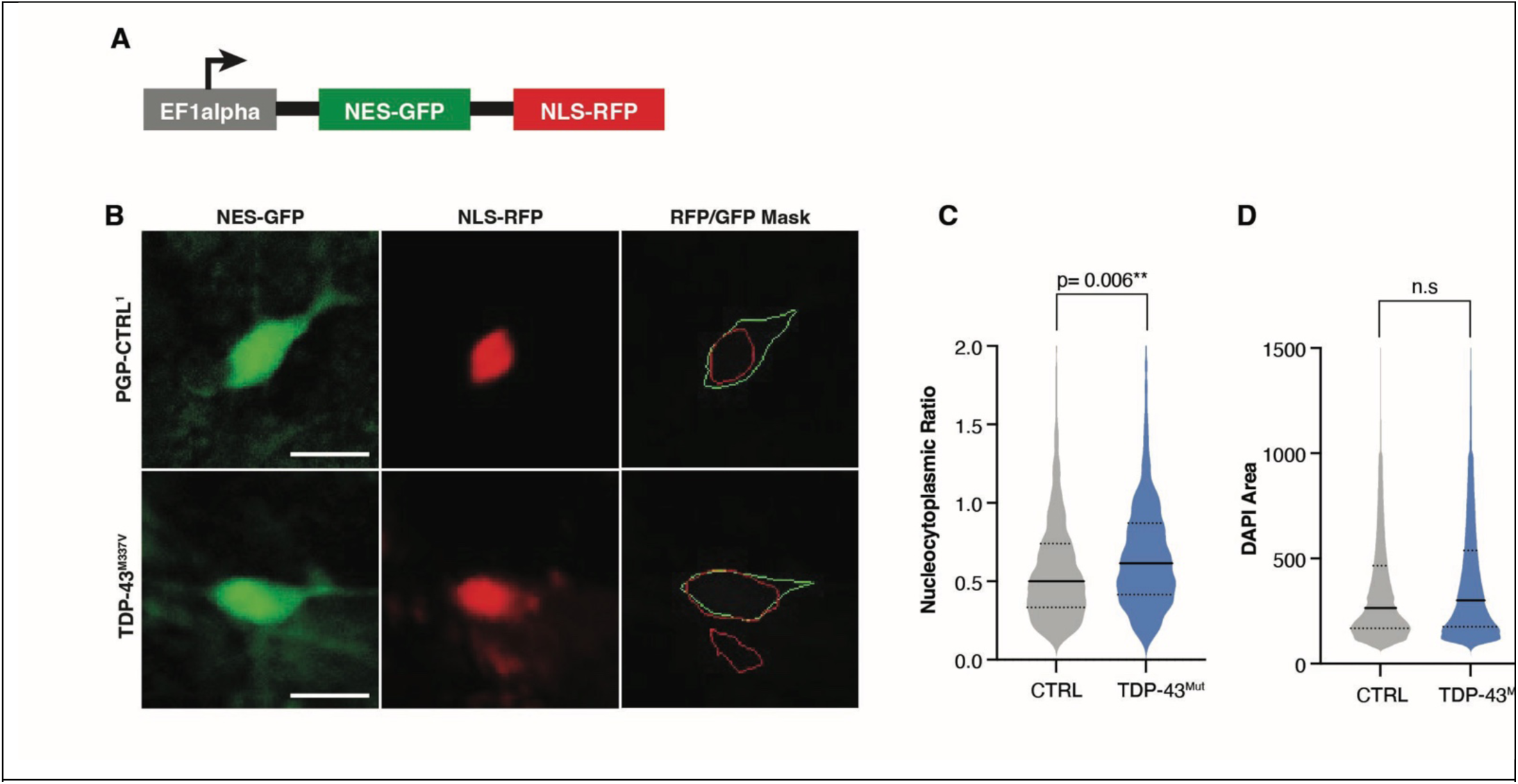
TDP-43^mut^ iMNs display cytoplasmic mislocalization of the nucleocytoplasmic shuttling biosensor compared to isogenic controls. **A**) Cartoon illustrating the NCT shuttling biosensor. **B**) Representative images showing the distribution of NES-GFP and NLS-RFP expressed from the 2GiR construct in a control and TDP-43^Q331K^. The far-right panels show the outlines of the masks of the two channels, RFP and GFP. Scale bar = 15 µm. **C**) The NC ratio (ratio of RFP to GFP mask) is significantly higher in TDP-43^mut^ iMNs. See also **Supplementary** Figure 14. Violin plots show the NCT ratio of each cell across the three TDP-43^mut^ lines and controls. Estimate = 0.16, p = 0.006, n = 4 experiments; Control = 2,467 cells; TDP-43^muts^ = 2,018 cells. **D**) Nuclear size is not significantly different in TDP-43^mut^ lines compared to isogenic controls. n= 4 experiments, Control = 444,989 cells, TDP-43^muts^ = 293,515 cells. Estimate = 0.04, p= 0.68. All statistical analyses were performed using a GLMM^110^.

### sALS and C9orf72^mutant^ iPSCs pattern normally into motor neurons

We surmised that if ML models could distinguish iMNs harboring *TARDBP* mutations from controls, they might also detect discriminating features of other forms of ALS. We obtained from Answer ALS^87^ iPSC lines from ALS patients carrying *C9ORF72* expansions (C9orf72^muts^) and from ALS patients with no known ALS-causing genetic mutations (sALS), as well as unrelated control lines from donors with no history of ALS. We chose lines from ALS patients that had early to typical age of onset (AO) (45 to 68 years old) and age-matched controls (**Figure 1 and Supplementary Table 1**). We differentiated the cells to iMNs and after ∼5 weeks in culture we stained them with various neuronal and motor neuron markers as we had done for the TDP-43^mut^ iMNs and controls. We observed robust MAP-2 staining and moderate expression of motor neuron markers such as ISLET1 and NKX6.2 across all lines (**Supplementary Figures 17, 18**). In addition, the cultures contained very low numbers of KI67-positive cells (**Supplementary Figure 19**). For both ISLET1 and NKX6.2, the overall number of stained cells averaged 25% and ranged between cell lines from ∼15 to 50% (**Supplementary Figure 20, A, B, D,E**), indicating that a subset of the cells patterned toward a motor neuron-like fate, consistent with previous reports^87^. The C9^1^ line showed a higher number of NKX6.2-positive cells than the CTRL^1^ control (**Supplementary Figure 20 D**). Given the variability observed among all ALS lines, this difference is likely not biologically meaningful. Consistent with this notion, when we combined the C9orf72^mut^ and sALS lines and compared them to controls, we did not observe significant differences in the number of ISLET1 and NKX6.2 iMNs across the groups (**Supplementary Figure 20 C F**). We also observed TDP-43 staining in C9orf72^muts^ and a subset of sALS compared to controls (**Supplementary Figure 21)**. The average frequency of TDP-43–positive cells ranged from ∼35 to 60% (**Supplementary Figure 20 G, H).** We observed a subtle but significant increase in the number of TDP-43–expressing cells in sALS compared with controls (**Supplementary Figure 21 H)**, consistent with similar reports in sALS patients^203, 204^.

To examine the iMNs’ morphology, we transduced them with RGEDI as described above. We found that they displayed typical neuronal morphology with dendrites and branching processes (**Figure 6A**). At DIV∼30, we observed no significant differences in neurite length, number of processes or branch ends between the C9orf72^mut^, sALS iMNs and controls (**Supplementary Figure 22 A, B, D, E, F**). We observed a very subtle increase in the number of trunks per neuron in the C9orf72^muts^ but not sALS iMNs compared to control iMNs (**Supplementary Figure 22 C**). Overall, these findings suggest that the sALS and C9orf72^mut^ genetic backgrounds do not affect the cell lines’ ability to differentiated into morphologically normal iMNs.

**Figure 6.**
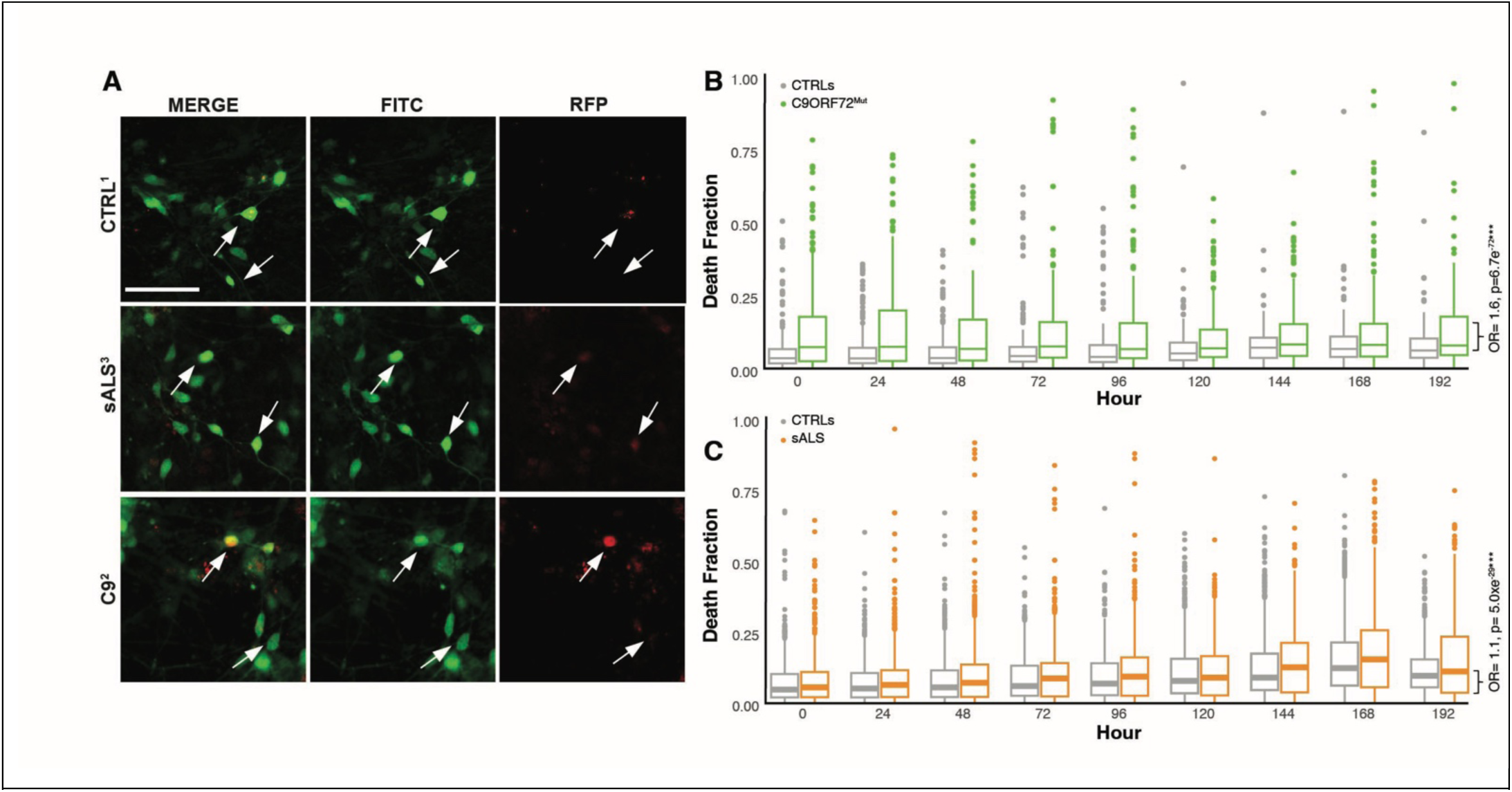
sALS and C9orf72^muts^ iMNs display a higher rate of cell death than control iMNs. **A**) Representative images of iMNs from one sALS, one C9orf72^mut^ and one control line expressing the GEDI construct. Cells are viewed in the FITC channel (morphology; middle panels), RFP channel (cell death; right panels), or merged channels (right panels) (scale bar = 74 µM). Arrows indicate dead or dying cells expressing elevated levels of RGEDI. **B, C**) Distribution of death fraction in iMNS combined from (**B**) 2 C9orf72^mut^ lines (C9^1^ and C9^2^) vs. 2 control lines (CTRL^1^ and CTRL^2^), and from (**C**) 7 sALS lines (sALS^1-7^) vs. 6 control lines (CTRL^1-6^). Boxplots display the median and interquartile range; whiskers represent data range. Individual dots show outlier wells. **B**) C9^1-2^ lines display a higher OR-CD than controls (OR = 1.6, p = 6.7xe^-72^; n = 14 experiments, 435,116 live cells, 37,767 dead cells. **C**) sALS^1-7^ iMNs display a moderately higher OR-CD than CTRLS^1-6^ (OR= 1.1, p= 1.0xe-29***, n=19 experiments, 1,773,578 live cells, 197,794 dead cells.)

### sALS and C9orf72^mut^ iMNs degenerate in the absence of stressors

Previous studies observed heightened neurodegeneration of fALS iMNs with *C9ORF72* mutations^205^, as well as sALS iMNs under stress conditions such a growth factor withdrawal^206^. We used RGEDI to monitor cell death in iMNs from 6 sALS, 2 C9orf72^mut^ and as many control lines cultured without growth factor removal and calculated the OR-CD^97, 106^ as described above. The two C9orf72^muts^ displayed a greater OR-CD than the controls (OR-CD = 1.6; **Figure 6B**), and a slightly different OR-CD from one another (**Supplementary Figure 23A**). OR-CDs were highly variable among sALS lines but proved overall subtly yet significantly higher than those of controls (**Figure 6C, Supplementary Figure 23 B, C**). As the OR-CD of the sALS lines compared to controls was low (1.1), we corroborated these results by hand-curating a subset of the lines and subjecting them to Kaplan-Meir and COX-proportional hazards analysis to evaluate the cumulative risk of death^62–71, 87, 102, 156^. By this secondary metric, we observed a significant increase death rate in a subset of the sALS iMNs compared to controls (HR = 1.4) (**Supplementary Figure 23 D**).

The six sALS lines were derived from different patients with different disease outcomes and, presumably, different etiologies. We therefore wondered if death rate in these lines correlated to clinical outcomes. Linear regression and Spearman’s correlation did not reveal any correlation between age of onset and rate of cell death (**data not shown**). On the other hand, we found a significant negative correlation between the overall fraction of dead cells and the proportion of ISLET1-positive iMNs found at the conclusion of live cell imaging (Spearman r = −0.81, p = 0.0011) (**Supplementary Figure 24 A**), suggesting that the ISLET1-positive iMNs are the cells most susceptible to degeneration. We did not observe a similar correlation for the NKX6.2-positive iMNs (**Supplementary Figure 24 B**).

We next asked whether the neurodegeneration observed in sALS iMNs was solely attributable to ISLET1 expression, or if all sALS lines had an increased death rate relative to controls, regardless of ISLET1 levels. To answer this question, we stratified the lines into “low-frequency ISLET1 expressors” (CRTL^1^, CTRL^3^, CTRL^4^, sALS^2^, sALS^5^, sALS^6^), defined as those with an average of ≤ 20% ISLET1-positive cells, and “high-frequency ISLET1 expressors” (CTRL^2^, CTRL^5^, CTRL^6^, sALS^1^, sALS^3^, sALS^4^) defined as those with an average of ≥ 25% ISLET1-positive cells. In both groups, sALS iMNs exhibited a significantly higher OR-CD than did control iMNs (**Supplementary Figure 24C, D**). Notably, the difference in cell death between sALS and control iMNs was more pronounced among the low-frequency ISLET1 expressors, suggesting a connection between the fraction of motor-neuron-like identity and neurodegeneration. This may be because as sALS iMNs more effectively acquire a motor neuron like identity, they become increasingly vulnerable to degeneration, such that fewer cells of this type survive to the end of the experiment, when fixation and staining occur. Overall, these findings indicate that the heightened neurodegeneration in sALS iMNs reflects an intrinsic disease-related vulnerability, rather than being solely driven by ISLET1 expression levels. However, the correlation between low rate of ISLET expression and higher death rates suggests that the vulnerability to neurodegeneration may, in part, be driven by cell-type specific factors.

### SML and Deep DNNs moderately discriminate sALS and C9orf72 mutants from controls

We postulated that as for TDP-43^mut^ iMNs, there may be a morphological signal underlying neurodegeneration in the sALS and C9orf72^mut^ iMNs and distinguishing them from controls. Therefore, we generated single-cell crops from live cell images acquired using RM at ∼DIV 28 and 33 (**Figure 7A)**, split the images 80/10/10 into train/test/validation sets (**Supplementary Table 3**) and trained both SML and the ResNet50 DNNs to discriminate C9orf72^mut^ iMNs or sALS iMNs from controls (**Figure 7A)**. The C9orf72^mut^ dataset came from 8 individual experimental replicates (**Supplementary Table 3**). The SML model showed a moderate performance at T1 (AUC = 0.60) (**Figure 7A)** with CMs indicating near-balanced classification of CTRL and C9orf72^mut^ iMNs (**Supplementary Figure 25 A**). However, when trained and tested on scrambled images, classifier performance deteriorated to be no better than chance-level performance (**Supplementary Figure 25 B**). Consistent with this the ROC for the scrambled condition yielded an AUC = 0.49 which was entirely flat (**Supplementary Figure 25 C**). In contrast, the ResNet-50 model at T1 demonstrated improved classification accuracy (**Figure 7B)** with ROC = 0.68 with and an improved separation between CTRL and C9orf72^mut^ (**Supplementary Figure 25 D**). The PRC analysis showed an AP of 0.64, well above the no-skill baseline (**Supplementary Figure 25 E**). Notably, performance metrics for the ResNet-50 model trained on scrambled images indicate a loss of discriminatory ability (**Supplementary Figure 25 F, G**). suggesting features that capture C9orf72 pathology are present at subtle, but genuine levels.

**Figure 7.**
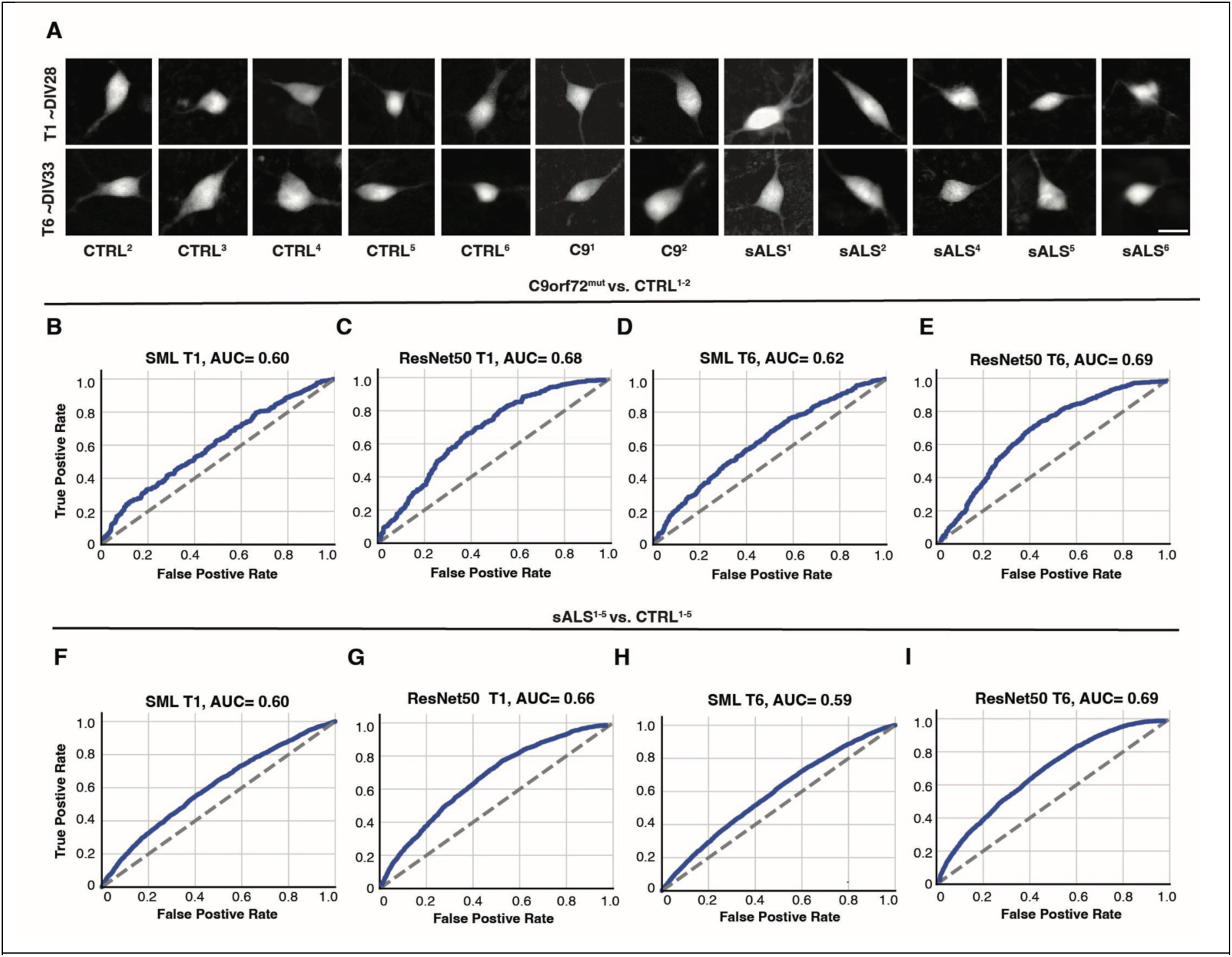
SML and Resnet50 DNN moderately discriminate sALS and C9orf72^mut^ iMNs from controls. **A**) Examples of cropped images used to train both SML and ResNet50 DNN. Scale bar= ∼11 µM. **B-E**) ROC curves showing the ability of SML (**B**, **D**) and ResNet50 DNN (**C**, **E**) to classify the same set of cells accurately as C9orf72^mut^ iMNs vs. controls at early (T1) and later (T6) time points. AUC values range between 0.60 and 0.68, indicating moderate performance. The ResNet50 DNN T6 model achieved the highest AUC (AUC = 0.69). **F–I**) ROC curves showing the ability of SML (**F**, **H**) and ResNet50 DNN (**G**, **I**) to classify the same set of cells accurately as sALS vs. CTRL iMNs at T1 and T6. AUC values range between 0.59 and 0.69, indicating modest performance. ResNet50 DNN model outperforms SML at both T1 (AUC = 0.66) and T6 (AUC = 0.69). For all AUC curves, blue traces represent model performance (true positive rates) across threshold values (false positive rates), and grey

At the later T6 time point, SML classifiers trained on control and C9orf72^mut^ iMNs also showed moderate classification capability (**Figure 7D**) but displayed improved correct classification of C9orf72^mut^ iMNs relative to CTRL (**Supplementary Figure 25 H**). Again, scrambling the images completely abolished this signal, resulting in a collapsed ROC AUC of 0.49 (**Supplementary Figure 25 I, J**). The ResNet-50 model at T6 trained on the same images achieved a more robust classification performance with an AUC = 0.69 (**Figure 7E**), with CMs showing strong prediction accuracy for both classes (**Supplementary Figure 23 H**). PRC yielded an AP of 0.64 (**Supplementary Figure 25 H**), consistent with the performance observed at T1 and indicating sustained discriminative capacity over time. Image scrambling again reduced classification to random levels (**Supplementary Figure 25 M, N**).

We next investigated the performance of these models in classifying ALS among individuals without known genetic mutations. Cropped images from six control lines and seven sALS lines were generated from 8 independent experiments. Three lines produced insufficient image numbers and created class imbalance; therefore, these were excluded. The models were therefore trained on five sALS lines and five control lines. SML performance remained modest to classify sALS from control at T1 (AUC = 0.60) (**Figure 7 F**) and the CM displayed limited class separation, with substantial overlap between control and sALS predictions (**Supplementary Figure 26 A**). However, scrambling the images completely reduced classification ability as shown by the collapsed ROC (**Supplementary Figure 26 B, C).** The ResNet DNN models demonstrated improved predictive capacity to classify sALS from control at T1 (AUC = 0.66) (**Figure 7G**) showing with higher correct classification rates for both CTRL and sALS iMNs (**Supplementary Figure 26 D**). PRC yielded an AP of 0.63, exceeding the no-skill baseline (**Supplementary Figure 26 E**). Performance metrics for the ResNet-50 model trained on scrambled images flatlined and the models classified no better than chance (**Supplementary Figure 26 F, G**).

At the T6 time point, SML classifiers again showed modest performance when trained on sALS versus control images (**Figure 7G**), however with a slightly better discrimination towards sALS (**Supplementary Figure 26 H**). Image scrambling abolished this trend (**Supplementary Figure 26 I, J**). By contrast, the ResNet-50 model at T6 exhibited more robust classification (AUC = 0.69) (**Figure 8 I**) and more consistent discrimination correctly identifying a large fraction of both CTRL and sALS samples (**Supplementary Figure 26 K**). PRC analysis revealed an AP of 0.66 (**Supplementary Figure 26 L**), indicating enhanced discriminative power at later stages of differentiation. The model flatlined upon image scrambling (**Supplementary Figure 26 M N**).

**Figure 8.**
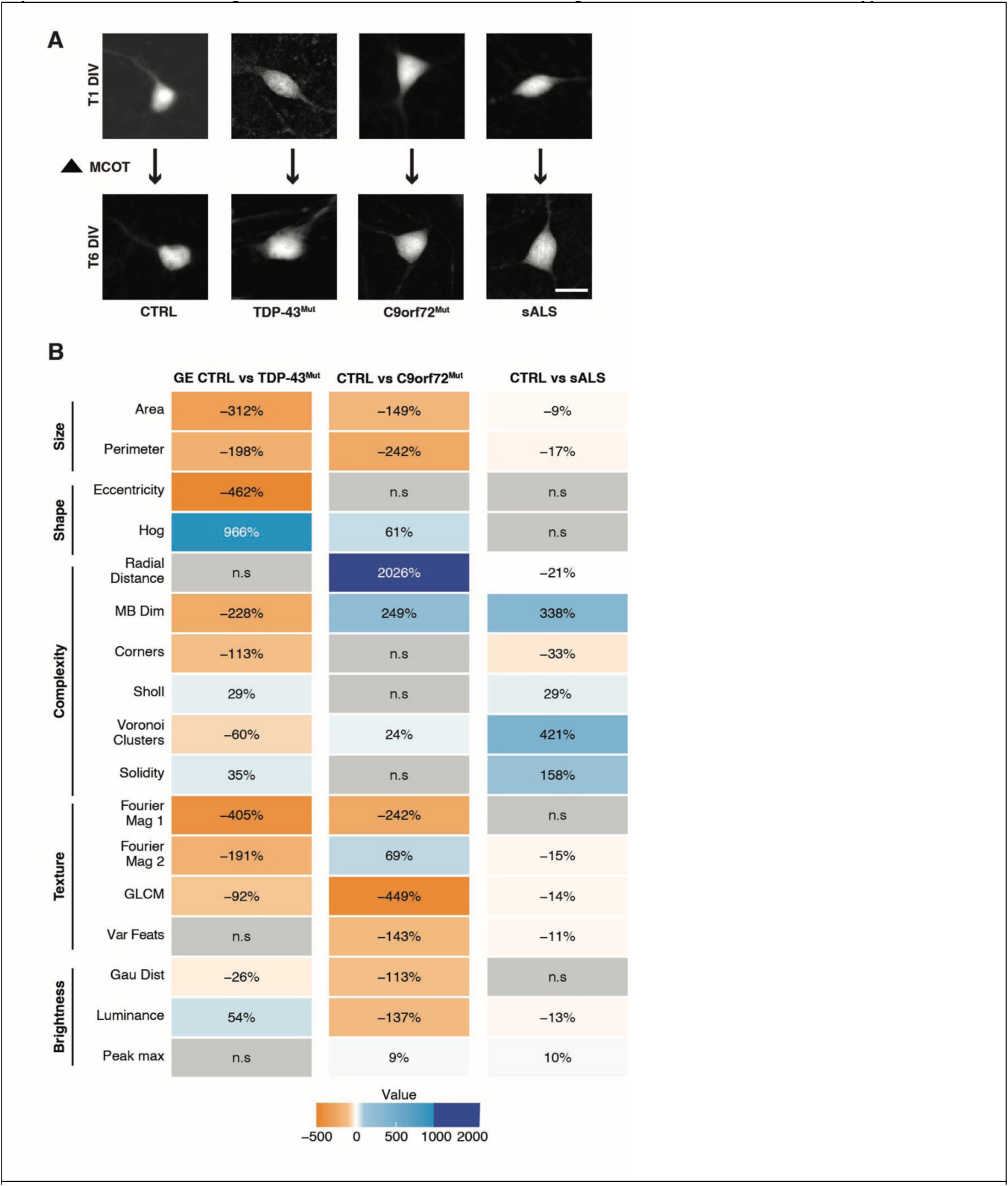
Longitudinal morphological profiling reveals distinct and genotype-specific alterations in ALS iMNs. **A**) Representative fluorescence images of neurons from control (CTRL), TDP-43^muts^, C9orf72^muts^, and sALS groups at T1 and T6 (∼DIV 27, DIV 33). Scale bar= ∼11 µM. **B**) Heatmap depicting relative percentage differences in quantitative morphological features between CTRL and each ALS subgroup, organized by feature class (size, shape, complexity, texture, and brightness). Positive (blue) and negative (orange) values indicate increases or decreases relative to CTRL, respectively, with color intensity reflecting effect magnitude. Gray boxes denote non-significant (n.s.) comparisons. Feature values extracted from SML models applied on crop images described above that were used for ML (see **Supplementary Tables 3 and 4**) were modeled using a linear mixed-effects model with random intercepts for experimental batch and cell line. Fixed effects included disease status (ALS vs CTRL), timepoint, and their interaction. The interaction term (time × disease status) captures the additional change over time in disease groups

Across both ALS groups, the ResNet DNN models consistently outperformed SML in terms of ROC AUC and resulted in better discrimination as observed by the confusion matrices. This is also reflected when observing the specificity/sensitivity scores as calculated for each ALS group, which we are using to estimate per class accuracy. ResNet50 tends to outperform SML except for sALS T1 (higher specificity) and sALS T6 (higher sensitivity). This divergence in sALS data implies that initially, SML is better at identifying controls but at a later timepoint, it is instead better at identifying sALS samples. This may be explained by the pathological signature being subtle and closer to the control distribution at T1, so our SML learned conservative decision boundaries that favored correct rejection of positives, resulting in higher specificity. Then, at T6, sALS-related features become more pronounced and more linearly separable.

Even so, the fact that scrambling the images rendered all models entirely nonfunctional, as shown by a complete collapse of the ROC curve and CMs showing no discriminatory ability, suggests that an underlying morphological signal is driving some degree of disease-specific classification, even if the models themselves are limited. Interestingly, both models demonstrated better performance at T6 than T1, indicating that ALS-associated phenotypes may take time to develop in culture. However, despite consistent improvements, overall classification performance remained moderate which suggests that C9orf72 and sALS-associated morphological signatures are more complex than the signatures of iMNs with TDP-43 mutations.

### Feature-based time interaction models reveal robust changes in morphology over time for TDP-43^mut^, C9orf72^mut^ and sALS compared to controls

Although ALS typically presents later in adulthood and follows a rapidly progressive disease course after symptom onset^207–210^, disease-associated pathology has been detected before clinical onset in individuals with C9orf72 mutations^211^ and other forms of ALS^212^. In human ALS models, alterations in gene expression were reported early on during neuronal development and maturation^213^ and TDP-43 pathology and associated gene expression changes and cryptic exon splicing become more severe as iMNs mature^213^.

We first compared feature weights across individual time points and across the three sub-models within the SML stacked classifier to determine whether any distinguishing features were specific to particular groups. Although some subtle differences were observed, no clear or consistent pattern emerged (**Supplementary Table 4**). We therefore wondered whether features associated with longitudinal morphological changes would provide a more sensitive characterization of intergroup differences than cross-sectional, single time-point assessments. In order to identify temporal phenotypic changes associated with TDP-43^muts^, C9orf72^muts^ and sALS versus control iMNs, we used a custom pipeline we previously developed that quantifies dynamic changes such as size, shape, complexity and texture over time^84^. Using the features extracted from our SML models (**Supplementary Figure 10 A-P**) for each image described above, we estimated the change in these morphological readouts from T1 to T6 (T1 →T6) for each group of cropped images for TDP-43mut, C9orf72^mut^ and sALS compared to control iMNs differentiated in parallel (**Figure 8 A)**. To estimate this shift in the feature space across the two time-points for each group, we used a Linear Mixed Effects Model (LMM), which models the mean change in disease status via an interaction term^109, 155^. We call this shift the <u>m</u>orphology <u>c</u>hange <u>o</u>ver time or MCOT^84^. The time-by-disease-status interaction terms were scaled to represent the MCOT in the TDP-43^muts^, C9orf72^muts^ and sALS versus control and expressed as percentage of change (**Figure 8)**. In this way, we developed a comprehensive panel of morphological, structural, and texture-based features over time across the different ALS groups and calculated percent deviation from controls. These changes may reflect dynamic differences in how the iMNS are patterning and behaving as they grow or degenerate. Further, because we capture changes over time, the signal is substantially amplified compared to static, single–timepoint measurements.

We grouped the SML features based on various characteristics such as those that capture size, shape, texture or brightness of objects within each crop (**Figure 8 and Supplementary Figure 10** and methods section). We used the same image crops that were used in the SML and DNN methods (**Supplementary Table 5)** but in this case we modeled the MCOT of each feature. For example, the area and perimeter of all control iMNs increased from T1 to T6 while in all ALS groups, they remained more static (**Figure 9B and Supplementary Figure 27 A, B).** Interestingly, the change in area varied in size but was constant in direction across all groups, with the largest effect shown in TDP-43^muts^ (312% change compared to controls) followed by C9orf72^muts^ (149% change) and sALS (a very moderate 9% change). The change in cell perimeter displayed similar trends (**Figure 8B and Supplementary Figure 27 B).** Together this suggests that the ALS lines grow much more slowly than the control lines.

**Figure 9.**
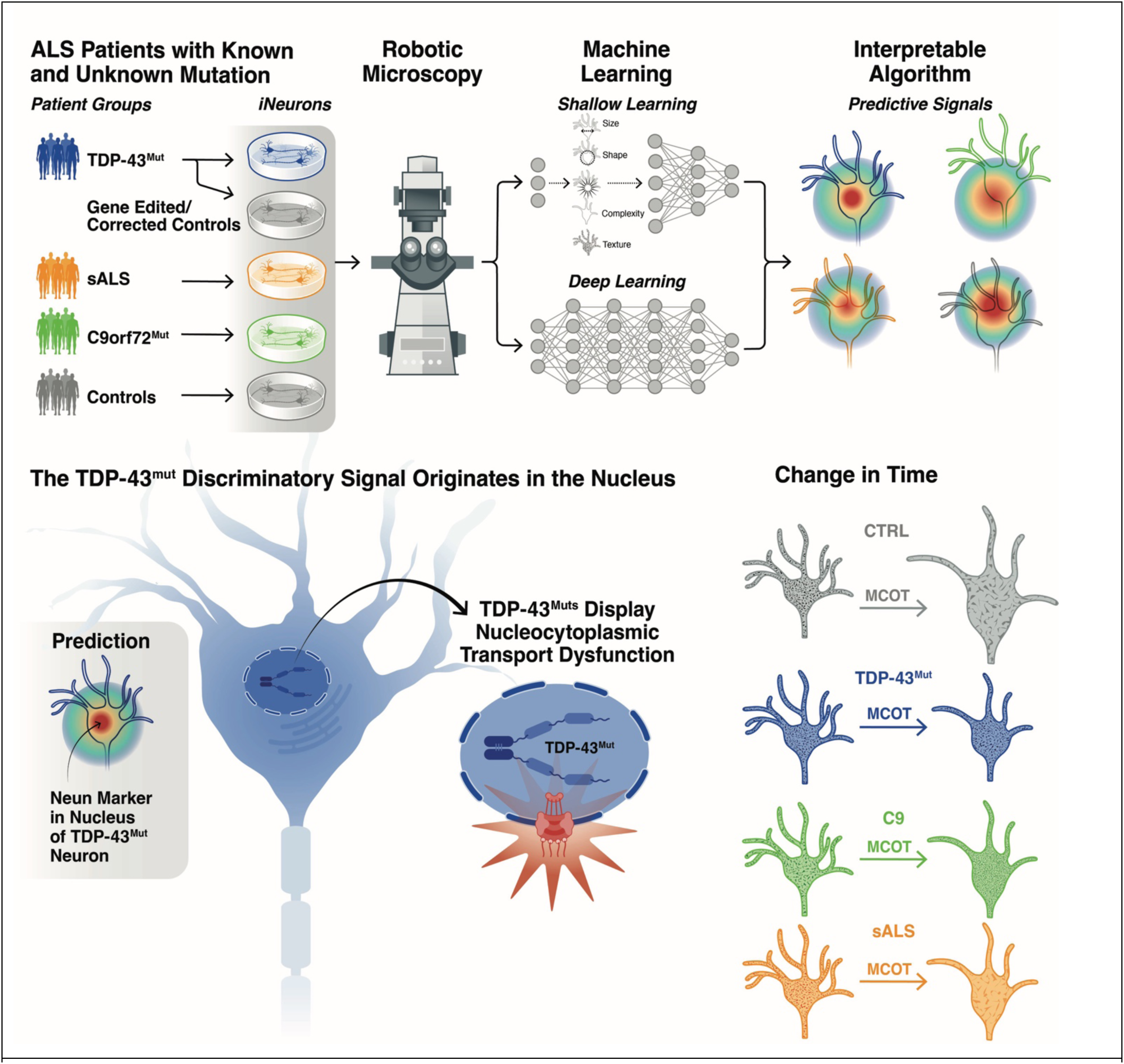
RM integrated with interpretable ML reveals predictive cellular signatures of ALS and provides a platform for mechanistic discovery across neurodegenerative and neurodevelopmental processes.

The TDP-43^mut^ iMNs demonstrated the most widespread decreases across multiple feature classes and displayed the largest number of changes that were consistent in the other ALS groups (**Figure 9B and Supplementary Figure 27, 28).** For example, the MCOT for features that capture “shape” such as eccentricity (e.g. round versus elliptical)^120^ decreased in the TDP-43^muts^ compared to controls, suggesting that the TDP-43^muts^ become rounder faster over time. This was not observed in C9orf72^muts^ or sALS compared to controls. As neurons die, they begin to round up and display fragmented structures^214–217^. The greater drop in eccentricity MCOT in TDP-43^muts^ relative to other ALS lines is consistent with their more robust neurodegeneration rate (**Figure 3**). Another “shape” feature is the “HOG” (histogram of gradients)^218^, which detects areas of significant intensity changes in the cell image that are indicative of the presence of edges or branching patterns. There was an extremely large difference between T1 and T6 in the HOG MCOT in the TDP-43^muts^ iMNs, whereas the controls were much more static (**Figure 8B and Supplementary Figure 27 D).** The C9orf72^muts^ showed a similar though less severe change in the HOG MCOT, whereas the sALS were no different than controls.

Other features that were most disrupted in TDP-43^muts^ compared to controls are features that capture complexity such as such as Minkowski-Bouligand Fractal Dimension (Mink Dim)^128^, which has been used to detect differences in complexity in human tissues^219^, and Corners and Voronoi clustering^220–222^, which has been used to capture information about neuronal structure^223–225^ (**Figure 8B and Supplementary Figure 27 F-J)**. Losses in complexity measures may reflect alterations in cytoskeletal organization and/or changes in microtubules or dendritic stability, all of which have been previously associated with TDP-43 pathology in ALS^226–228^. Interestingly, the only “complexity” MCOT that was significantly different in the C9orf72^muts^ was the “radial distance” MCOT, which had the largest difference of all features and was 20 times larger (2000% change) in C9orf72^muts^ than in controls (**Figure 8B and Supplementary Figure 27 E-J).** On the other hand, this MCOT was no different in the TDP-43^muts^ compared to controls and very slightly different in the sALS compared to their controls but in the opposite direction. The Mink Dim^128^ decreased across all ALS groups (**Supplementary Figure 28 F),** suggesting that a decrease in complexity over time is a unifying hallmark of ALS. Such decrease may be consistent with soma shrinkage and retraction, which are hallmarks of stressed or degenerating motor neurons in human brains and mouse models of ALS^229–231^. This is also consistent with the increased cell death observed across all groups relative to controls (**Figures 3, 7**). Although the direction of the MinkDim appears to differ across TDP-43^muts^ compared to controls (∼-230%) vs. C9orf72^muts^ and sALS compared to controls (∼+250%, +338% respectively) this case the change over time is in the control MCOTs and not the ALS MCOTs, which all decrease from T1 →T6 (**Figure 8B and Supplementary Figure 27 F).**

The “sholl”^232^ feature captures how many structures extend from the object in the middle of each crop and can be interpreted as a way to characterize the number of processes emanating from each neuron. Interestingly, we observe that for controls and ALS there is a sharp decline in the number of neurites emanating from each soma from T1 to T6. This decline is much faster in the TDP-43^muts^ and sALS iMNs (∼ +30% for both groups) compared to controls whereas it is similar in C9orf72^muts^ versus controls (**Figure 8B and Supplementary Figure 27 H).** The Voronoi MCOT differed across all groups but increased in both the C9orf72^muts^ and sALS iMNs compared to controls, with the largest change in the sALS (∼+420%) (**Figure 8B and Supplementary Figure 27 J).**

Other features capture information regarding the texture of objects in each crop. For instance, Fast Fourier transform-based texture descriptors use frequency domain to describe patterns in cells and animals^233–235^. Other texture features include Gray-Level Co-occurrence Matrix (GLCM)^236^, which measures the spatial relationships between neighboring pixel intensities while the Variance feature measures variance of the object/cell (**Figure 8B and Supplementary Figure 28 A-C**). The “texture” MCOT changed most significantly in the TDP-43^muts^ compared to controls, followed by C9orf72^muts^, and while the changes were in the same direction in sALS, they were much subtler (**Figure 8B and Supplementary Figure 28 A-C**). The last set of features captured information about the brightness of the iMNs within each crop such as “gaussian”, “luminance” and “peak max”. The largest changes were in C9orf72^muts^ versus controls followed by the TDP-43^muts^ with the least robust changes in the sALS (**Figure 8B and Supplementary Figure 28 D-F**).

Collectively, these analyses demonstrate that longitudinal morphological profiling captures robust, disease-specific phenotypic trajectories in ALS iMNs that may not be apparent from single time-point assessments. By quantifying the MCOT, we uncovered shared features of ALS, including reduced cellular growth and declining structural complexity, alongside mutation-specific patterns that distinguish TDP-43^muts^ C9orf72^muts^, and sALS. TDP-43^mut^ iMNs exhibited the most pronounced and widespread alterations across size, shape, complexity, texture, and neurite-associated features, consistent with their heightened neurodegenerative burden. In contrast, C9orf72^mut^ and sALS iMNs displayed more selective and subtler changes, with notable divergence in complexity and brightness-related features. Together, these results highlight MCOT as a human-interpretable sensitive framework for resolving dynamic disease-associated phenotypes, revealing both convergent and divergent cellular trajectories across ALS subtypes during neuronal maturation and degeneration.

## Discussion

Here we describe a novel approach to tackle a fundamental question in ALS: Is ALS a single disease entity with shared mechanisms across all etiologies or a collection of partially overlapping subtypes? Motor neuron degeneration is a universal hallmark of ALS, while TDP-43 proteinopathy occurs in almost all cases of ALS. As the vast majority of ALS cases are sporadic^2^, with no known cause, these shared hallmarks suggest that there must be common threads across cases. Some of these commonalties must be genetic as inferred from twin studies^237, 238^. The remaining unexplained risk suggests that non-genetic influences must also play an important role in ALS etiology. It’s currently unclear how to best unravel these common and disparate aspects of the disease and our study proposes a framework to begin disentangling these mechanisms (**Figure 9**).

We examined if simple cell-based hallmarks could distinguish ALS patients harboring known mutations such as *TARDBP* and *C9ORF72* mutations from each other and from ALS patients without known genetic mutations. To do this, we used high-content live-cell imaging of human iMNs and found that while ALS iMNs appeared to differentiate in a similar fashion to controls, they displayed spontaneous and natural neurodegeneration. The iMNs with *TARDBP* mutations died at the highest rate (**Figure 2, Supplementary Figure 8)**, followed by those with *C9ORF72* mutations. In sALS iMNs, the rate of cell death was variable across lines, but was overall higher than in controls (**Figure 6, Supplementary Figure 23)**. This suggests that at least *in vitro*, highly penetrant ALS-causing mutations display more robust degeneration than sALS lines which are likely influenced by combinations of low-penetrance risk loci. Hence, the fact that we have captured neurodegeneration as a penetrance-dependent gradient reflects both the biological fidelity and sensitivity of our cell imaging platform. Further, the fact that spontaneous neurodegeneration is a shared hallmark across all lines suggests a common thread in the disease and raises the possibility that familial and sporadic ALS converge on shared downstream mechanisms, despite distinct genetic architectures.

In addition, the difference in neurodegeneration rate mirrors some of the clinical features reported, though not perfectly. For example, ALS patients with *C9ORF72* expansions have been documented to display a more aggressive clinical course—e.g. shorter survival from onset—than patients with sALS^239^, which is somewhat consistent with our *in vitro* neurodegeneration data. However, C9ORF72 ALS is also more aggressive than ALS caused by *TARDBP* mutations^240, 241^, whereas in our cell lines, *TARDBP* mutations appeared more severe than *C9ORF72* mutations. In fact, much clinical variability has been reported among patient with *TARDBP* mutations^159, 242^ and in some carriers, disease progression is no different than in sALS^159^. On the other hand, there is some evidence that patients with *TARDBP* patients display an earlier onset but a slower progression compared to sALS^243^. The increased rates of neurodegeneration observed in our TDP-43^muts^ and C9orf72^muts^ may suggest that this model may capture cellular phenotypes that reflect aspects of clinical disease progression. However, these differences may also reflect other unknown factors, and it may be difficult to directly translate the behavior of isolated, immature neurons *in vitro* to the disease course observed *in vivo*. One important caveat is that our iMNs are monocultures, and they lack glial cells. Recent studies have shown that co-culture with glia is necessary and sufficient to drive C9orf72-mediated neurodegeneration^244–246^. Therefore, it’s possible that the aggressiveness of the TDP-43^mut^ phenotype in our system reflects an incomplete recapitulation of C9orf72 pathology, which depends on cell non-autonomous mechanisms and the interplay between loss- and gain-of-function effects that may require additional cell types to be fully expressed. Future studies should employ more sophisticated co-culture systems or organoid models to more faithfully recapitulate neurodegeneration.

ML on images from these lines reveals that underlying their neurodegeneration, there is a more complex pathological signal. We found that reproducible morphological patterns are present and apparent in ***live*** iMNs and sufficient to distinguish cells carrying known ALS-causing mutations from controls, with the strongest signal detected in TDP-43^muts^ lines (**Figures 3, 7, Supplementary Figures 12, 25, 26)**. The existence of morphology-based signals that render mutant cells visually distinguishable from healthy cells suggests that the ALS-associated signal is holistic rather than subtle or obscure. While these signals are not readily detectable by the human eye, they can be captured and quantified by ML. Further, that this signal is strongest in TDP-43^muts^ lines could mean that there is something unique to the pathology of these lines. Alternatively, the stronger signature in TDP-43^muts^ lines may simply reflect the use of gene-corrected or isogenic edited lines as controls, where reduced genetic variability enhances the detectable differences. Consistent with this interpretation, the sALS lines exhibited the weakest discriminative signal, which may reflect the greater genetic heterogeneity across these unrelated patient-derived lines.

Using the interpretable algorithm SmoothGrad in conjunction with immunohistochemistry allowed us to triangulate the predictive signal in TDP-43^muts^ iMNs to the nuclear/perinuclear region (**Figure 4, Supplementary Figure 15)**. This finding is consistent with previous reports that mutant TDP-43 disrupts nuclear pore complex integrity, leading to mislocalization of nucleoporins and defects in nuclear import and RNA export^247^. Further, assessment of NCT with a shuttling biosensor revealed nuclear reporter leakage or mislocalization in TDP-43^muts^ iMNs, consistent with impaired NCT fidelity (**Figure 5, Supplementary Figure 16**). These findings align with a growing body of evidence implicating nuclear pore complex dysfunction and NCT defects in TDP-43 proteinopathies^32, 247, 248^

### Capturing morphology changes over time is a sensitive approach to model ALS

Using a novel approach, MCOT, to quantify changes in cell size, shape or complexity between 2 timepoints, we uncovered robust shared and subtype-specific phenotypic trajectories (**Figure 8, Supplementary Figures 27, 28**). This suggested to us that incorporating time as a phenotyping dimension captures ALS-related phenotypes that emerge during maturation and degeneration and would not be captured by punctual observations. Interestingly, this analysis showed both overlapping and distinct features between TDP-43^muts^, C9orf72^muts^ and sALS. These findings support the idea that ALS may not represent a single uniform disorder, but rather a collection of partially overlapping disease subtypes with both common and distinct cellular manifestations.

In addition, by incorporating time as a parameter in our analyses, we may be able to capture both developmental and neurodegenerative differences as they unfold. For example, the ALS iMNs changed little in size between the two timepoints, whereas their control counterparts grew larger, suggesting a neurodevelopmental delay in the ALS cells (**Figure 8, 9, Supplementary Figure 27**). Interestingly, recent reports in C9orf72 expansion carriers suggest the presence of neurodevelopmental alterations such as changes in brain size in both white and grey matter well before the onset of ALS^85, 86^. Similarly, mouse models harboring the *C9ORF72* expansion display changes in thalamic size. These observations indicate that aspects of C9orf72-associated pathology arise during development rather than solely through late neurodegenerative processes^249^. Other reports with other ALS mutations report developmental delays in iPSC-derived spinal motor neurons^250^. Our findings that neuron growth is stunted not just in C9orf72^muts^ iMNs but also in TDP-43^muts^ and sALS iMNs suggest there also may be developmental alterations in other ALS forms, which warrants future investigations.

While stunted growth was observed in all mutant iMNs, complexity, another feature that could reflect development or neurodegeneration, declined in TDP-43^muts^ and sALS relative to controls over time but curiously not in C9orf72^muts^ (**Figure 8, 9, Supplementary Figure 27**). Reduced dendritic outgrowth and complexity has been reported in fly models of mutant TDP-43^251^ mouse models^252^ and in the brains of sALS patients^252^.

The MCOT metric also revealed subtype-specific patterns. For example, TDP-43^muts^ iMNs showed the most severe changes across the most feature classes, consistent with the observation that these lines displayed the largest amount of neurodegeneration. The shape features such as eccentricity shifted in directions consistent with neuronal rounding, which is a hallmark of cellular injury and impending death. Shape and edge-based measures such as HOG also diverged strongly from controls, suggesting altered boundary structure or process organization. Curiously, C9orf72^muts^ and sALS iMNs exhibited more selective changes, such as alterations in a subset of complexity features as well as intensity or brightness.

Together, these longitudinal results suggest that ALS iMNs are not merely distinguishable by static morphological differences, but by altered developmental and degenerative trajectories. MCOT provides a sensitive framework for identifying early phenotypes that precede overt cell death and for pinpointing which aspects of neuronal growth, structural organization, and intracellular texture diverge first across disease subtypes.

### SML versus DNN performance

Both SML and DNNs distinguished TDP-43^muts^ from control simply from live EGFP signal, but the ResNet DNNs outperformed the SML models (**Figure 3, Supplementary Figure 12**). Label-scrambling abolished performance across models and time points, arguing that the classifiers leveraged biologically meaningful structure rather than experimental artifacts or dataset leakage. ResNet DNNs also outperformed SML models for C9orf72 (**Supplementary Figure 25**) and sALS (**Supplementary 26**) models. Likewise, when scrambling labels for C9Orf72 we also observe chance-level performance for both T1 and T6 (**Supplementary 25 B**), and for sALS (**Supplementary 26 B**). Interestingly, classification slightly improved at the later time point, most notably for TDP-43^mut^ iMNs and to a lesser degree with C9orf72^mut^ iMNs. This may suggest that disease-related phenotypes become more apparent as iMNs mature. This is consistent with recent reports that revealed that as iMNs mature, there is a progressive emergence of TDP-43–associated molecular defects^213^. Likewise, previous reports show that the C9orf72 pathology becomes more apparent as neurons mature^202^. The signal distinguishing sALS iMNs from controls was consistently more modest, even with the DNNs, nevertheless, a clear ALS-discriminating pattern is detectable. The fact that the performance of both the SML and DNN remained above chance and was eliminated by scrambling proves that detectable, genuine signals exist in these datasets, but they may require larger cohorts, higher-resolution images or the use of different cellular structures and/or other types of stratification to resolve.

The comparatively lower performance of the shallow connected ML algorithms model (**Figure 3B-C, Supplementary Figure 12,25, 26**) relative to the DNNs can be attributed to limitations in feature representation (the model’s ability to capture meaningful patterns from raw data) as well as inductive bias (the built-in assumptions that a model makes about how to generalize from training examples). The stacked classifier framework relies on the sensitivity of the engineered features. In single-cell image classification, discriminative cues can be subtle, high-dimensional, and spatially structured. Given that several of the engineered features, predominantly those dealing with shape, rely on the image being flattened, overall there may be an effect where these features are unable to capture complex patterns and spatial hierarchies that CNNs learn end-to-end. In contrast, both ResNet architectures benefit from hierarchical feature extraction tailored to image data, allowing them to capture patterns ranging from low-level edges to higher-level cellular morphology. ResNet50 models have additional depth and representational capacity that may allow it to learn subtle differences between cell types than a shallower model like our Stacked Classifier model can derive. This applies for our C9orf72, sALS, and TDP-43 models.

The superior performance of the ImageNet-pretrained ResNet18 (**Figure 3D-E, Supplementary Figure 12**) over the SimCLR-pretrained ResNet50 for the TDP-43 model, suggests that transfer learning from large, labeled natural image datasets can provide more task-relevant inductive priors than self-supervised representations learned on a potentially mismatched or limited domain. We did experiment with a ResNet18 backbone pretrained using SimCLR, and this failed to surpass the performance of the fully supervised ImageNet-pretrained ResNet18. This may reinforce the importance of large-scale supervised pretraining in this setting. In addition, the smaller ResNet18 model may generalize better under limited training data, as the ResNet50 is known to be higher capacity and so more prone to overfitting when fine-tuned on relatively small single-cell datasets.

Several strategies could enable performance improvements beyond the current best model. For the SML model, incorporating deep learned features (such as embeddings extracted from a pretrained CNN) into the stacking pipeline could potentially enhance representational power. For the ResNet50, performance may be improved through larger scale domain-adaptive pretraining, stronger regularization, and optimized fine-tuning strategies such as layer-wise learning rate decay (which prevents early layers from being updated too aggressively and overwriting useful low-level features) or selective layer freezing (which retains pretrained representations in earlier layers while allowing task-specific adaptation in deeper ones). Although the ImageNet-pretrained ResNet18 currently performs best, it could potentially be surpassed by architectures that combine its strong transfer learning initialization with task-specific enhancements, including multi-scale feature fusion (where features extracted at different depths of the network are combined, allowing the model to simultaneously leverage fine-grained local textures and broader structural patterns) or ensemble approaches. These approaches aim to better align model capacity and learned representations with the characteristics of single-cell imagery, which theoretically would lead to gains in classification accuracy.

The image filtering process was designed to isolate high quality single-cell crops while preserving as much usable data as possible. Since the goal was to analyze individual cells rather than whole fields of view, we experimented with a range of filtering parameters, including shape constraints, noise metrics, etc. However, overly stringent criteria substantially reduced the dataset, and ultimately we adopted one minimal, but biologically meaningful constraint, which as previously mentioned in the Methods section involves excluding any cell that touched the crop boundaries. Cells that intersect the image edge are often partially captured, which can distort downstream feature extraction. Therefore, with this rule we ensured that each retained image contained a complete cell while maintaining maximal sample diversity and statistical power.

Going forward, we aim to replace manual/rule-based filtering with a supervised machine learning model capable of automatically distinguishing high-quality single-cell images from artifacts or suboptimal crops. This model could theoretically learn nuanced quality features, such as focus sharpness, segmentation accuracy, signal-to-noise ratio, and morphological plausibility, without relying on rigid thresholds. Such an approach could improve scalability, reduce subjective bias, and allow more consistent quality control across large imaging datasets.

### Implications for ALS Therapeutic Discovery

Increasingly, clinical trials are failing for lack of efficacy rather than safety, and that raises questions about the preclinical approach we are using to find targets and drugs. Perhaps the approach is too narrow and fixing a single phenotypic abnormality may not be enough to produce an efficacy signal in a clinical trial that is sufficient to reach an approvable endpoint^253, 254^. If ALS is a disease of a complex system, then we may need to understand it and treat it as a system rather than to search for a single target. Further, by harnessing the power of ML to look wholistically at a system and to identify its distinctive features, we might be able to redefine what an ALS phenotype is. Further, by studying the process in cells derived from patients, we can capture mechanisms that might emanate from multiple genetic foci and converge on intracellular targets and pathways, some of which are not commonly measured. Some have suggested that an approach like this could lead to an exciting redefinition of neurodegenerative diseases based on pathway dysfunction rather than clinical profile^255^. If so, then we could have a new way to screen for drugs that uses this multidimensional definition of ALS in latent space to search for drugs (rather than single endpoint assays) that make a sick cell more fully healthy and might have a greater chance of being effective enough in clinic to become an approved drug.

Our work illustrates a practical workflow for discovering disease mechanisms in ALS and for uncovering both shared and distinguishing signals across subtypes. By using ML to detect otherwise subtle phenotypes in live cells, and combining interpretability algorithms and staining to localize the signals, we can generate hypothesis-driven, target-specific assays to test the underlying biology. This framework may also be broadly applicable to other diseases in future studies.

## Acknowledgements

We would like to acknowledge these sources of funding that supported this work: Genentech, CIRM (DISC0-13914), Answer ALS, ALS FindingACure, Target ALS IL-2023-C4-L2, U.S. Army Medical Research Acquisition Activity (USAMRAA), W81XWH-22-1-0721, CIRM (DISC0-16039), P01AG073082 and the ALS Association, NIH R01 LM 013617, the Robert Packard Center for ALS Research. We thank Matthew Harms for confirming the C9orf72 repeat expansion in one of the lines. We also thank Françoise Chanut for editorial assistance, Tami Tolpa for scientific illustrations, and Kelley Nelson and Gayane Abramova for assistance with EndNote formatting and many of other aspects of this work.

## Author contributions

Conceptualization, J.A.K, A.J, S.M.F, Methodology, N.A, U.C, N.B, Z.F, M.A, R.T, M.B, D.L, J.A.K, Software, N.A, M.A, R.T, M.B, D.L, Formal Analysis, J.A.K, N.A, U.C, Z.F, M.A, R.T, W.R, E.V, K.R, D.L, Investigation, J.A.K, N.A, U.C, Z.F, M.A, R.T, D.L, Resources, J.A.K, A.J, SMF, Data Curation, N.A, U.C, Z.F, W.R, E.V, K.R,, Writing – Original Draft, J.A.K, N.A; Writing – Review & Editing, J.A.K, A.J, D.L, S.M.F. Visualization J.A.K, N.A, R.T Supervision, J.A.K, SMF. Project Administration, J.A.K, Funding Acquisition, J.A.K, S.M.F, A.J

## Notes

### Competing Interest Statement

The authors have declared no competing interest.

