## Supplementary Figures and Tables for "Predictive Cellular Signatures from Live Human Motor Neurons Distinguish TDP-43 ALS and Enable ALS Subtype Stratification": Kaye et al_Supplementary Figures and Tables_Bioarchives_C.pdf

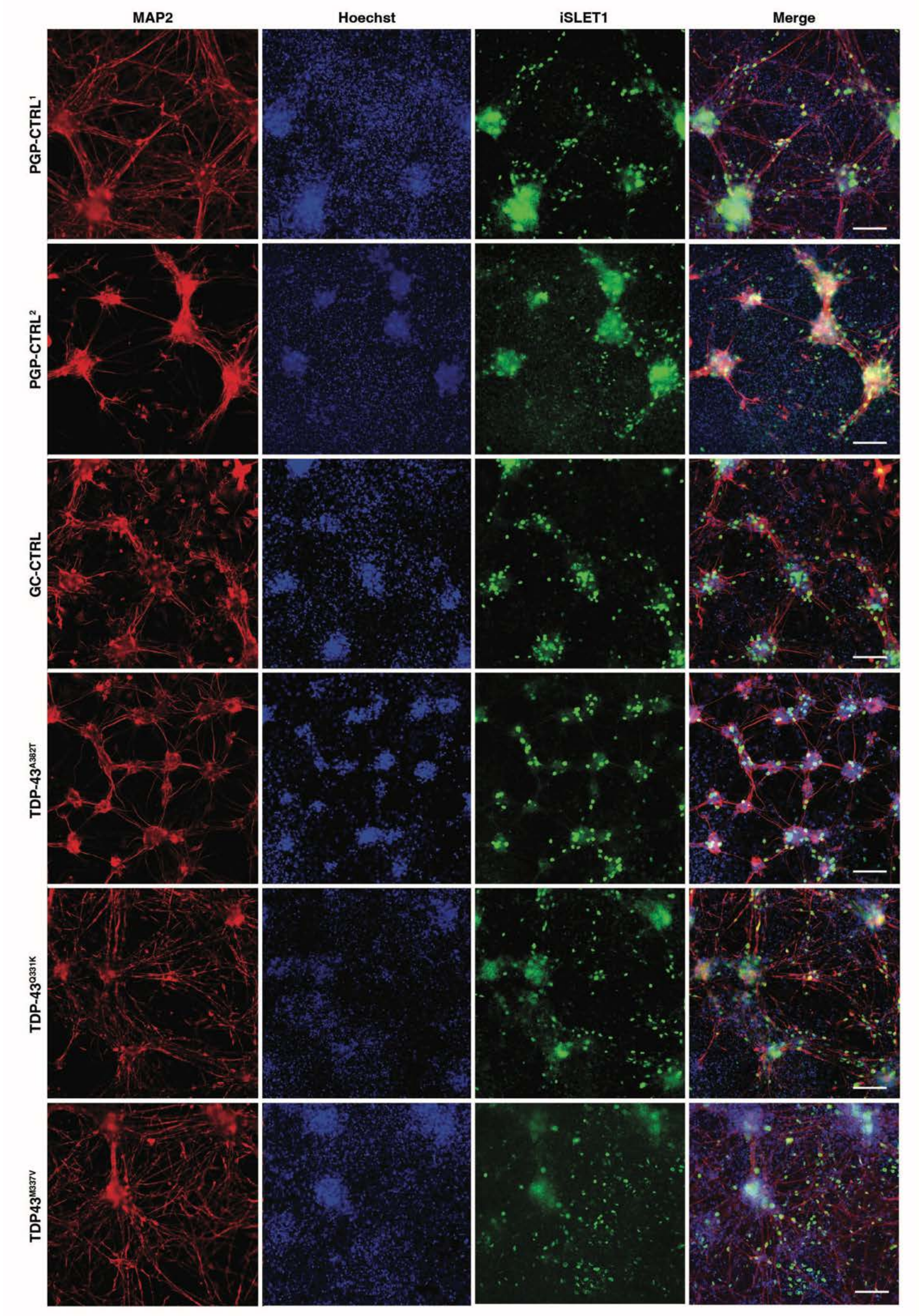


**Supplementary Figure 1. Expression of the motor neuron marker ISLET1 in TDP-43$^{mut}$ and control iMNs.** Representative images of isogenic control and TDP-43$^{mut}$ iMNs stained for the motor neuron marker ISLET1. At differentiation day ~ DIV 35, iMNs were fixed and stained using antibodies against MAP-2 (red), ISLET1 (green) and DAPI (blue). Merged image is on the far right, scale bar = ~90 uM.

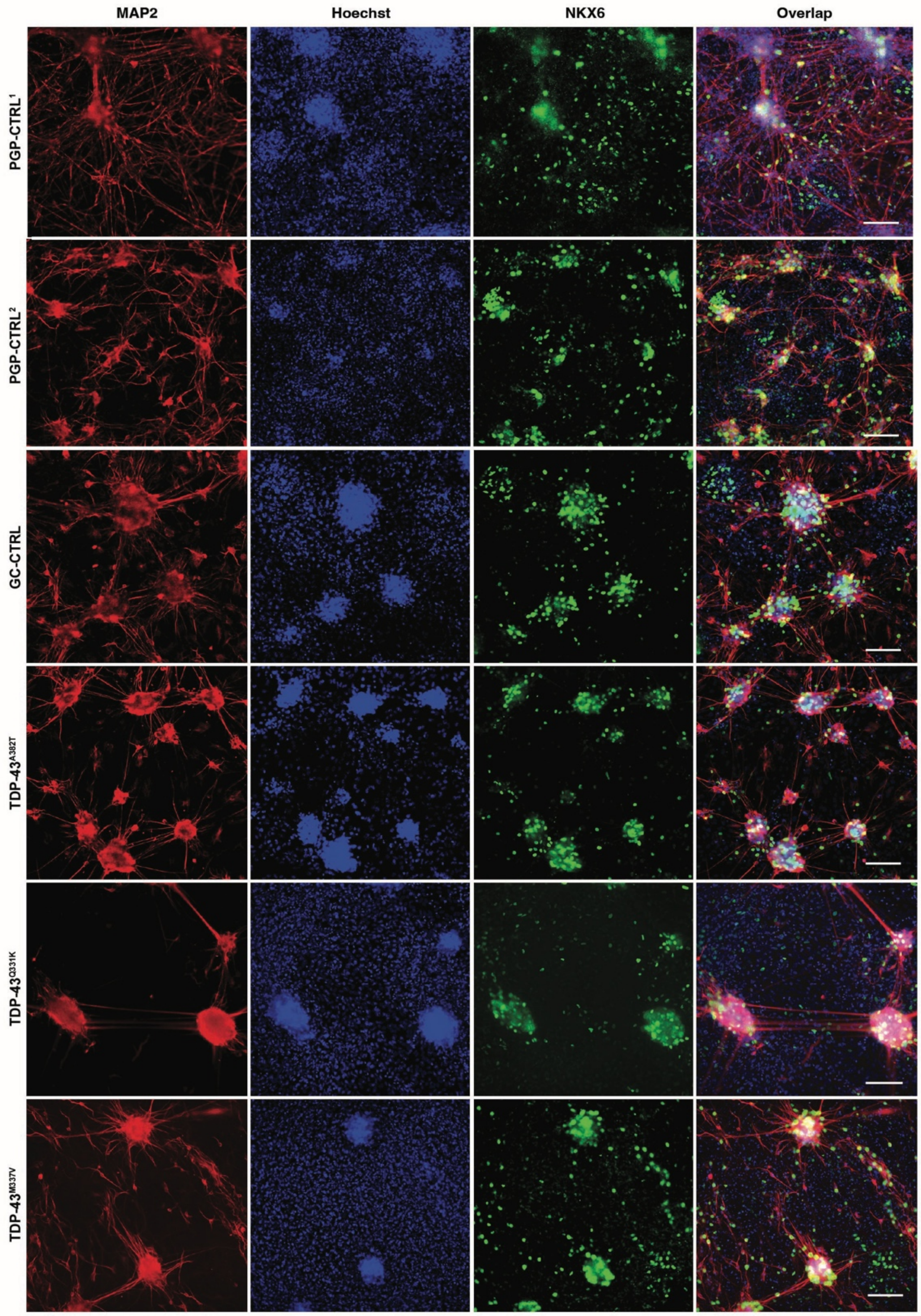


**Supplementary Figure 2. Expression of motor neuron marker NKX6.2 in TDP-43$^{mut}$ and control iMNs.** Representative images of isogenic control and TDP-43$^{mut}$ iMNs stained for the motor neuron marker NKX6.2. At differentiation day ~ DIV 35, iMNs were fixed and stained using antibodies against MAP-2 (red), NKX6.2 (green), and DAPI (blue). Merged image is on the far right, scale bar = ~90 uM.

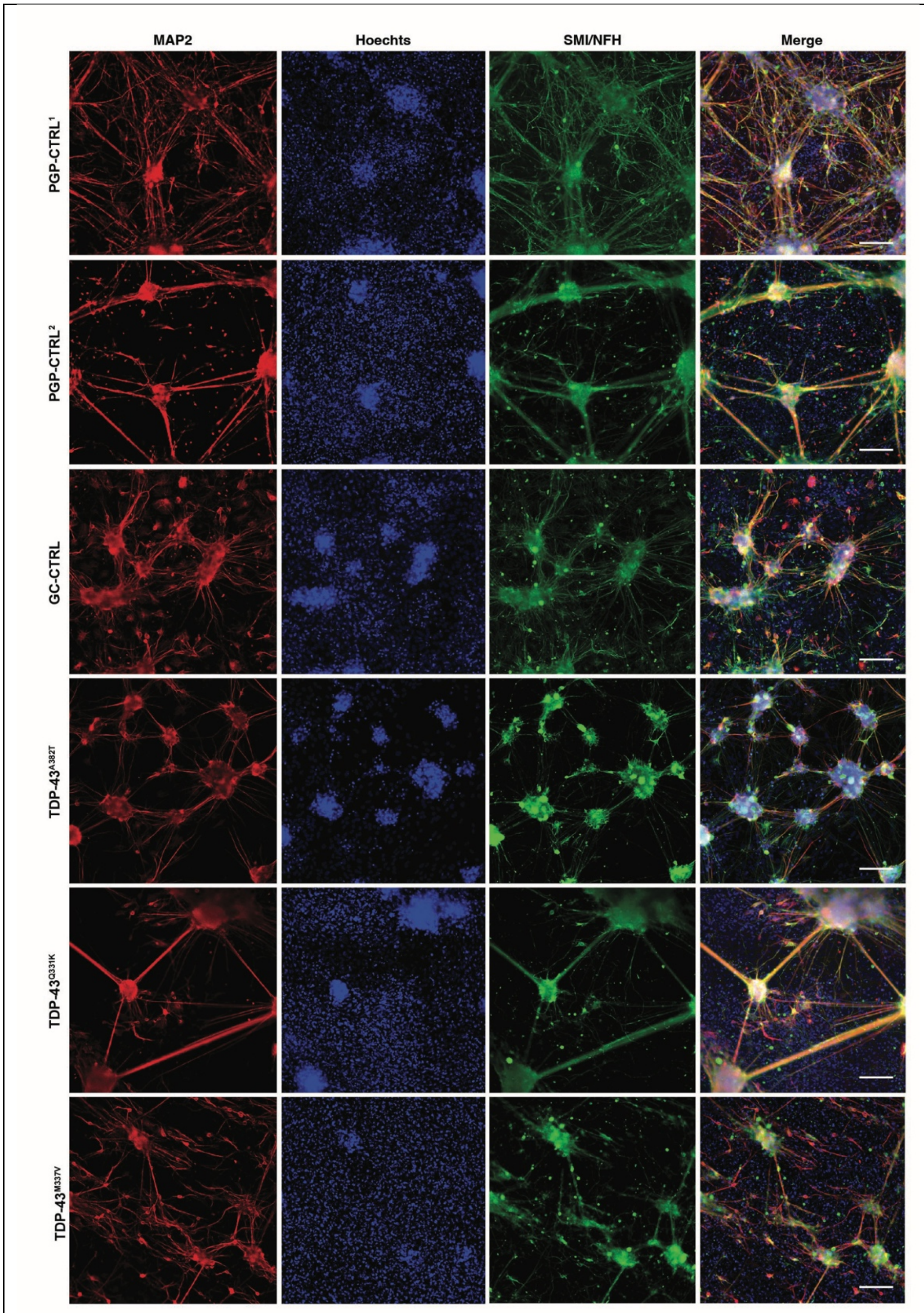


**Supplementary Figure 3. Expression of neurofilament H in TDP-43$^{mut}$ and control iMNs.** Representative images of isogenic control and TDP-43$^{mut}$ iMNs stained for the neurofilament H. At differentiation day ~ DIV 35, iMNs were fixed and stained using antibodies against MAP-2 (red), SMI/NFH (green), and DAPI (blue). Merged image is on the far right, scale bar = ~90 uM.

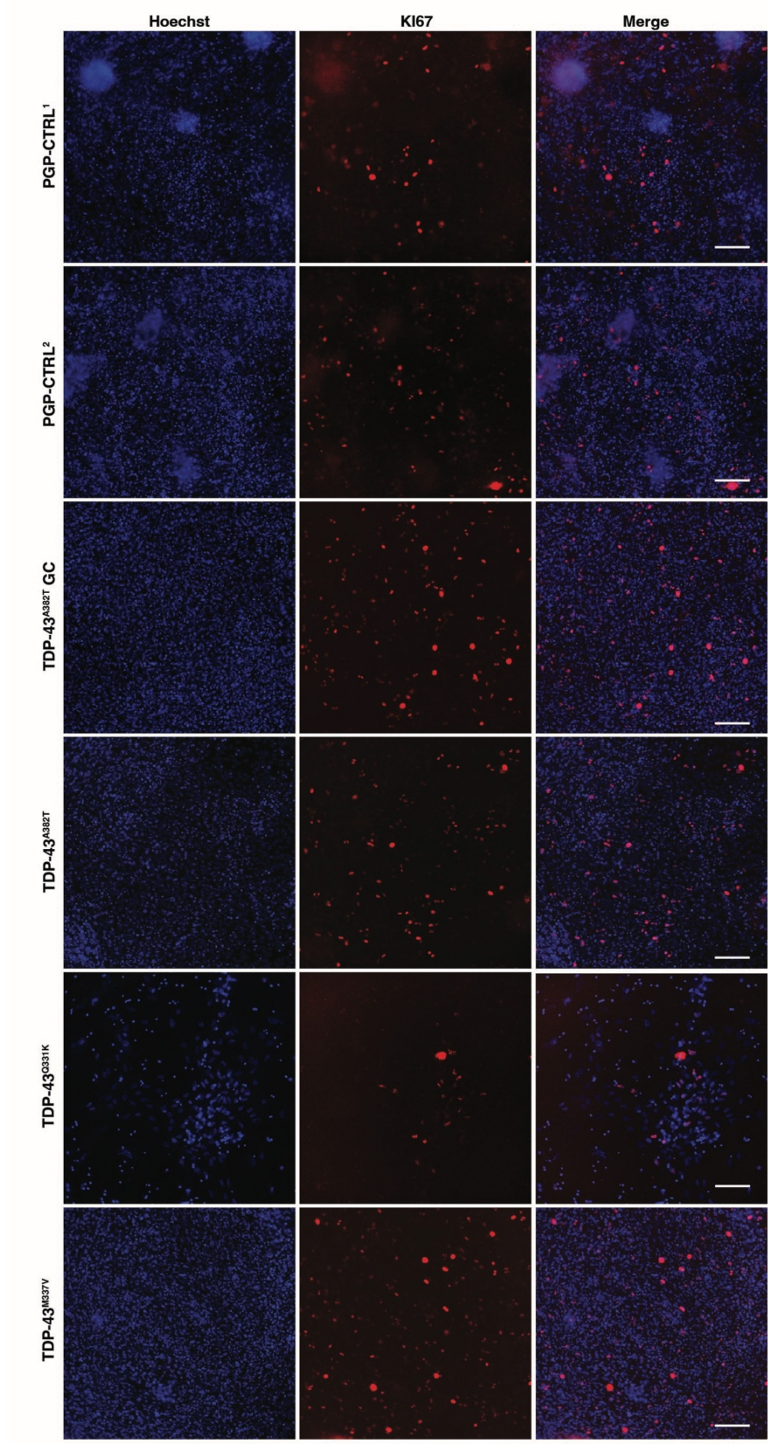


**Supplementary Figure 4. Expression of KI67 in TDP-43$^{mut}$ and control iMNs.** Representative images of isogenic control and TDP-43$^{mut}$ iMNs stained for the proliferation marker KI67. At differentiation day ~ DIV 35, iMNs were fixed and stained using antibodies against KI67 (red) and DAPI (blue). Merged image is on the far-right, scale bar = 100 uM.

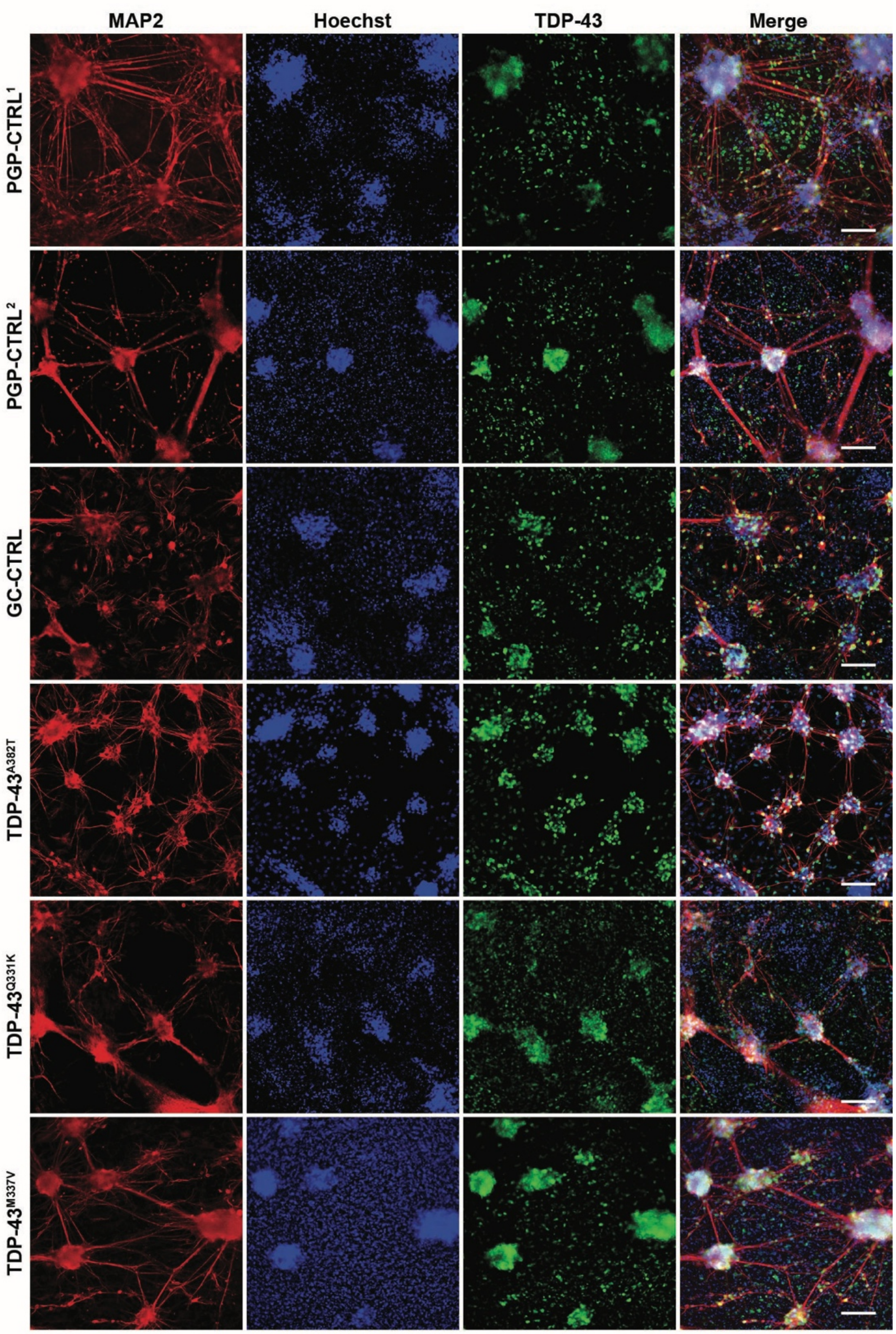


**Supplementary Figure 5. Expression of TDP-43 in TDP-43$^{mut}$ and control iMNs.** Representative images of isogenic control and TDP-43$^{mut}$ iMNs stained for TDP-43. At differentiation day ~ DIV 35, iMNs were fixed and stained using antibodies against MAP-2 (red), TDP-43 (green), and DAPI (blue). Merged image is on the far right, scale bar = ~90 uM.

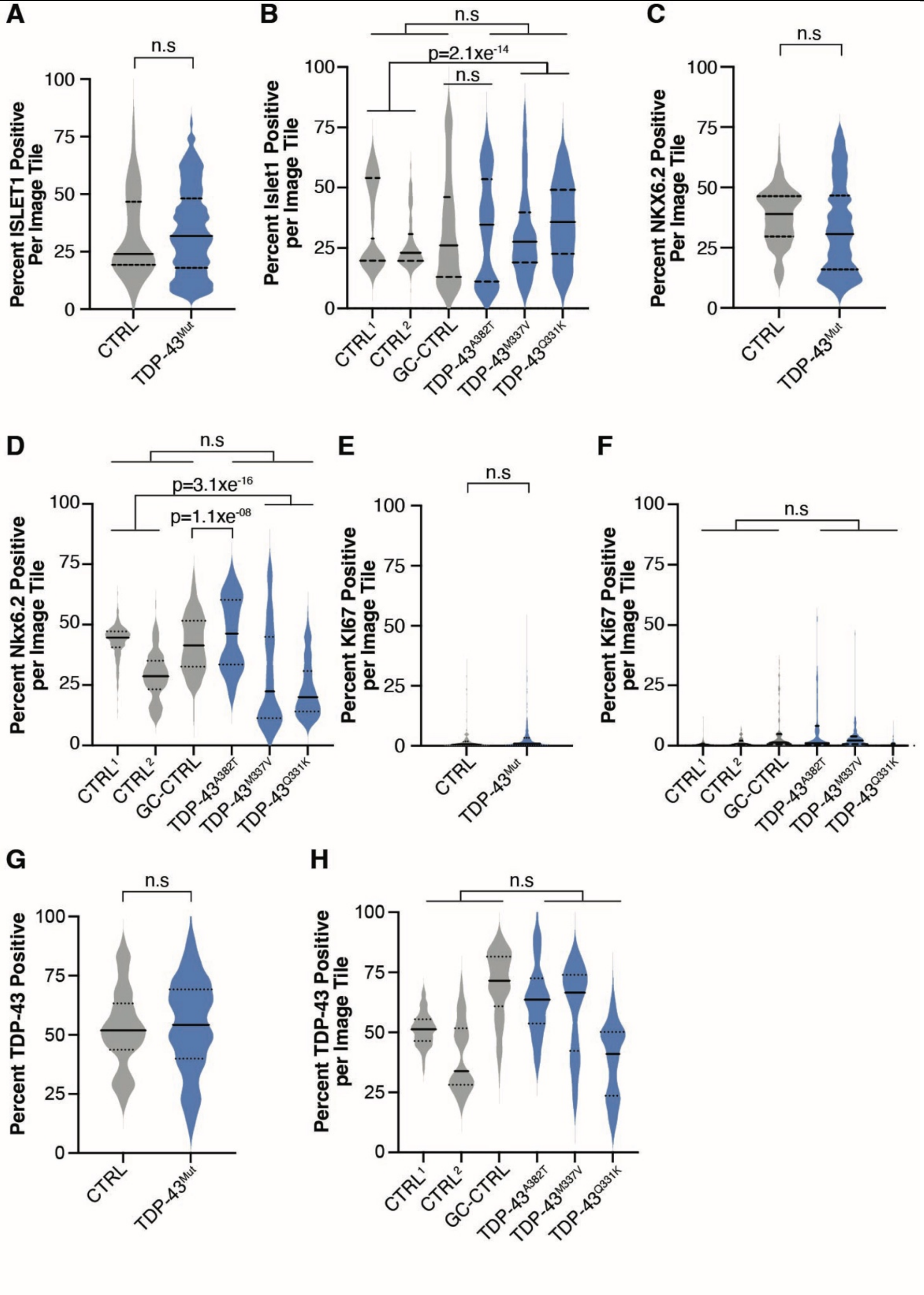


**Supplementary Figure 6. TDP-43$^{mut}$ and control iPSC lines produce comparable amounts of ISLET1, NKX6.2-, KI67 and TDP-43-positive iMNs.** Violin plots showing the proportion of cells staining for indicated markers relative to DAPI-stained cells across individual tiles. Images of isogenic control and TDP-43$^{mut}$ iMNs were measured using Cell Profiler[84, 97]. In all stains, n = 3–4 experiments per cell line. The best fit model for all data was a Poisson distribution, so estimates and p values were generated using a Poisson GLMM[110] with post-hoc Benjamini Hochberg[256] multiple-comparison correction. Some variability is observed across all cell lines but differences between all TDP-43$^{mut}$ and control cultures are not significant. **A**) There is no difference in the proportion of ISLET1-expressing cells between control and TDP-43$^{mut}$ iMNs, and tiles contained on average 33% ISLET1-expressing cells in mutant and control lines. n= ~8 independent experimental replicates, 3-4 experiments per cell line, 531 image tiles for controls and 441 image tiles for controls and TDP-43$^{mut}$, 1,197,412 cells. (estimate= -0.23, p= 0.45). **B**) Violin plots showing each TDP-43$^{mut}$ and control line. When comparing the CTRL$^{1-2}$ lines to the gene-edited TDP-43$^{M337V/Q331K}$ lines, there is a significant increase in the number of ISLET1 – positive iMNs (estimate= -0.47, p= 2.1xE$^{-14}$). There is no difference in the proportion of ISLET1- positive cells in the TDP-43$^{A382T}$ line compared to its gene- corrected control, (estimate= -0.04, p= 0.6). However, when comparing the CTRL$^{1-2}$ lines to the gene- edited TDP-43$^{M337V/Q331K}$ lines, there is a significant decrease in the number of ISLET1 – positive iMNs (estimate= -0.54, p= 3.1xE$^{-16}$). TDP-43$^{M337V}$ n = 126,630 cells, TDP-43$^{Q331K}$ n = 230,111 cells, TDP-43$^{A382T}$ n = 134,858 cells, PGP-CTRL$^{1}$ n = 238,895 cells, PGP-CTRL$^{2}$ n = 278,502 cells, GC-CTRL$^{1}$ n= 188,416 cells. **C**) The number of NKX6.2-expressing cells is similar in all TDP-43$^{mut}$ iMNs and controls (~34%). n = 8 separate experiments, 3-4 experiments per cell line, 531 tiles for controls and 426 tiles for TDP-43$^{mut}$ 1,199,587 cells. (Estimate= -0.17, p=0.6). **D**) However, when comparing the CTRL$^{1-2}$ lines to the gene- edited TDP-43$^{M337V/Q331K}$ lines, there is a significant decrease in the number of NKX6.2 – positive iMNs (estimate= -0.54, p= 3.1xE$^{-16}$) while there is a significant increase in the in the number of TDP-43$^{A382T}$ positive iMNs compared to its gene-corrected control (Estimate= 0.3, p= 3.1xE$^{-08}$). TDP-43$^{M337V}$ n = 126,630 cells, TDP-43$^{Q331K}$ n = 230,111 cells, TDP-43$^{A382T}$ n = 134,858 cells, PGP-CTRL$^{1}$ n = 238,895 cells, PGP-CTRL$^{2}$ n = 278,502 cells, GC-CTRL$^{1}$ n= 188,416 cells. **E**) There is no difference in the number of KI67 expressing cells for control and TDP-43$^{mut}$ iMNs. The number of proliferating KI67 positive cells is equally low (~ 3% overall) in control and TDP-43$^{mut}$ iMNs (n = 8 separate experiments, 3-4 experiments per cell line, 424 image tiles for controls and 425 image tiles for TDP-43$^{mut}$, 877, 602 cells. (Estimate = 0.17, p= 0.81). **F**) Violin Plots showing each line. TDP-43$^{M337V}$ n = 104,455 cells, TDP-43$^{Q331K}$ n = 139,651 cells, TDP-43$^{A382T}$ n = 121,45 cells, PGP-CTRL$^{1}$ n = 323,447 cells, PGP-CTRL$^{2}$ n = 347,467 cells, GC-CTRL$^{1}$ n = 163,113 cells. **G**) There is no difference in the number of TDP-43 expressing cells for control and TDP-43$^{mut}$ iMNs. (Estimate = -0.19, p= 0.59). n=9 independent experimental replicates, 3-4 experiments per cell line, 522 image tiles for controls and 408 image tiles TDP-43$^{mut}$ iMNs = 520,392 cells and TDP-43$^{mut}$, 739,798 cells. **F**) Violin Plots showing each line. When comparing the CTRL$^{1-2}$ lines to the gene- edited TDP-43$^{M337V/Q331K}$ lines, there is no significant decrease in the number of TDP-43 – positive iMNs (estimate= -0.05, p= 0.83) while there is a significant decrease in the in the number of TDP-43$^{A382T}$ positive iMNs compared to its gene- corrected control (Estimate= -0.14, p= 3.4xE$^{-14}$). TDP-43$^{M337V}$ n = 155,722 cells, TDP-43$^{Q331K}$ n = 233,655 cells, TDP-43$^{A382T}$ n = 131,015 cells, PGP-CTRL$^{1}$ n = 316,057 cells, PGP-CTRL$^{2}$ n = 229,565 cells, GC-CTRL$^{1}$ n = 194,176 cells.

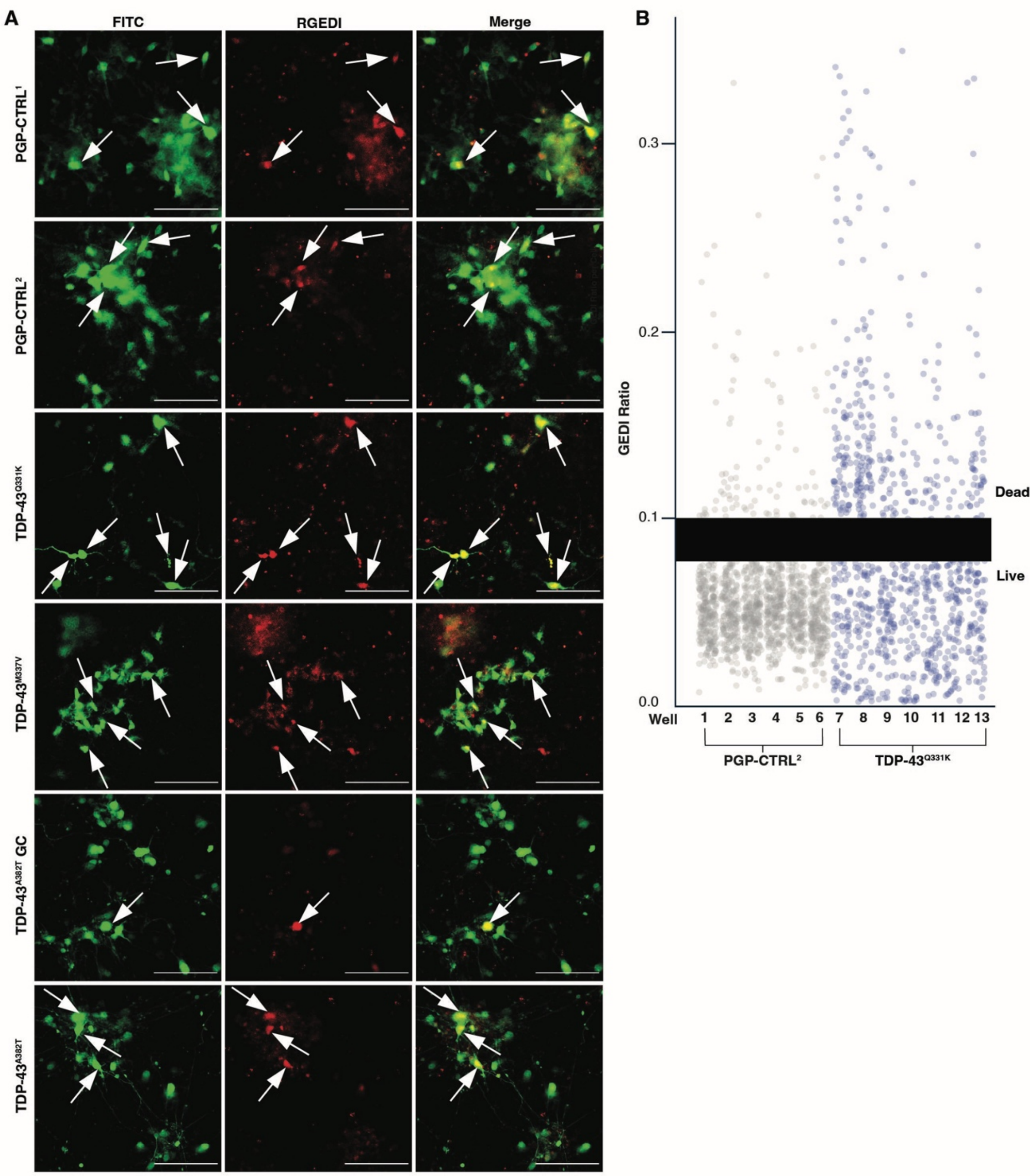


**Supplementary Figure 7. Using the GEDI biosensor to measure the rate of cell death in the TDP-43$^{mut}$ and control iMNs. A**) Representative images TDP-43$^{mut}$ and isogenic control iMNs expressing GEDI on ~DIV 30. Left panels display the GFP morphology marker of GEDI, the center panels the RGEDI cell death marker, and the right panels the merged images. The arrows indicate dead or dying cells within the frame. Scale bar = 200 µm. **B**) Example plot showing the distribution of GEDI ratios (RGEDI signal/GFP signal) in parallel cultures of TDP-43$^{Q331K}$ and PGP-CTRL$^{2}$ iMNs. Each dot represents a single cell. Cells with GEDI ratios below 0.08 are considered “alive” whereas cells with GEDI ratios above 0.1 are classified as “dead”. We exclude cells that are between 0.08 and 1.0. For each experiment, we determine the “live” and “dead” based on the segmentation and/ or treating cells with sodium arsenate to induce death (not shown). The numbers at the bottom represent the well from which the GEDI values were obtained.

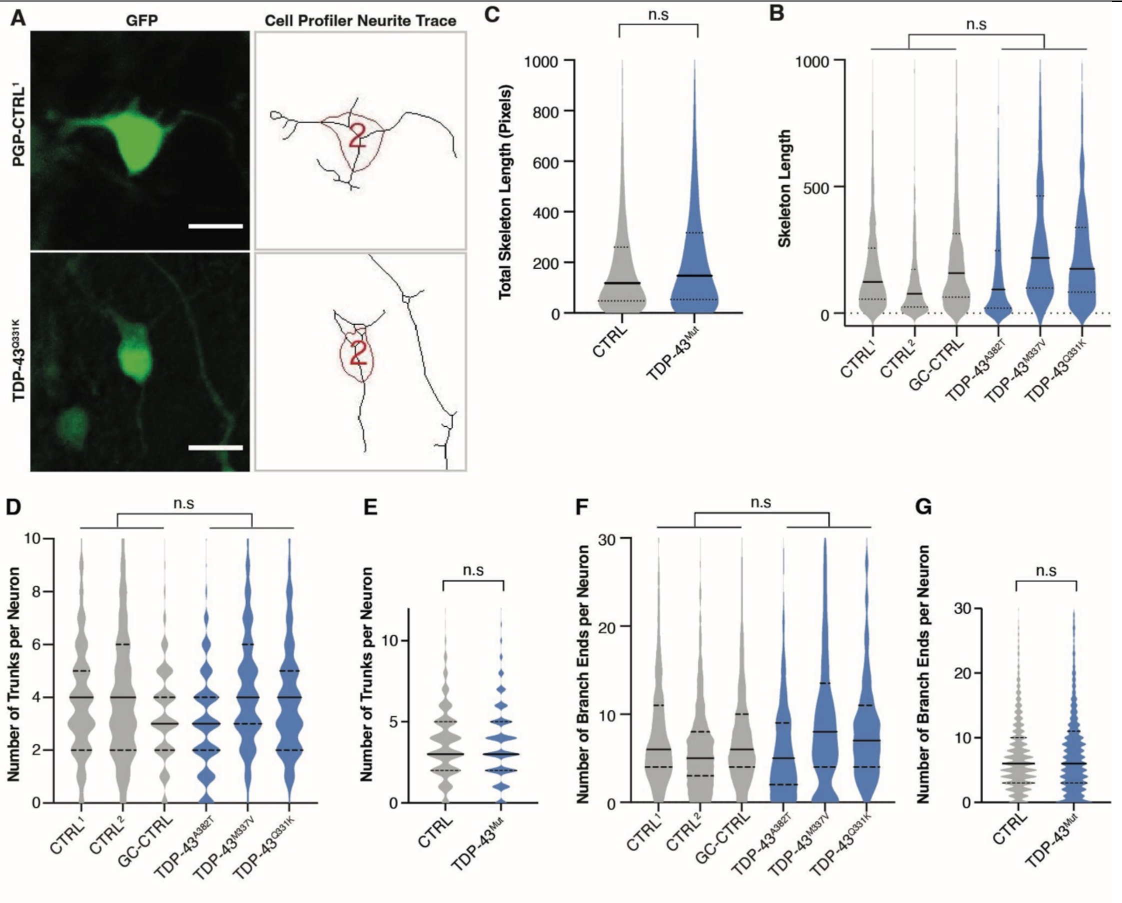


**Supplementary Figure 8. Neurites from TDP-43$^{mut}$ and i control iMNs have comparable lengths and branching patterns. A**) GFP images of DIV 32 neurons expressing GEDI were subjected to a neurite-segmenting and tracing pipeline as previously described[84, 97, 100] to capture neurite length and branching outlines. Images on the right show the trace corresponding to the GFP image on the left. **B**) Violin plots of all TDP-43$^{mut}$ and control iMNs neurite length shows no significant difference. Neurite length (estimate= 0.25, p = 0.35), **C**) Violin plot showing combined TDP-43$^{mut}$ and controls. **D**) Violin plots of all TDP-43$^{mut}$ and control iMNs for number of trunks shows no significant difference. Number of trunks (estimate = -0.02, p = 0.96) **E**) Violin plot showing combined TDP-43$^{mut}$ and controls for number of trunks. **F)** Violin plots of all TDP-43$^{mut}$ and control iMNs for number of neurite branches (estimate = 0.56, p = 0.49). **G**) Combined Violin Plots. n = 4 experiments, TDP-43$^{M337V}$ n = 705 cells, TDP-43$^{Q331K}$ n = 591 cells, TDP-43$^{A382T}$ n = 1,245 cells, PGP-CTRL$^{1}$ n = 929 cells, PGP-CTRL$^{2}$ n =1,093 cells, GC-CTRL$^{1}$ n = 1,654 cells. The best fit model for all data was a Poisson distribution so estimates and p values were generated using a Poisson GLMM[110] with post-hoc Benjamini Hochberg[256] multiple-comparison correction.

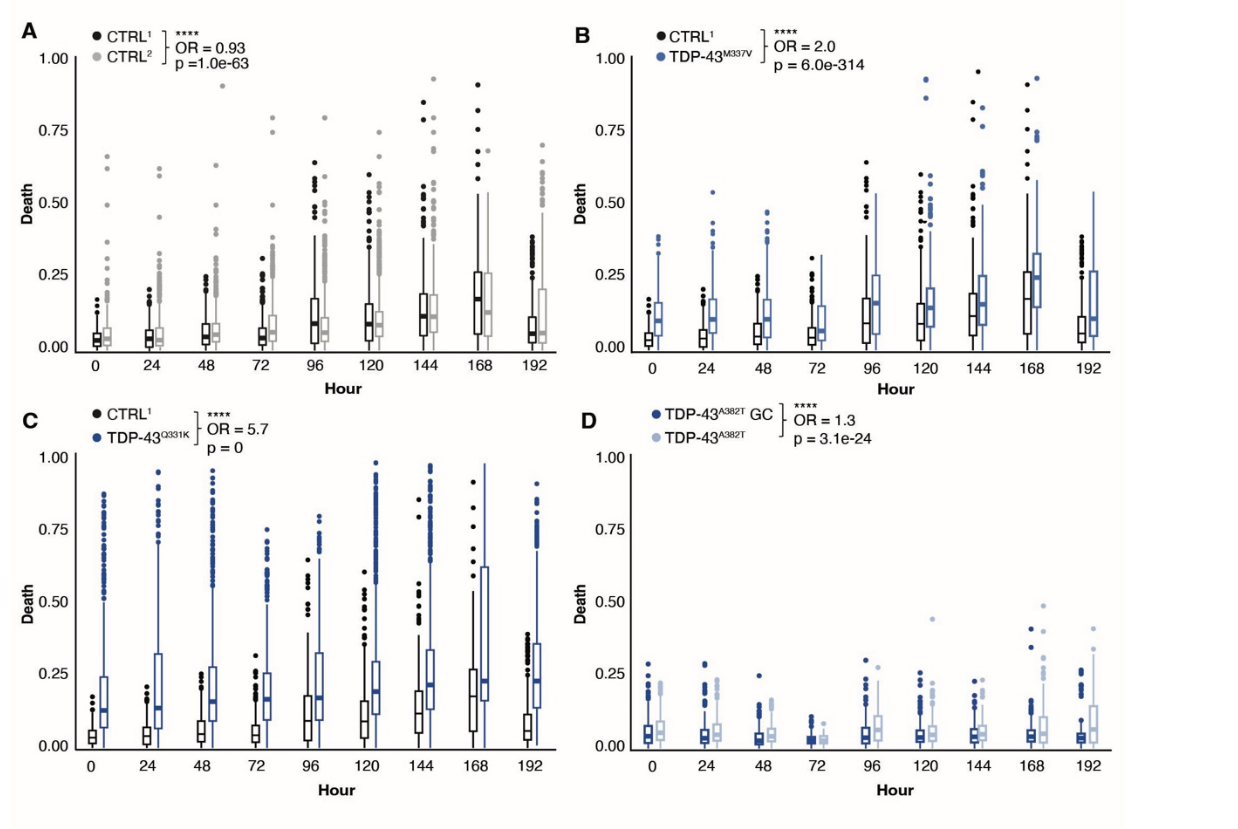


**Supplementary Figure 9. Longitudinal quantification of cell death in TDP-43$^{mut}$ compared to control iMNs using GEDI.** Plots show the proportion of dead cells in each well over time, quantified from GEDI ratios measured every 24 hours for up to 192 hours. Boxplots display the median and interquartile range; whiskers represent data range. Individual dots show outlier wells. **A**) CTRL$^1$ iMNs survive slightly better than CTRL$^2$ iMNs. OR-CD = 0.93, p = 1.0e–63. **B**) TDP-43$^{M337V}$ iMNs exhibit a higher rate of death than their isogenic control CTRL$^1$. OR-CD = 2.0, p = 6.0e$^{-314}$. **C**) TDP-43$^{Q331K}$ iMNs show a pronounced increase in cell death compared to isogenic controls. OR-CD = 5.7, p = 0. **D**) TDP-43$^{A382T}$ iMNs have a moderately higher death rate than their GC controls (OR-CD = 1.3, p = 3.1e$^{-24}$). TDP-43$^{M337V}$ n = 328,909 cells, TDP-43$^{Q331K}$ n = 335, 298 cells, TDP-43$^{A382T}$ n = 196,051cells, PGP-CTRL$^1$ n = 380, 463 cells, PGP-CTRL$^2$ n = 445, 609 cells, GC-CTRL$^1$ n =245, 028 cells.

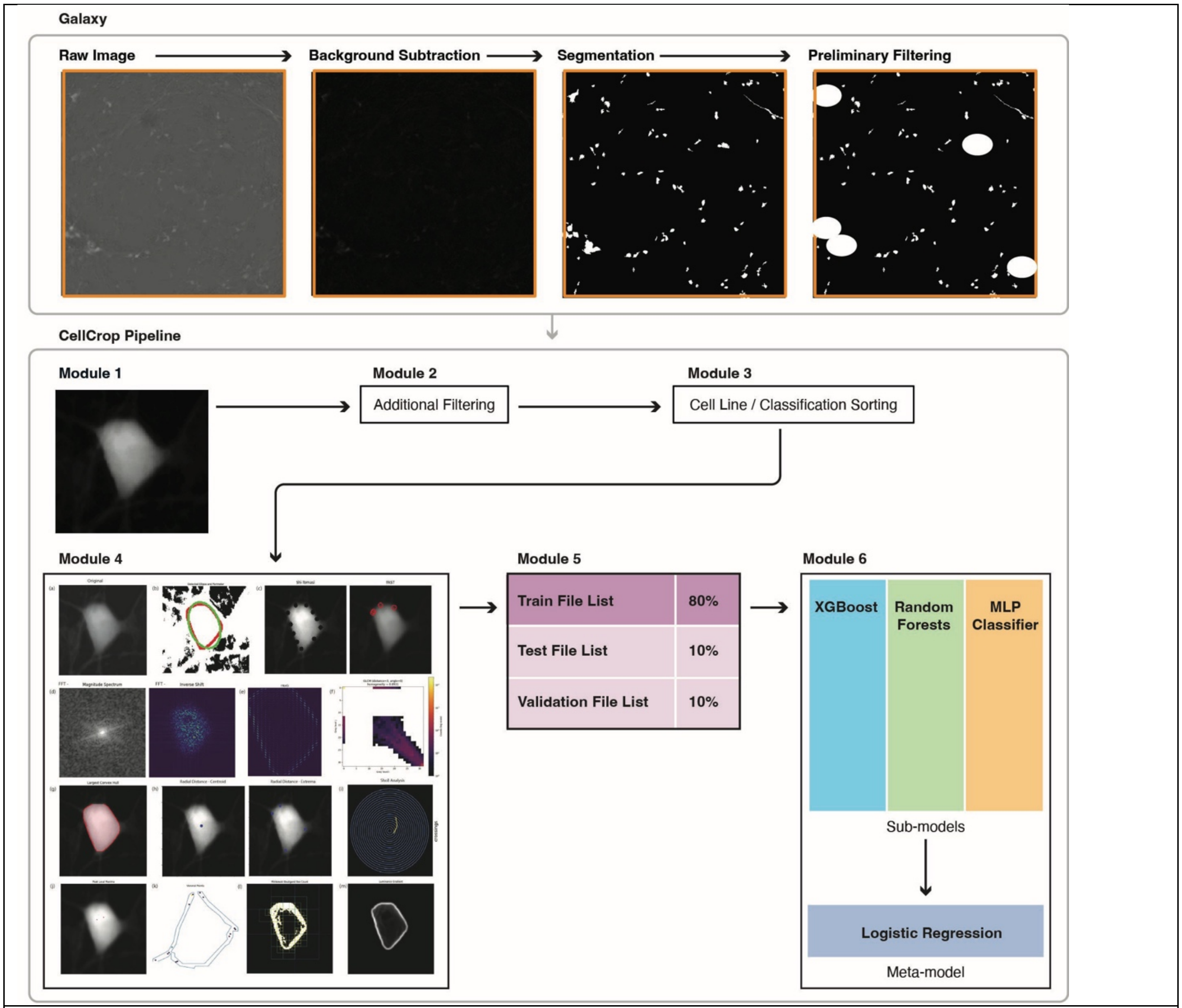


**Supplementary Figure 10. Galaxy and CellCropPipeline workflows for image processing and data preparation for ML.** The Galaxy[84, 87] workflow performs initial image processing steps, including background subtraction, segmentation to generate encoded cell masks, and filtering based on eccentricity and intensity thresholds. The processed montages and associated cell masks are then input into our custom- built CropCellPipeline for downstream analysis. In Module 1, 115 × 115-pixel crops are generated around detected cells. Module 2 applies additional filtering, removing crops in which cells touch image borders. Module 3 organizes cropped images into their respective cell lines and into control versus ALS classification categories. Module 4 extracts a set of engineered features from each crop and compiles the results into a CSV file. Module 5 balances the dataset by experiment, cell line, and classification group, then partitions it into training, validation, and test subsets using an 80%/10%/10% split. Finally, Module 6 uses these subsets to train a stacked classifier model composed of XGBoost Random Forest, and Multilayer Perceptron sub-models, with a Logistic Regression meta-model for final prediction.

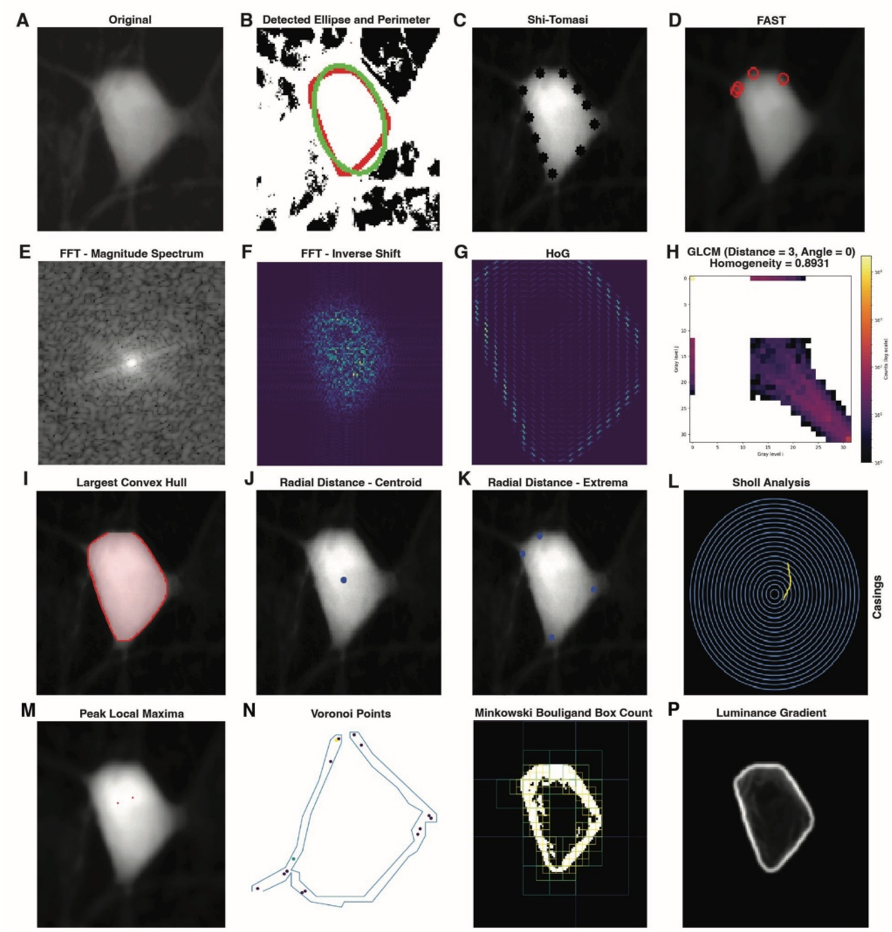


**Supplementary Figure 11. Feature extraction captures diverse morphological and textural features in cellular images. A)** Representative image crop of an iMN and the extracted features used in the SML aspect of this study that capture distinct aspects of cell morphology, texture, and spatial organization. Images on the top row show examples of features that capture shape and edge detection results, showing cellular boundaries and local feature points identified by ellipse fitting such as **B**) contour perimeter[120], contour area[120], and contour eccentricity as calculated with ellipse fitting[121]. **C**) Shi-Tomasi weighted corner detection[134] and FAST[135]. Features on the second row highlight texture and frequency-domain characteristics using **D**) Fast Fourier Transform Shift Magnitude Median[126] and Inverse FFT Shift Magnitude[126]. **E**) Histogram of gradients feature (HOG)[127]. **F**) Gray-Level Co-occurrence Matrix Homogeneity[136] which together reveal patterns of structural anisotropy and local texture contrast. The third row shows morphological descriptors, such as **G**) Solidity[122] as calculated from the convex hull subtracted from the contour area. **H**) Radial Distance as calculated from centroid and extrema of cell[123] which together reveal patterns of structural anisotropy and local texture contrast. **I**) Sholl analysis, quantifying cell shape complexity and branching organization[130] with skeletonization transformation[131] . The bottom row depicts higher-order structural and spatial measures, including **J**) Local Peak Maxima[124]. **K**) Voronoi Points[132] evaluated as Clusters with DBSCAN[133]. **L**) The fractal dimension Minkowski–Bouligand box counting[128]. **M**) Luminance gradients outlining boundary definition as Luminance as measured with Sobel Gradient (LG)[125]. Together, these analyses combine classical image processing and geometric feature extraction to generate a comprehensive quantitative representation of cell morphology and texture that uncover multiple geometric and textural dimensions of cellular architecture relevant for quantitative phenotypic characterization.

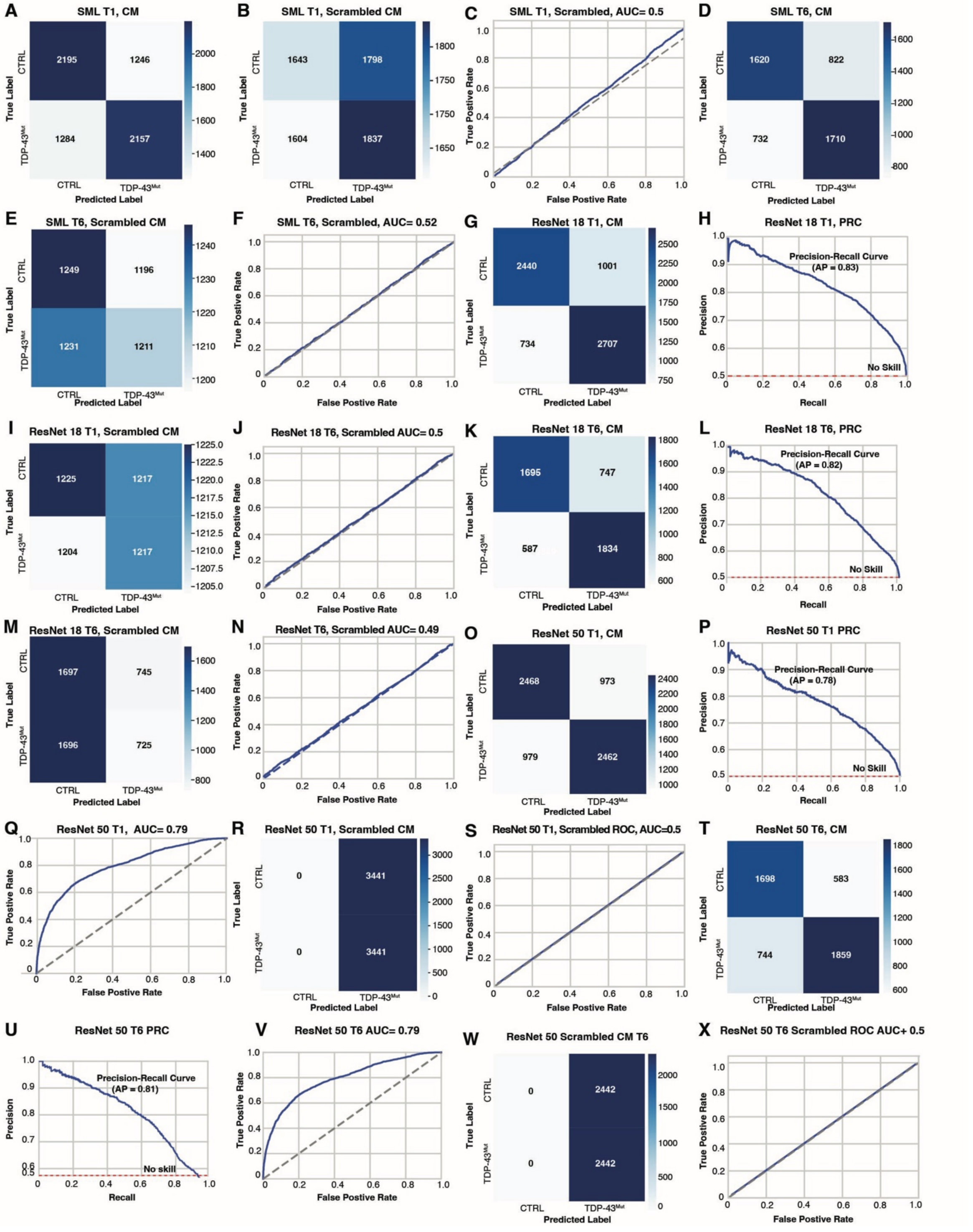


**Supplementary Figure 12. Performance of SML and DNN models for T1 and T6 classification**. **A**) Confusion matrix (CM) illustrating the performance of the SML Stacked Ensemble Classifier in distinguishing CTRL and TDP-43mut groups in T1. The CM displays the number of samples for each combination of true labels (rows) and predicted labels (columns). Correct classifications are shown along the diagonal, with 2,195 CTRL and 2,157 TDP-43$^{mut}$ samples correctly identified. Off-diagonal values represent misclassifications, where CTRL samples were predicted as TDP-43$^{mut}$ (1,246) and

TDP-43$^{mut}$ samples were predicted as CTRL (1,284). Color intensity reflects the number of samples in each cell. **B**) CM showing model predictions after random permutation of class labels in the SML T1 condition. Correct and incorrect classifications are distributed approximately evenly across classes, consistent with chance-level performance and indicating the absence of meaningful class-associated signal following label scrambling. **C**). ROC curve corresponding to the SML T1 model trained with randomized labels show an AUC of 0.5, indicating chance-level discrimination and confirming that model performance depends on true class labels rather than data artifacts or leakage. **D**) CM displaying the classification performance of the SML Stacked Ensemble Classifier in distinguishing CTRL and TDP-43$^{mut}$ groups in T6. This model performed slightly better than the T1 model. The model correctly classified 1620 CTRL samples and 1710 TDP-43$^{mut}$ samples. Misclassification was moderate with 732 CTRL samples predicted as TDP-43$^{mut}$ and 822 TDP-43mut samples predicted as CTRL. **E)** CM showing model predictions after random permutation of class labels in the SML T6 condition. Correct and incorrect classifications are distributed approximately evenly across classes, consistent with chance-level performance and indicating the absence of meaningful class-specific signal following label scrambling. F) ROC) curve for SML T6 classification with scrambled labels**. G)** CM for the T1 ResNet18 model classifying CTRL and TDP-43$^{mut}$ groups. The model correctly classified 2,440 CTRL and 2,707 TDP-43$^{mut}$ samples, with 1,001 CTRL predicted as TDP-43$^{mut}$ and 734 TDP-43$^{mut}$ predicted as CTRL. **H**) Precision Recall Curve (PRC) for the T1 ResNet18 classifier yields an average precision (AP) of 0.83, indicative of strong model performance. **I**) CM showing model predictions after random permutation of class labels of ResNet18 in T1. **J)** ROC curve for ResNet18 in T1 classification with scrambled labels. **K**) CM for the T6 ResNet18 classifier show similar precision as T1 shown in G. The model correctly identified 1,695 CTRL and 1,834 TDP-43$^{mut}$ iMNs with lower misclassification counts (747 CTRL predicted as TDP-43$^{mut}$ and 587 TDP-43$^{mut}$ predicted as CTRL). **L)** PRC for the T6 ResNet18 classifier showing and AP of 0.82. **M**) CM showing model predictions after random permutation of class labels of ResNet18 in T6. **N)** ROC curve for ResNet18 in T6 classification with scrambled labels. **O**) CM showing ResNet50 model performance at T1. The model correctly classified 2468 CTRL samples and 2462 TDP-43$^{mut}$ samples. Misclassification rates were balanced across groups showing no bias, with 973 CTRL samples predicted as TDP-43$^{mut}$ (false positives) and 979 TDP-43$^{mut}$ samples predicted as CTRL (false negatives). **P**) PRC at T1 for ResNet50 shows good discriminatory ability with an AP of 0.78. **R**) CM showing model predictions after random permutation of class labels of ResNet50 in T1. **S)** ROC curve for ResNet50 in T1 classification with scrambled labels. **T)** CM for ResNet50 at T6. The model correctly classified 1698 CTRL samples and 1859 TDP-43$^{mut}$ samples. Misclassification was slightly higher with TDP-43$^{mut}$ being classified as controls, with 583 CTRL samples predicted as TDP-43$^{mut}$ and 744 CTRL classified as TDP-43$^{mut}$ samples predicted as CTRL. **U**) PRC illustrating the performance of the ResNet50 DNN at T6 showing an AP of 0.81, indicating a stronger discriminatory ability at the later timepoint. **V**) ROC curve for ResNet50 at T1 AUC= 0.79. **L**) ROC curve for ResNet50 at T6, AUC= 0.79. For all PRCs, the red dashed line represents the expected performance of a classifier making random predictions. **W)** CM showing model discriminative ability after scrambling the class labels. **X)** ROC showing model predictions after random permutation of class labels of TDP-43$^{mut}$ and 744 CTRL iMNs.

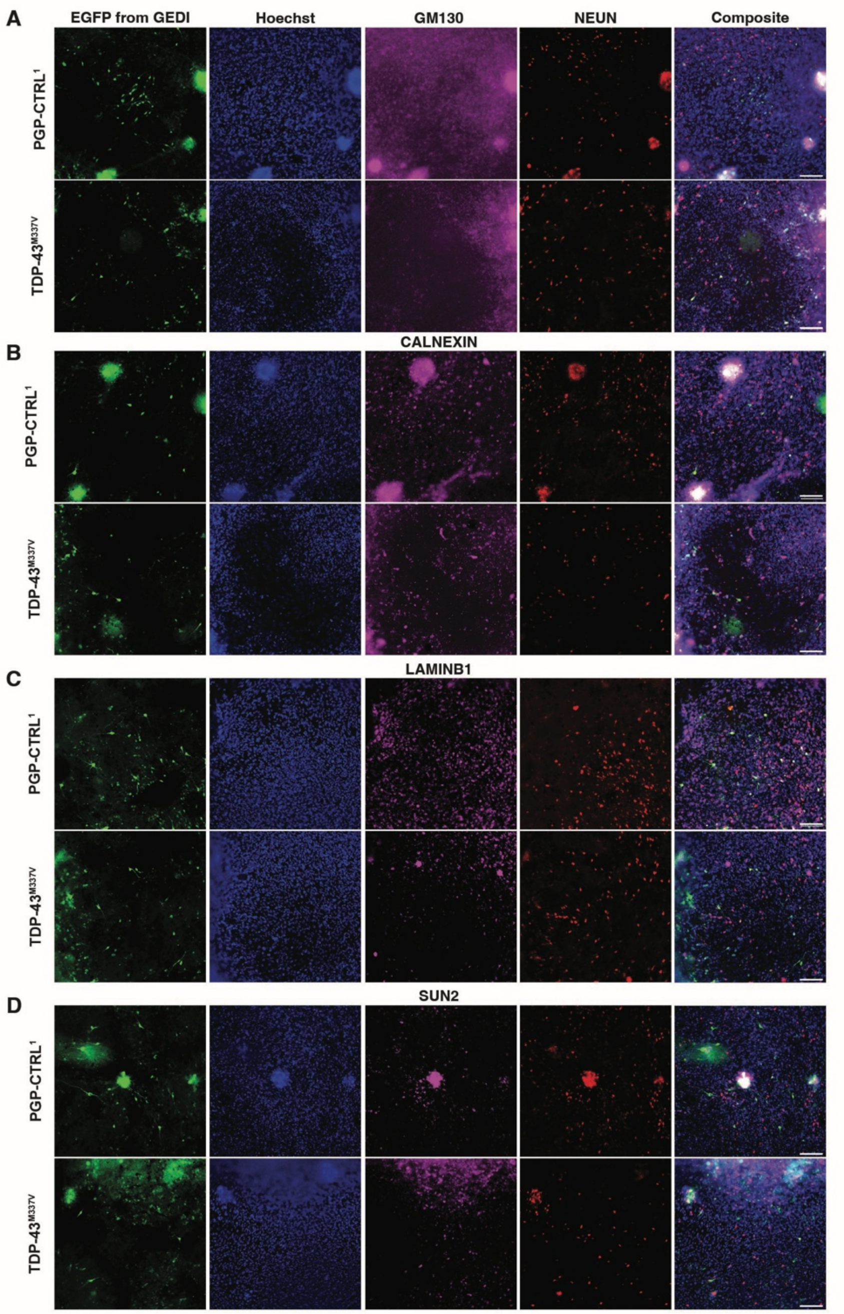


**Supplementary Figure 13. Example images of immunofluorescence staining using antibodies that label different cell compartments in iMNs. A-D**) Representative immunofluorescence images showing expression of EGFP (green) of GEDI, nuclear stain Hoechst (blue), organelle-specific markers (magenta), and neuronal marker NEUN (red) in control and TDP-43$^{M337V}$ iMNs. Organelle-specific markers are GM130 (Golgi), CALNEXIN (endoplasmic reticulum), and LAMINB1 and SUN2 (nuclear envelope). Far-right images are merges. Scale bars: ~90 µm.

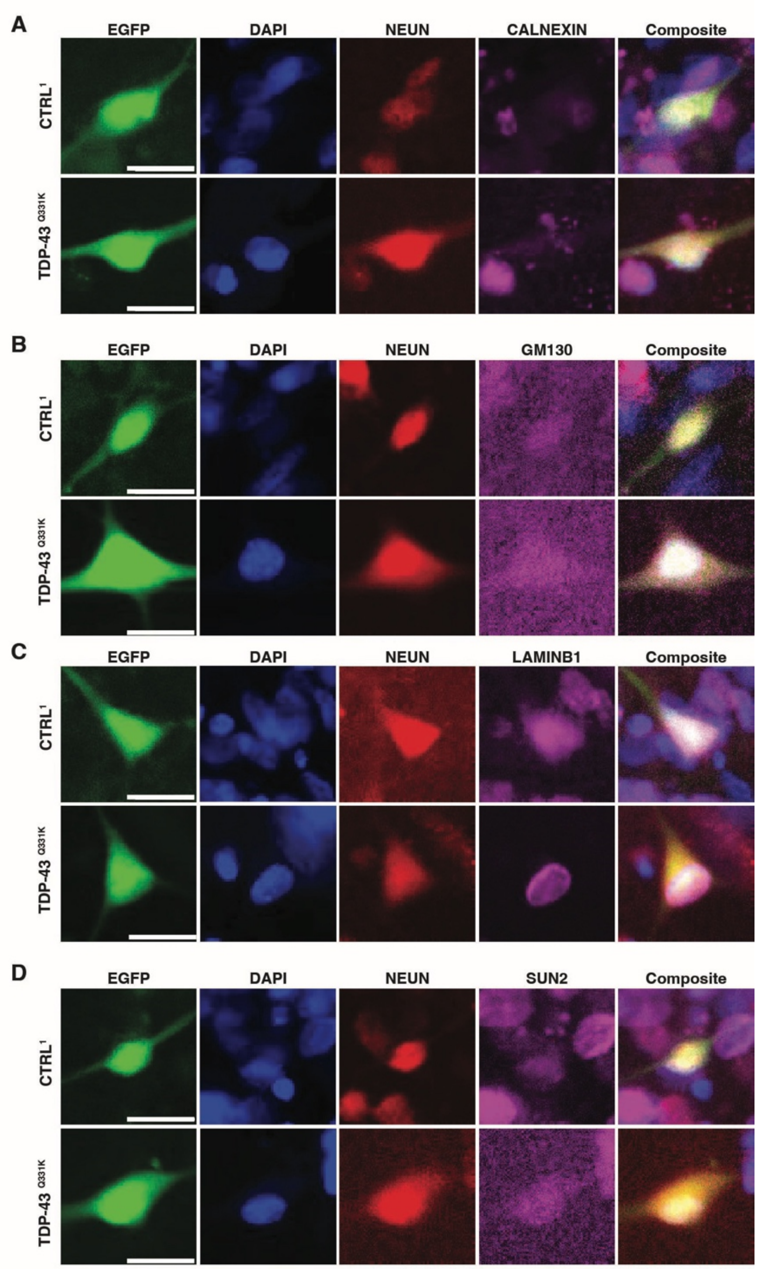


**Supplementary Figure 14. Example image crops of immunofluorescence-labelled CTRL and TDP-43$^{mut}$ iMNs used for saliency mapping. A-D)** Representative immunofluorescence images of 4 TDP-43$^{Q331K}$ iMNs and 4 control IMNs expressing the GEDI EGFP (green) and stained with DAPI (nuclei, blue), NEUN (neuronal marker, red), and **A**) CALNEXIN (endoplasmic reticulum), **B**) GM130 (Golgi), **C**) LAMIN B1 (nuclear envelope) and **D**) SUN2 (inner nuclear membrane). In all panels, scale bar = ~20 μm.

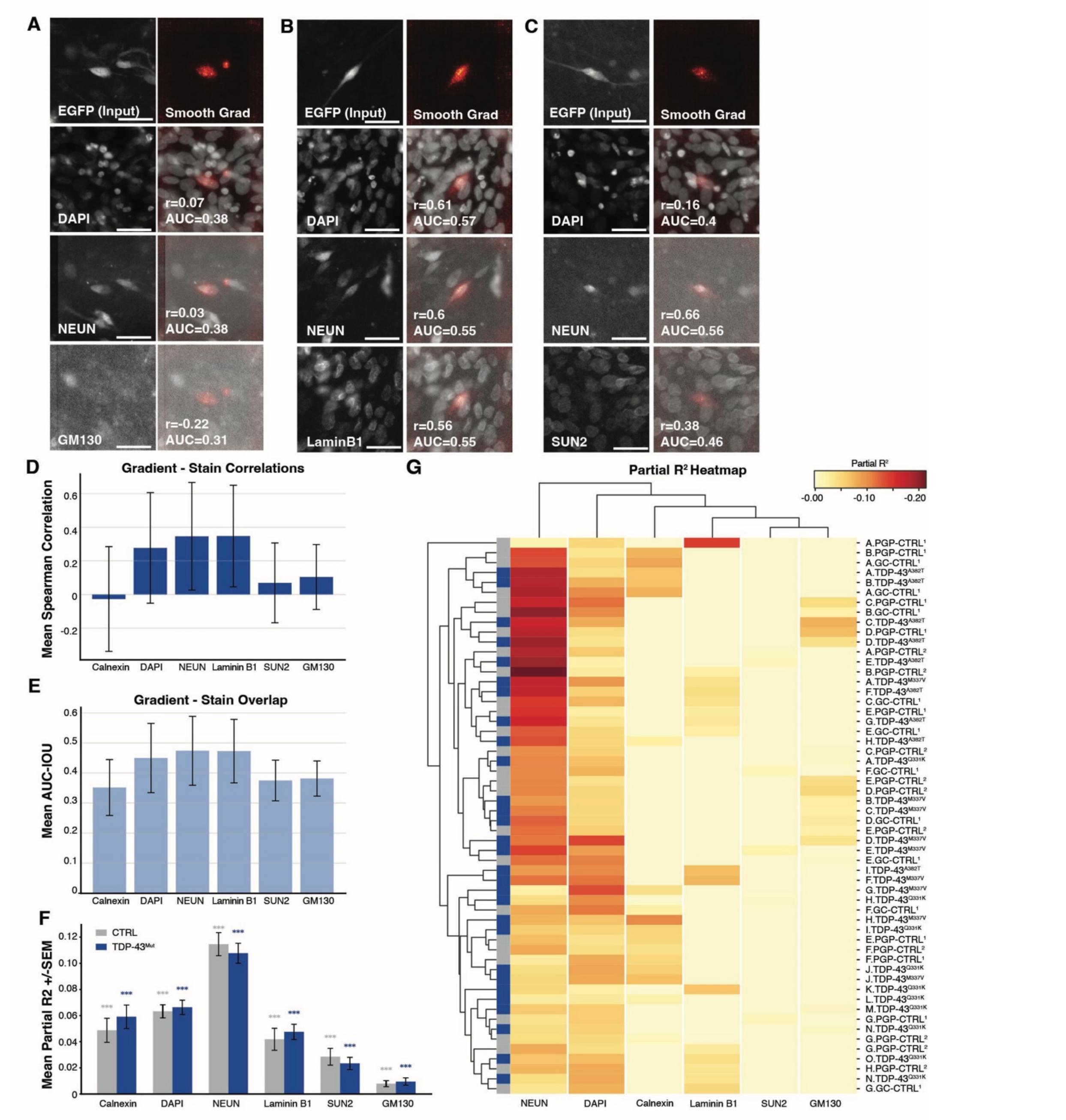


**Supplementary Figure 15. Gradient–stain correspondence and feature contributions across markers identify subcellular location of the signal that distinguishes TDP-43$^{mut}$ from controls. A–C**) Representative examples of EGFP from GEDI in the FITC channel input images (left) and corresponding SmoothGrad saliency maps (right) for control samples, shown alongside individual immunofluorescence channels. Panels include DAPI, NEUN, LaminB1 or SUN2 and GM130. Scale bar= ~10μM. For each stain, Spearman correlation (ρ) between stain intensity and gradient magnitude, as well as AUC-overlap (AUC-o), are reported. **D**) Boxplots showing mean Spearman correlations between SmoothGrad gradients and each stain across all samples (n=463 cells; Control n=218, TDP-43 n=245; sample sizes per stain: DAPI/NeuN n=462, SUN2 n=141, LaminB1 n=137, Calnexin n=129, GM130 n=54). LaminB1, NeuN, and DAPI exhibited the highest correlations. **E**) Mean AUC-overlap (AUC-o) values quantifying spatial correspondence between gradients and stain intensities. Error bars represent SEM.

NeuN, LaminB1, and DAPI show the highest overlap. **F**) Bar graph of mean partial R² values (±SEM) for each stain, comparing control (blue) and TDP-43 mutant (red) groups. NeuN showed the highest partial R² in both groups. No between-group differences in partial R² reached statistical significance. **G**) Hierarchically clustered heatmap of partial R² values for all samples, grouped by condition. Warmer colors indicate greater unique variance explained by each stain. Clustering highlights condition-specific gradient–stain patterns, with nuclear and neuronal markers contributing most strongly to classifier explainability.

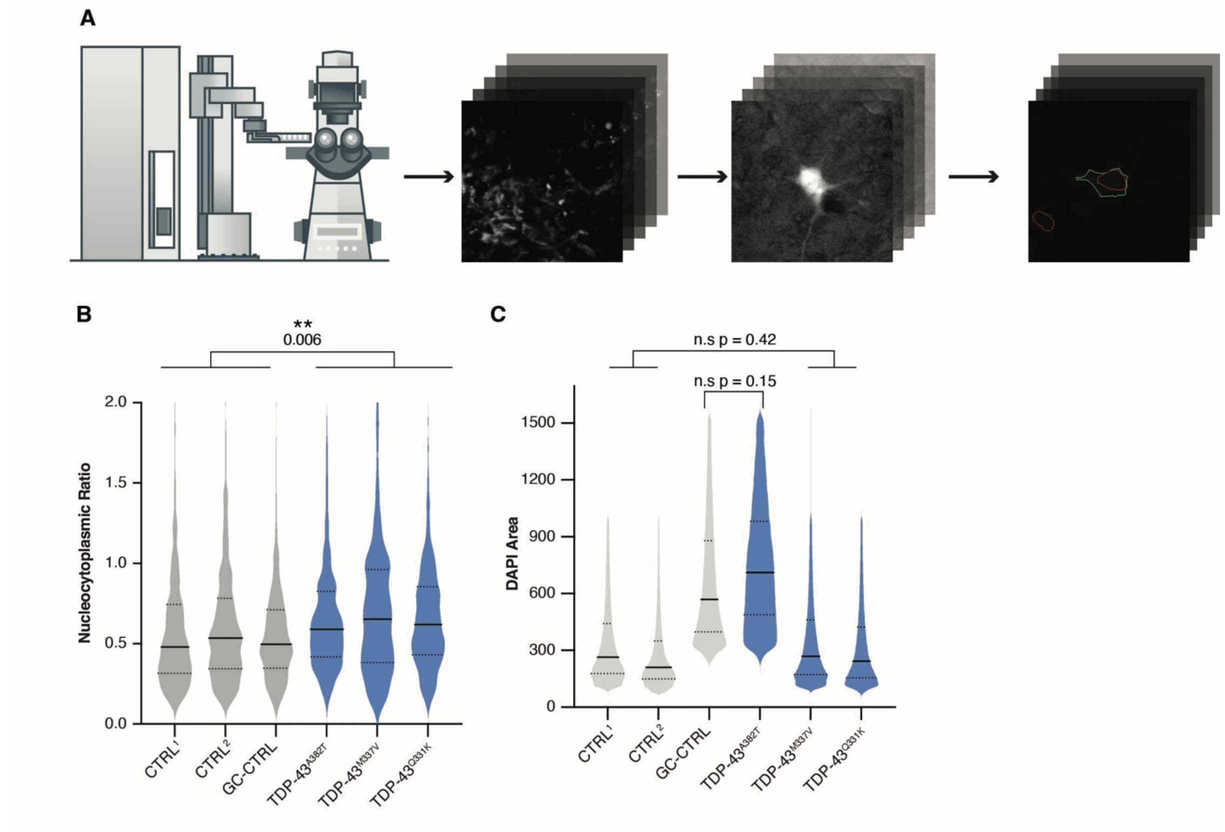


**Supplementary Figure 16. TDP-43$^{muts}$ display evidence of altered nucleocytoplasmic shuttling compared to gene- edited controls**.

**A**) Workflow for image acquisition of the nucleocytoplasmic shuttling biosensor to perform quantitative analysis of nuclear/cytoplasmic ratios. Images of iMNs containing the 2GiR construct 2Gi2R[96] first acquired using an automated microscope, followed by preprocessing to enhance signal and contrast. Individual cells are segmented to distinguish nuclear and cytoplasmic regions, the area occupied by each fluorescent biosensor (NES-GFP vs NLS-RFP) is captured and the nuclear-to-cytoplasmic ratio (RFP/GFP area ratio) is quantified on a per cell basis. **B**) Violin plots showing the distribution of nuclear-to-cytoplasmic ratios of the fluorescence of the biosensor across all lines. Data represent individual cell measurements; black lines indicate median values. LMM, 0.16, p = 0.006**. TDP-43$^{M337V}$ n = 668 cells, TDP-43$^{Q331K}$ n = 651 cells, TDP-43$^{A382T}$ n = 699 cells, PGP-CTRL$^{1}$ n = 1,064 cells, PGP-CTRL$^{2}$ n = 727 cells, GC-CTRL$^{1}$ n =676 cells. **C**) There is no significant difference in nuclear size between TDP-43$^{mut}$ iMNs and their gene- edited or corrected control (see also **Figure 5**). iMNs were fixed and stained with DAPI and imaged. The size of the nuclei was measured using a custom Galaxy pipeline described above. Data represent individual cell measurements; black lines indicate median values. Nuclei of the TDP-43$^{A382T}$ and its gene corrected control (GC-CTRL) display overall larger size compared to the PGP-CTRL$^{1}$, PGP-CTRL$^{2}$, TDP-43$^{M337V}$ and TDP-43$^{Q331K}$. Therefore, to determine if the TDP-43 mutations alter nucleus size, we analyzed the two groups separately. TDP-43$^{A382T}$ vs. GC-CTRL is n.s., p = 0.15, estimate = 0.12. PGP-CTRL$^{1}$, PGP-CTRL$^{2}$ vs. TDP-43$^{M337V}$ and TDP-43$^{Q331K}$, n.s., p = 0.42, estimate = 0.09. TDP-43$^{M337V}$ n = 26,071 cells, TDP-43$^{Q331K}$ n =125,816 cells, TDP-43$^{A382T}$ n = 41,628 cells, PGP-CTRL$^{1}$ n = 112,880 cells, PGP-CTRL$^{2}$ n = 261, 076 cells, GC-CTRL$^{1}$ n = 71,033 cells. All statistical analysis was performed using a GLMM[110].

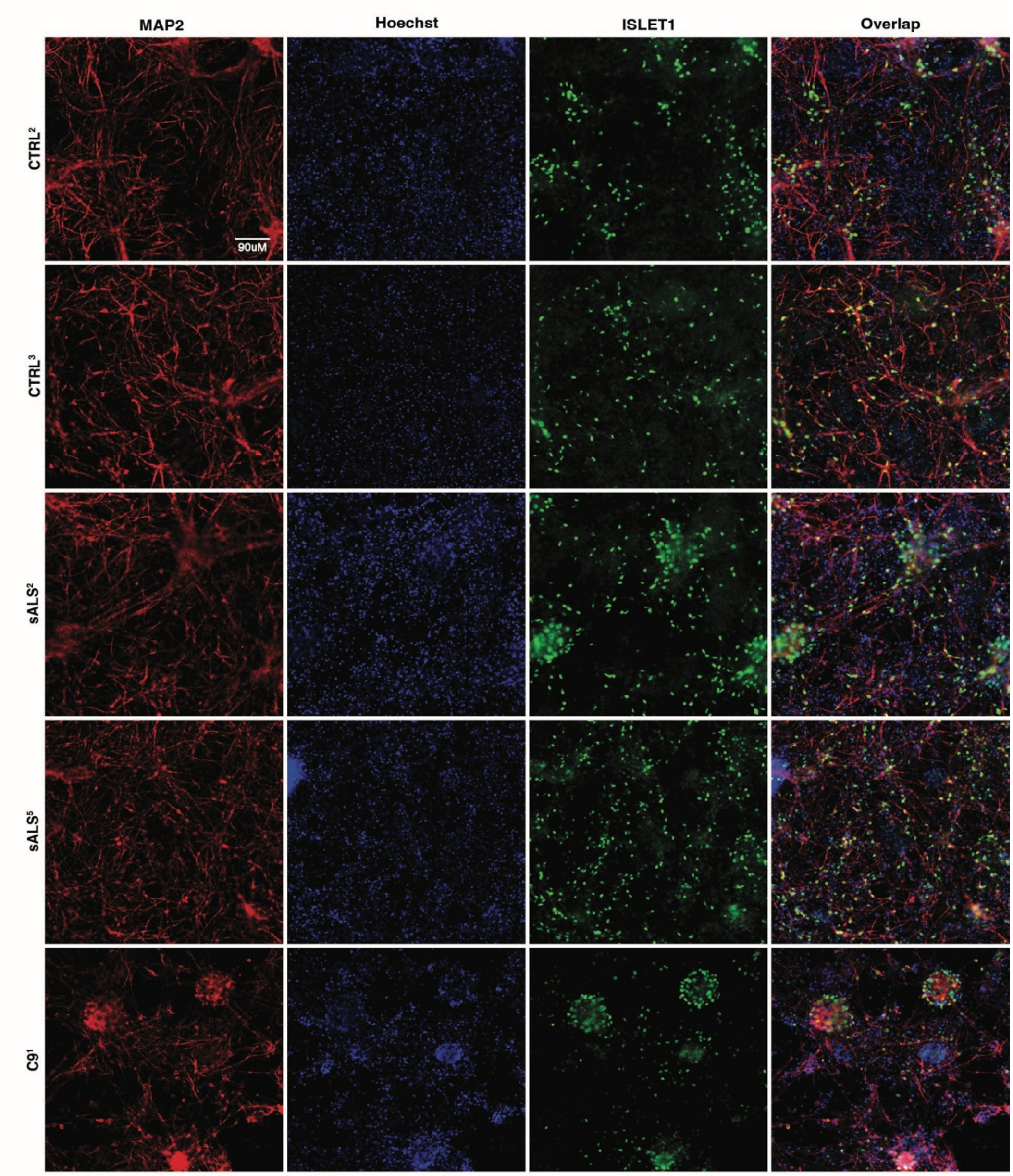


**Supplementary Figure 17. Expression of the motor neuron marker ISLET1 in C9orf72$^{mut}$, sALS and control iMNs.** Representative images of control, sALS and C9orf72$^{mut}$ iMNs stained for the motor neuron marker ISLET1. At differentiation day ~ DIV 35, iMNs were fixed and stained using antibodies against MAP-2 (red), ISLET1 (green) and DAPI (blue). Merged image is on the far right, scale bar = ~90 uM.

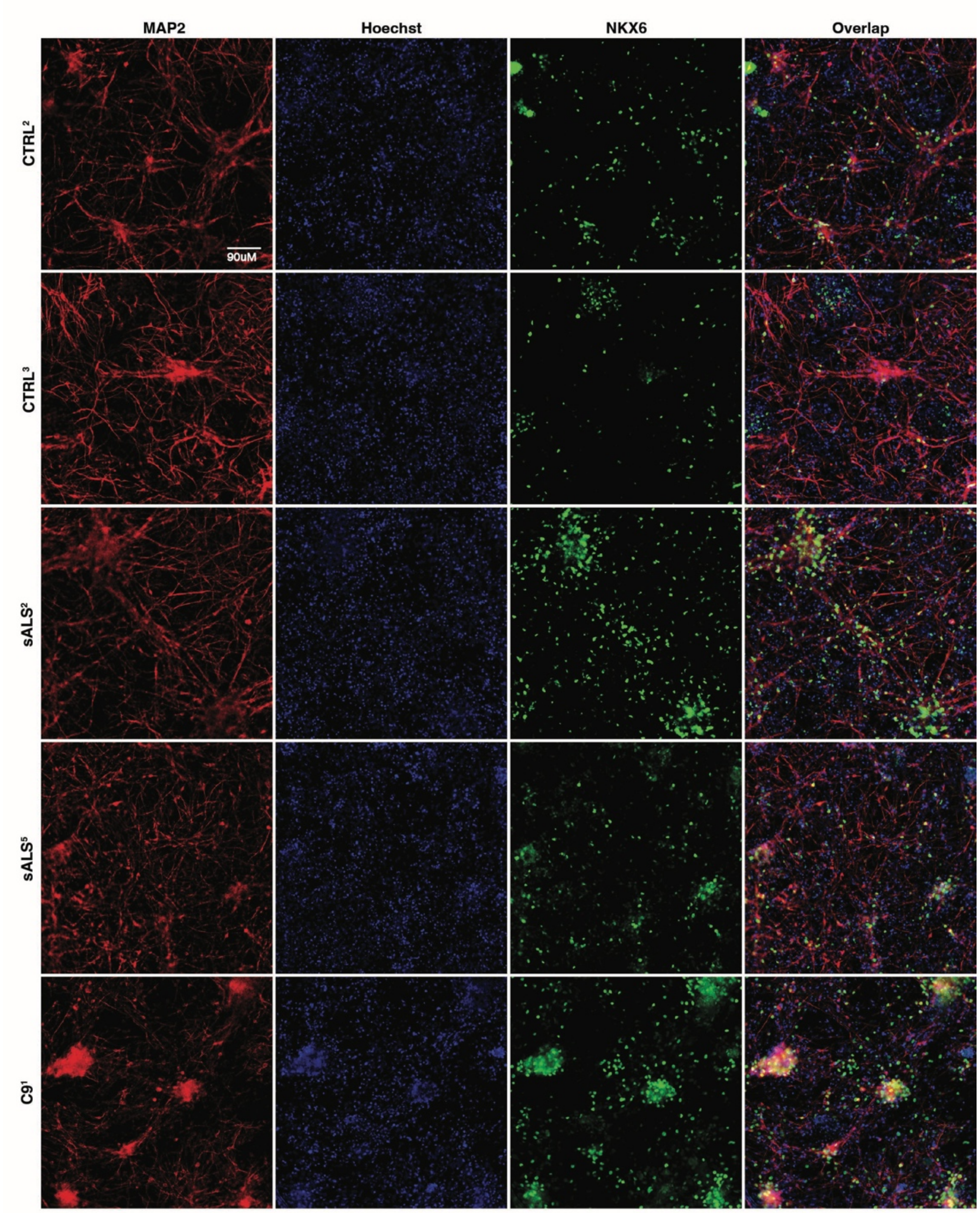


**Supplementary Figure 18. Expression of the motor neuron marker NKX6.2 in C9orf72$^{mut}$, sALS and control iMNs.** Representative images of sALS and C9orf72$^{mut}$ iMNs stained for the motor neuron marker NKX6.2. At differentiation day ~ DIV 35, iMNs were fixed and stained using antibodies against MAP-2 (red), NKX6.2 (green), and DAPI (blue). Merged image is on the far right, scale bar = ~90 uM.

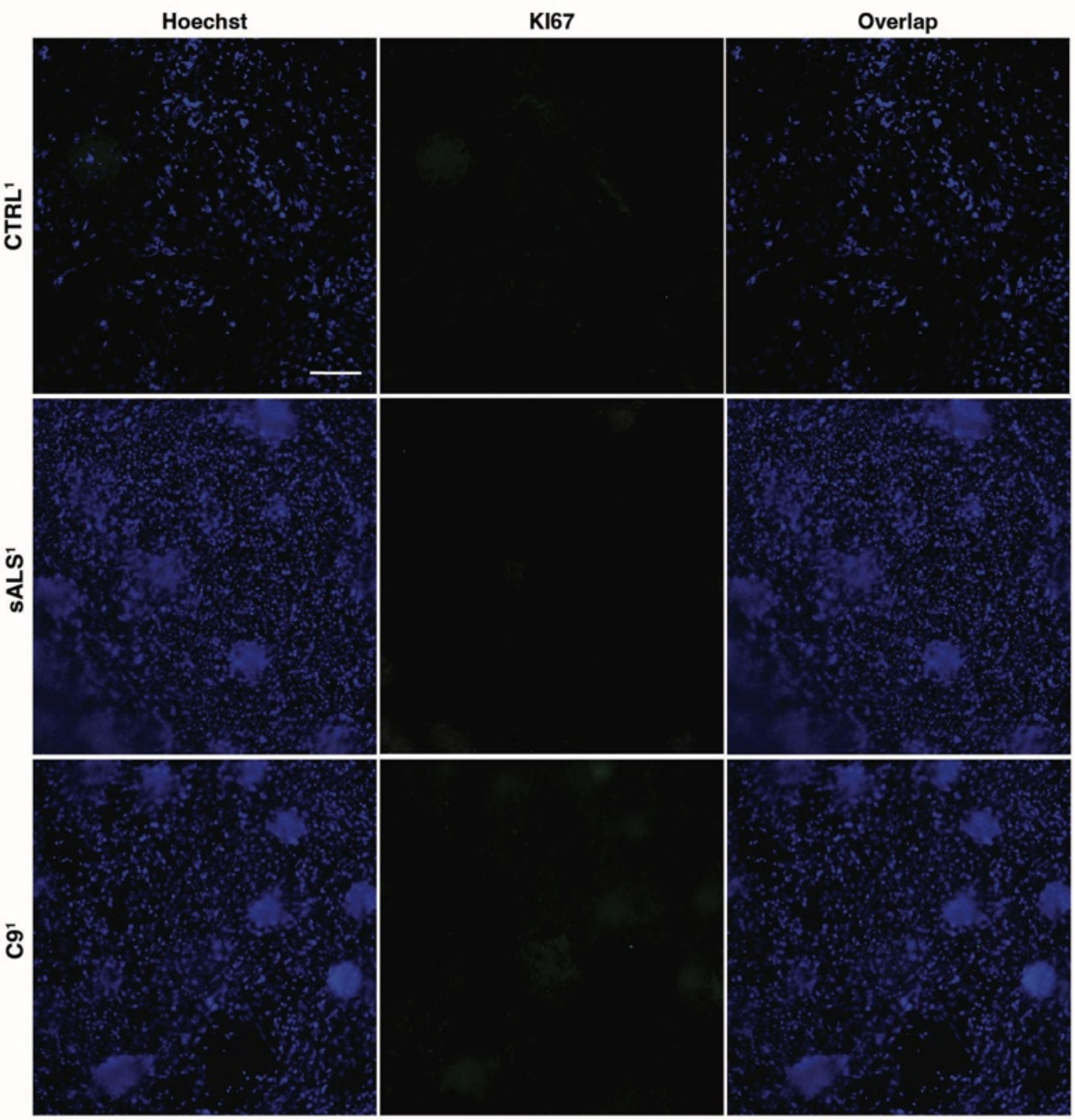


**Supplementary Figure 19. KI67 expression is very low in sALS, C9orf72$^{mut}$ and control iMNs.** Representative images of differentiation day ~DIV 35 sALS and C9orf72$^{mut}$ iMNs stained for the proliferation marker KI67 (red) and with DAPI (blue). Merged image is on the far right, scale bar = ~90 uM.

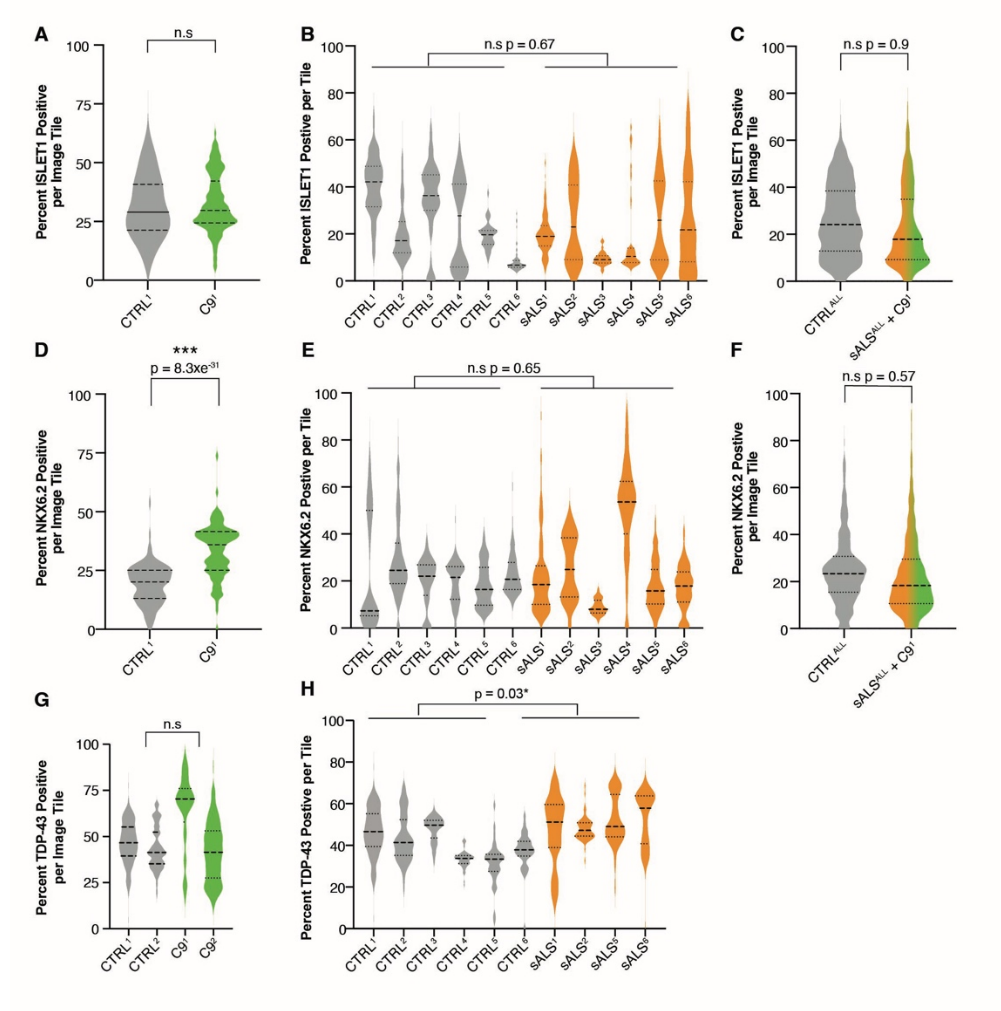


**Supplementary Figure 20. Quantification of motor neuron markers across sALS, C9orf72$^{mut}$ and control iMNs. A**) Violin plot showing the percentage of ISLET1-positive motor neurons per image tile in CTRL versus C9orf72$^{mut}$ iMNs. No significant difference was observed between groups (n.s.). A binomial GLMM confirmed the absence of group-level effects (estimate = 0.04, p = 0.49). CTRL$^{1}$ n= 93,570 cells, C9$^{1}$ n= 89,430 cells. **B**) Distribution of ISLET2-positive cells across six CTRL lines and six sALS lines. Inter-line variability is evident within both groups, but overall comparison shows no significant difference using a Poisson GLMM (n.s., estimate = 0.066, p = 0.67), suggesting that sALS and CTRL cultures contain similar proportions of ISLET2-expressing neurons. CTRL$^{1}$ n = 75,359 cells, CTRL$^{2}$ n = 467,354 cells, CTRL$^{3}$ n = 107,094 cells, CTRL$^{4}$ n = 231,133 cells, CTRL$^{5}$ n = 74,304 cells, CTRL$^{6}$ n = 127,996 cells, sALS$^{1}$ n = 308, 452 cells, sALS$^{2}$ n = 183,752 cells, sALS$^{3}$ n = 73,090 cells, sALS$^{4}$ n = 121,063 cells, sALS$^{5}$ n = 283,543 cells, sALS$^{6}$ n = 174,223 cells. **C**) Comparison of ISLET1-positive cell percentages between pooled CTRL, pooled sALS, and pooled sALS + C9 groups from **A** and **B**. No significant difference was detected using a Poisson GLMM (n.s., estimate = -0.01, p = 0.94), further supporting that ISLET1 expression is not altered across ALS subtypes. **D**) Violin plot showing the percentage of NKX6.2-positive cells per image tile for CTRL and C9orf72$^{mut}$ iMNs. A binomial GLMM shows significant increase in NKX6.2-positive cells in the C9orf72$^{mut}$ compared to controls (estimate = 0.8, p = 8.3 × $10^{-31}$*). CTRL$^{1}$ n = 33,479 cells, C9$^{1}$ n = 53,417 cells. **E**) Quantification of NKX6.2 positivity across individual CTRL and sALS lines. Despite variability among cell lines, no significant difference was detected between the CTRL and sALS groups using a Poisson GLMM (n.s., estimate = 0.14, p = 0.65), suggesting preserved NKX6.2 expression in sporadic ALS cultures. CTRL$^{1}$ n = 21,439 cells, CTRL$^{2}$ n = 243,581 cells, CTRL$^{3}$ n = 76,770 cells, CTRL$^{4}$ n = 148,460 cells, CTRL$^{5}$ n = 43,252 cells, CTRL$^{6}$ n = 114,291 cells, sALS$^{1}$ n = 178,896 cells, sALS$^{2}$ n = 103,469 cells, sALS$^{3}$ n = 49,446 cells, sALS$^{4}$ n = 7,378 cells, sALS$^{5}$ n = 230,081 cells, sALS$^{6}$ n = 140,542 cells. N = 7 experiments, 2-3 experiments per cell line. **F**) NKX6.2 quantification for pooled CTRL, sALS, + C9orf72$^{mut}$ groups. No significant group-level differences were observed (n.s., estimate = 0.14, p = 0.57), further indicating that NKX6.2 expression changes are specific to C9orf72 mutation rather than general ALS pathophysiology. **G**) Violin plot showing that there is no difference in the percentage of TDP-43-positive cells per image tile for CTRL and C9orf72$^{mut}$ groups using a negative binomial GLMM (n.s., estimate = 0.009, p = 0.93). CTRL$^{1}$ n = 346,894 cells, CTRL$^{2}$ n = 302,379 cells C9$^{1}$ n = 23, 734 cells, C$^{2}$ n = 300,151 cells, 2-4 experiments per line. **H**) Violin plot showing the percentage of TDP-43-positive cells per image tile sALS is higher than control using a negative binomial GLMM (estimate = 0.15, p = 0.03). CTRL$^{1}$ n = 346,894 cells, CTRL$^{2}$ n = 302,373 cells, CTRL$^{3}$ n = 140,148 cells, CTRL$^{4}$ n = 148,682 cells, CTRL$^{5}$ n = 62,203 cells, CTRL$^{6}$ n = 52,120 cells, sALS$^{1}$ n = 309,510 cells, sALS$^{2}$ n = 120,852 cells, sALS$^{5}$ n = 287,093 cells, sALS$^{6}$ n = 50,423 cells. All statistical analyses were performed using a GLMM[110]**.**

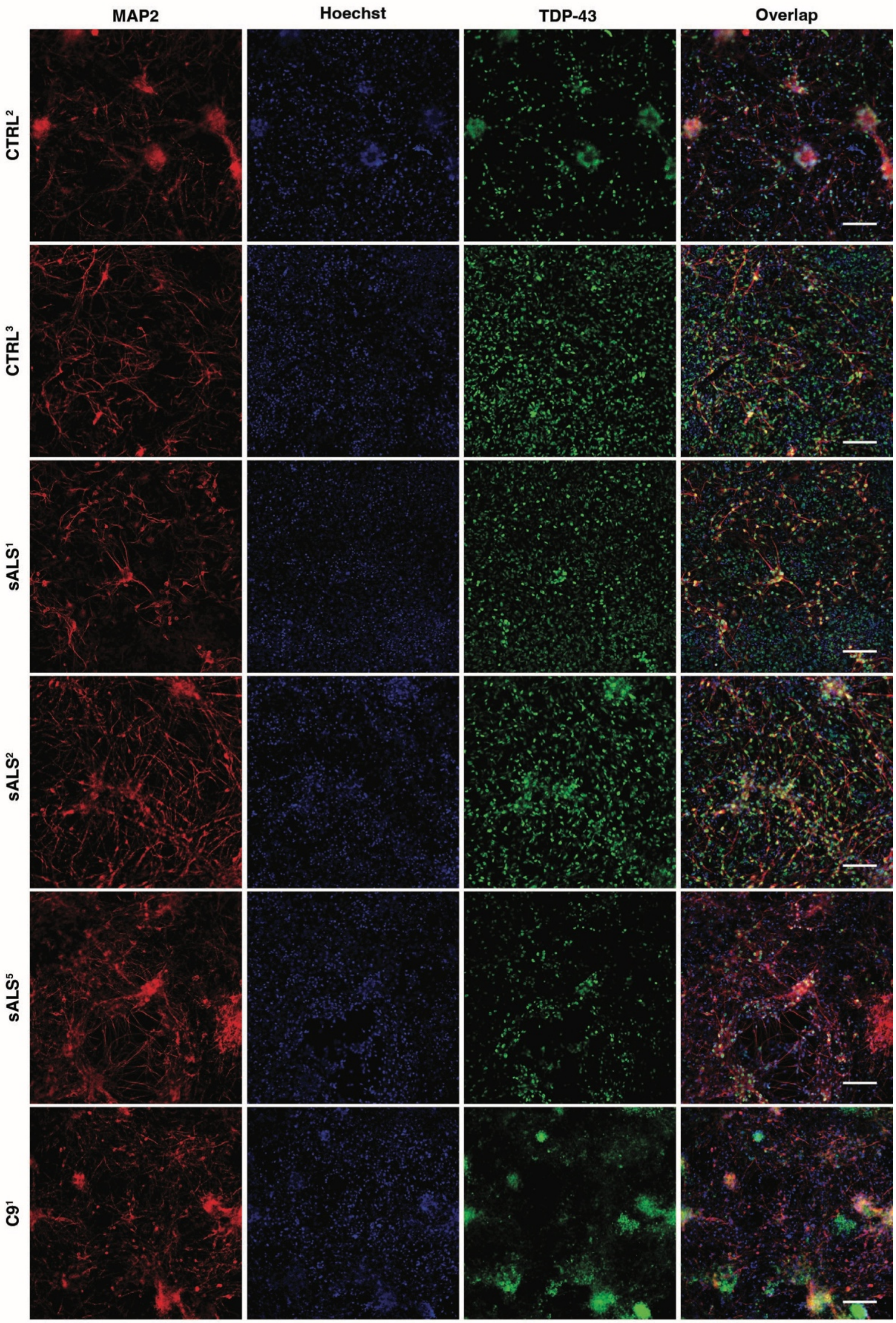


**Supplementary Figure 21. TDP-43 expression in sALS, C9orf72$^{mut}$ and control iMNs.** Representative images of differentiation day ~DIV 35 sALS and C9orf72$^{mut}$ iMNs stained for TDP-43 (red) and with DAPI (blue). Merged image is on the far right, scale bar = ~90 uM.

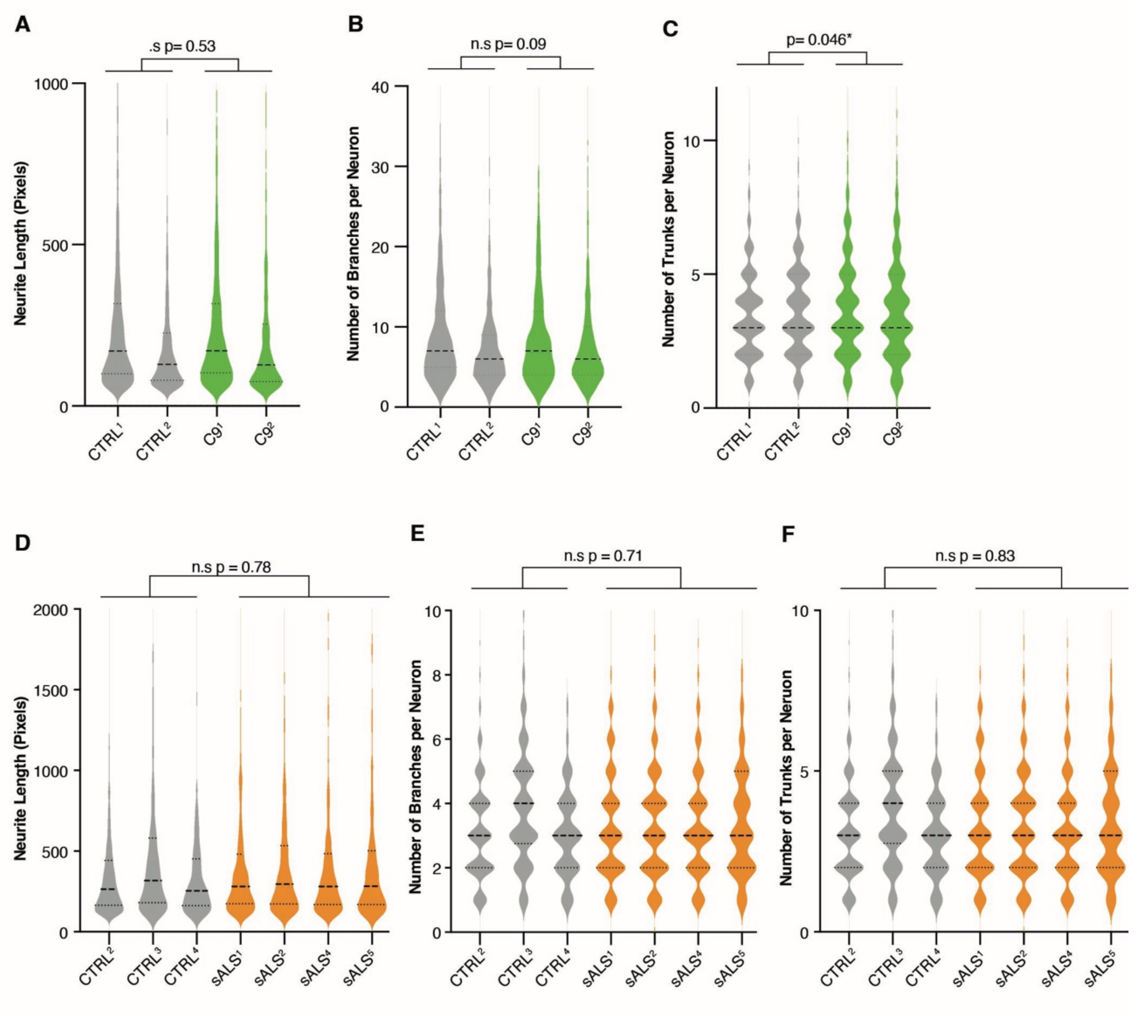


**Supplementary Figure 22. Neurite length and branching is comparable in C9orf72$^{mut}$ and sALS iMNs relative to controls. A–F**) Violin plots showing quantitative measurements of neurite morphology in iMNs derived from healthy controls (CTRL$^{1-4}$) and C9orf72$^{mut}$ patient iPSC lines (C9$^1$, C9$^2$) and sALS (sALS$^{1,2,4,5}$). **A**) Neurite length (pixels) is similar between control and C9orf72$^{mut}$ iMNs with no significant difference (estimate = -0.03, n.s., p = 0.53). B) Number of branches per neuron also shows no significant change between groups (estimate = -0.06, n.s., p = 0.09). **C**) Number of trunks per neuron is modestly, but significantly increased in C9 lines compared with controls (estimate = 0.05, p = 0.046). CTRL$^1$ n = 1,590 cells, CTRL$^2$ n = 706 cells, C9$^1$ n = 954 cells, C9$^2$ n = 685 cells. N = 6 experiments, 3 per line. **D**) Neurite length (pixels) per neuron is not significantly different between CTRL and sALS groups (estimate = 0.04, n.s., p = 0.78). **E**) The number of branches per neuron is not significantly different between CTRL and sALS neurons (estimate= -0.06, n.s., p = 0.71). **F**) Number of trunks per neuron likewise does not differ significantly between groups (estimate = -0.01, n.s., p = 0.83). CTRL[2] n = 1,107 cells, CTRL[3] n = 558 cells, CTRL[4] n = 200 cells, sALS[1] n = 675 cells, sALS[2] n = 710 cells, sALS[4] n = 646 cells, sALS[5] n = 318 cells, n = 4 experiments. Overall, these data show that baseline neuronal morphology is not significantly altered in C9 or sALS iMNs compared to controls. All statistical analyses were performed using a GLMM[110].

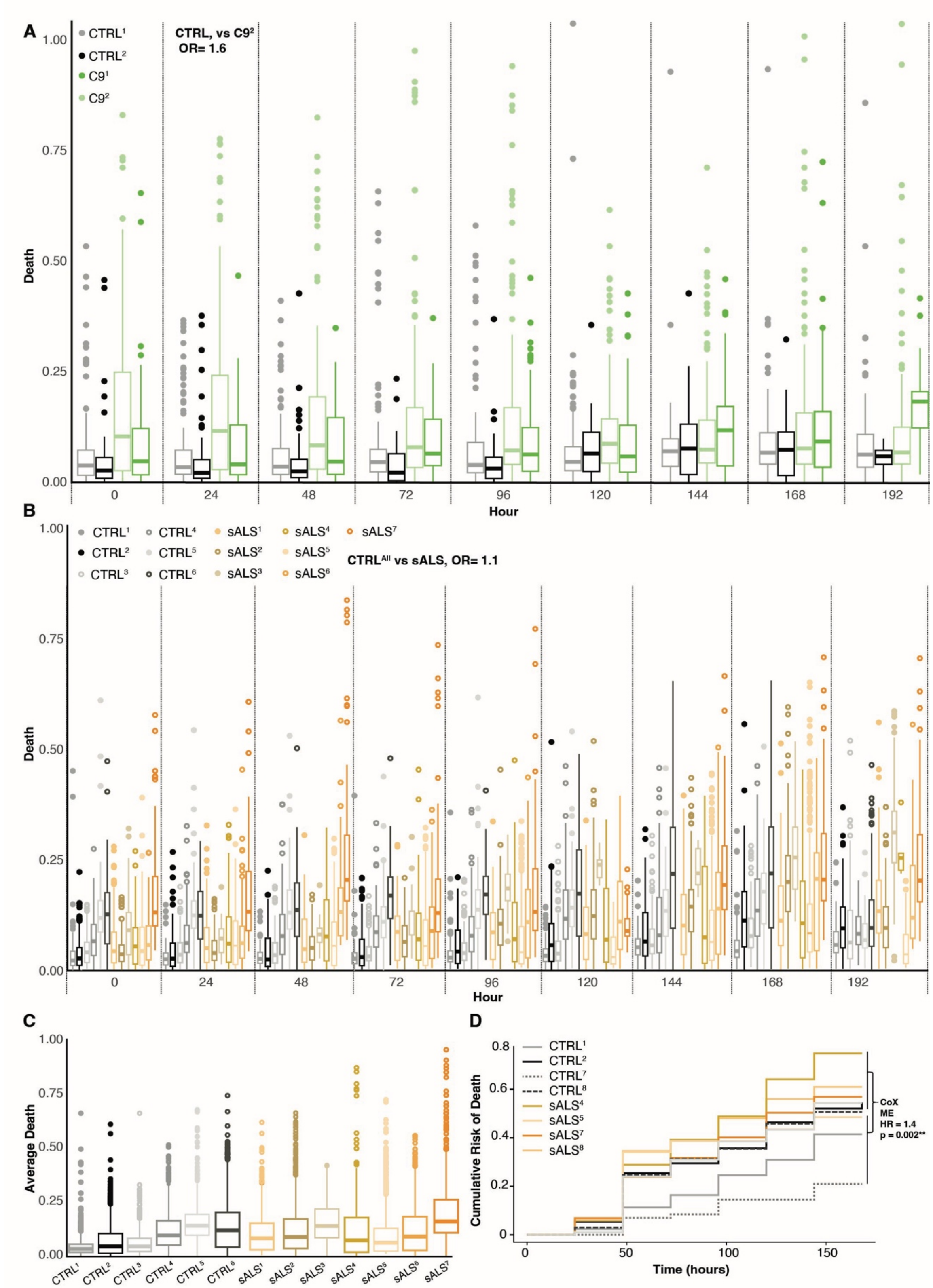


**Supplementary Figure 23. C9orf72$^{mut}$ and sALS iMNs display higher death rates than in controls. A**) Death fraction evaluated by GEDI reveals that C9$^1$ and C9$^2$ have elevated death rates compared to controls. CTRL$^1$ n = 217,107 cells, CTRL$^2$ n = 55,910 cells, C9$^1$ = 66,805 cells, C9$^2$ = 133,061 cells. From **Figure 7**, OR-

CD = 1.6, p = 6.7xe$^{-72.}$ **B**) There is high variability in the death fraction of control and sALS iMNs per line and over time. Nevertheless, the overall OR-CD is higher in sALS than control iMNs. From **Figure 7**, OR = 1.1, p = 1.0xe-29***. CTRL$^{1}$ n = 140,195 cells, CTRL$^{2}$ n = 390,275 cells, CTRL$^{3}$ n = 131,475 cells, CTRL$^{4}$ n = 63,278 cells, CTRL$^{5}$ n = 104,713 cells, CTRL$^{6}$ n = 115,246 cells, sALS$^{1}$ n = 281,844 cells, sALS$^{2}$ n = 177,053 cells, sALS$^{3}$ n = 22,705 cells, sALS$^{4}$ n = 78,182 cells, sALS$^{5}$ n = 251,155 cells, sALS$^{6}$ n = 175,111 cells, sALS$^{7}$ n =39,112 cells. **C**) The average death fraction across all time points from B. **D**) A subset of iMNs transduced with Synpasin:GFP were tracked and cell death was recorded using a custom Matlab script previously described[68, 74, 84, 87, 100, 103], which curates the last timepoint a human curator finds each neuron alive. The cumulative survival was determined by evaluating the instantaneous risk of death for individual neurons as previously described[68, 74, 84, 87, 100, 103] and modeled using a Cox mixed effects model to determine the rate of death for control and sALS iMNs. The HR = 1.4, p = 0.002. CTRL$^{1}$ n = 292 cells, CTRL$^{2}$ n = 293 cells, CTRL$^{3}$ n = 97 cells, CTRL$^{7}$ n = 73 cells, sALS$^{4}$ = 190 cells, sALS$^{5}$ = 299 cells, sALS$^{7}$ = 190 cells, sALS$^{8}$ = 266 cells, n = 4 experiments.

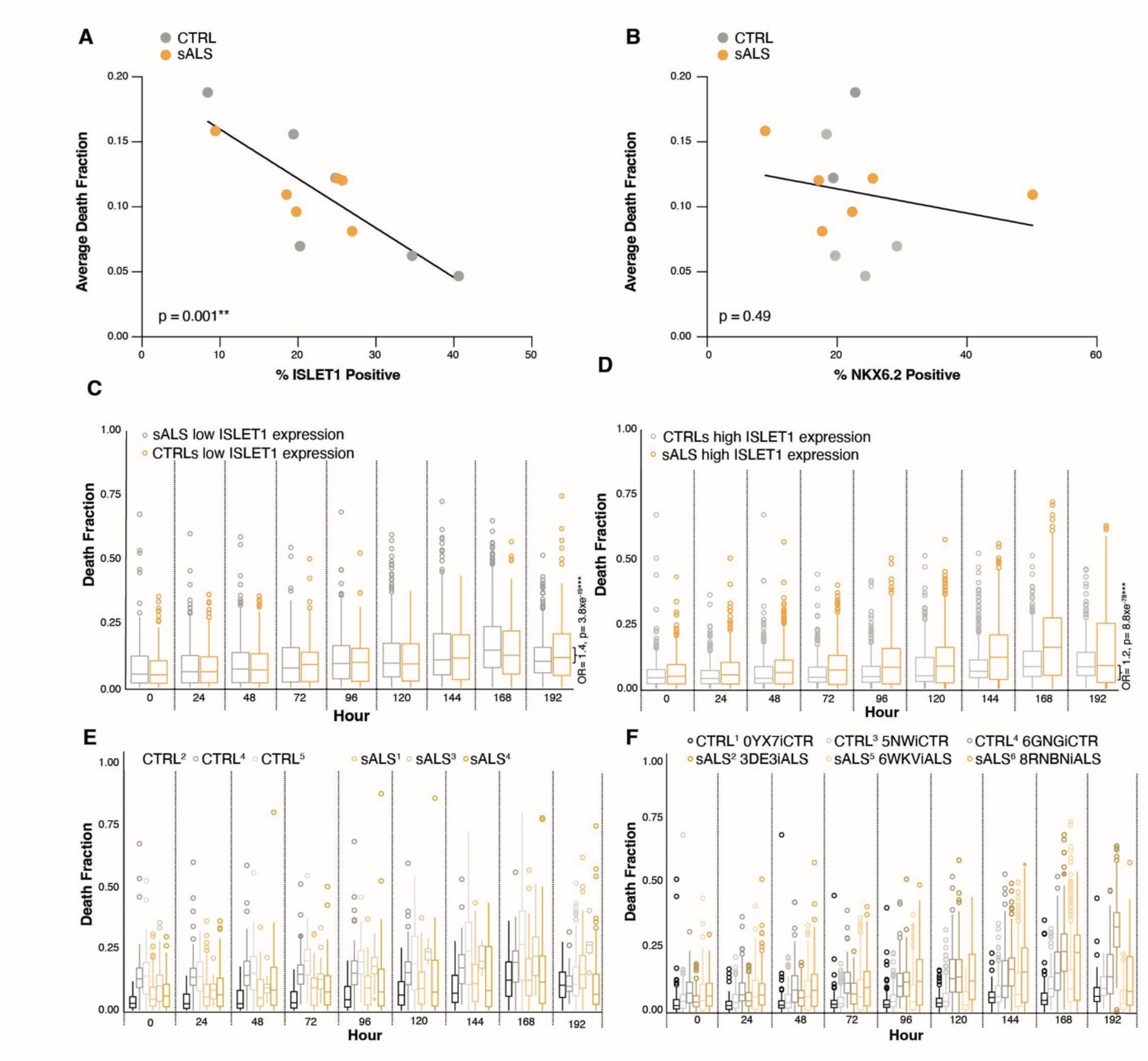


**Supplementary Figure 24. There is a correlation between motor neuron marker expression and cell death in control and sALS iMNs. A**) There is a striking negative correlation (Spearman Correlation, r = -

0.82, p = 0.001) between the percentage of ISLET1-positive motor neurons and the average cell death fraction, indicating that higher ISLET1 expression is associated with reduced cell death. **B**) There is no correlation between the percentage of NKX6.2-positive cells and average cell death fraction (from **Figure 7** and **Supplementary Figure 23**) across controls and sALS iMNs (r= -0.22, p = 0.49). **A B**) Each point represents an individual cell line; grey = controls, yellow = sALS. **C-F**) GEDI-expressing control and sALS iMNs were compared for cell death but stratified according to ISLET1 expression level. **C**) sALS iMNs that display low ISLET1 expression have a higher OR-CD than controls. OR = 1.4, p = 3.9x e-[19]. **D**) Whereas iMNs that display high ISLET expressing iMNs also display a less severe OR-CD for sALS iMNs compared to controls, OR = 1.2, p = 8.8x e-[78]**. E)** Fraction of dead cells plotted per cell line from **C**. **F**) Fraction of dead cells plotted per cell line from **D**. See **Supplementary Figure 23** for number of cells and experiments.

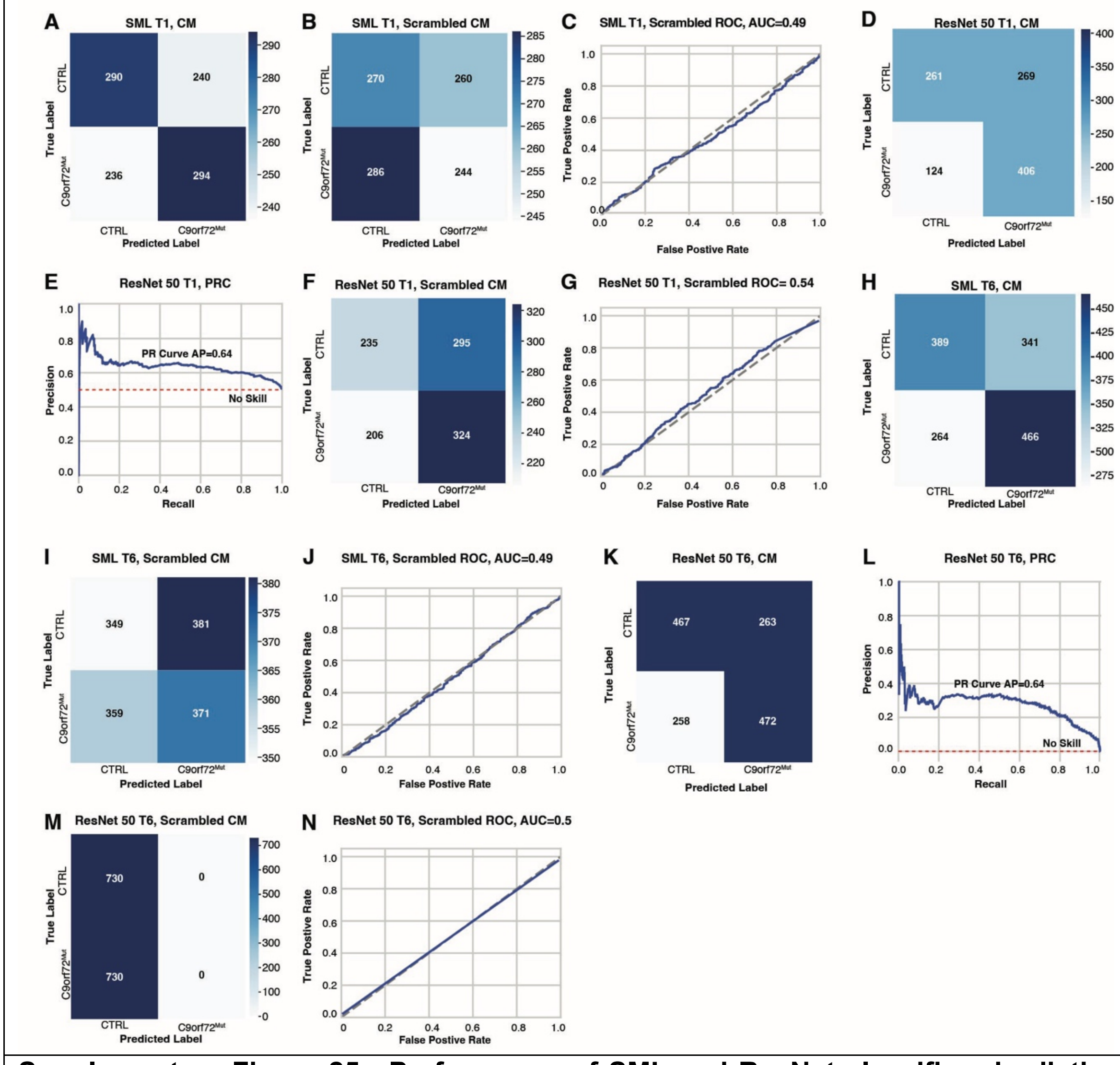


**Supplementary Figure 25. Performance of SML and ResNet classifiers in distinguishing C9orf72$^{mut}$ iMNs vs. control iMNs at early (T1) and later (T6) time points**. **A)** CM illustrating the performance of the

SML models in distinguishing C9orf72$^{mut}$ from control iMNs at T1. The model shows notable cross-class misclassification, with many C9orf72$^{mut}$ crops predicted as CTRL and vice versa. A total of 240 CTRL samples were FP and classified as C9orf72$^{muts}$ while 236 C9orf72$^{muts}$ crops were FNs and incorrectly predicted as CTRL. **B**) CM showing model predictions after random permutation of class labels in the SML T1 condition. Correct and incorrect classifications are distributed approximately evenly across classes, consistent with chance-level performance and indicating the absence of meaningful class-associated signal following label scrambling. **C**). ROC curve corresponding to the SML T1 model trained with randomized labels show an AUC of 0.5, indicating chance-level discrimination. **D**) CM shows that the ResNet50 classifier performs better than the SML model. The model shows moderate discriminative ability but still has substantial cross-class confusion. There were 124 FPs and 269 FNs, indicating difficulty distinguishing control iMNs while it preforms better discriminating C9orf72$^{muts}$. **E**) PRC for the ResNet50 classifier at T1, demonstrating an AP of ~0.64. The curve shows moderate precision across a broad recall range, reflecting limited threshold-independent discrimination. The dashed line represents the no-skill baseline. **F**) CM summarizing ResNet50 predictions after random permutation of class labels in the SML T1 condition. **G**) ROC curve corresponding to the ResNet50 T1 model trained with randomized labels. **H**) CM for the SML T6 model shows improved classification compared to T1. The SML T6 model correctly classified 389 CTRL samples and 466 C9orf72$^{mut}$ samples. Misclassifications were higher for 341 CTRL samples predicted as C9orf72$^{mut}$ as there were less FNs with 264 C9orf72$^{mut}$ samples predicted as CTRL. **I**) CM showing model predictions following random permutation of class labels in the SML T6 condition. **J**) ROC shown that scrambled labels demonstrates that model performance is at chance level for the SML T6 model. **K**) CM of the ResNet50 model trained on T6 images shows similar performance to the T1 ResNet50 model. **L**) PRC for the T6 ResNet50 sALS classifier, achieving an AP of ~0.64 with similar recall relative to T1. **M**) CM summarizing ResNet50 predictions after random permutation of class labels in the SML T6 condition. **N**) ROC curve corresponding to the ResNet50 T6 model trained with randomized labels. Color intensity reflects the number of samples per classification outcome, with lighter colors indicating higher counts. A continuous color scale bar to the right denotes sample count magnitude. For all PRCs, the red dashed line represents the expected performance of a classifier making random predictions. Higher precision values at lower recall levels demonstrate strong model performance, with a gradual decline as recall approaches.

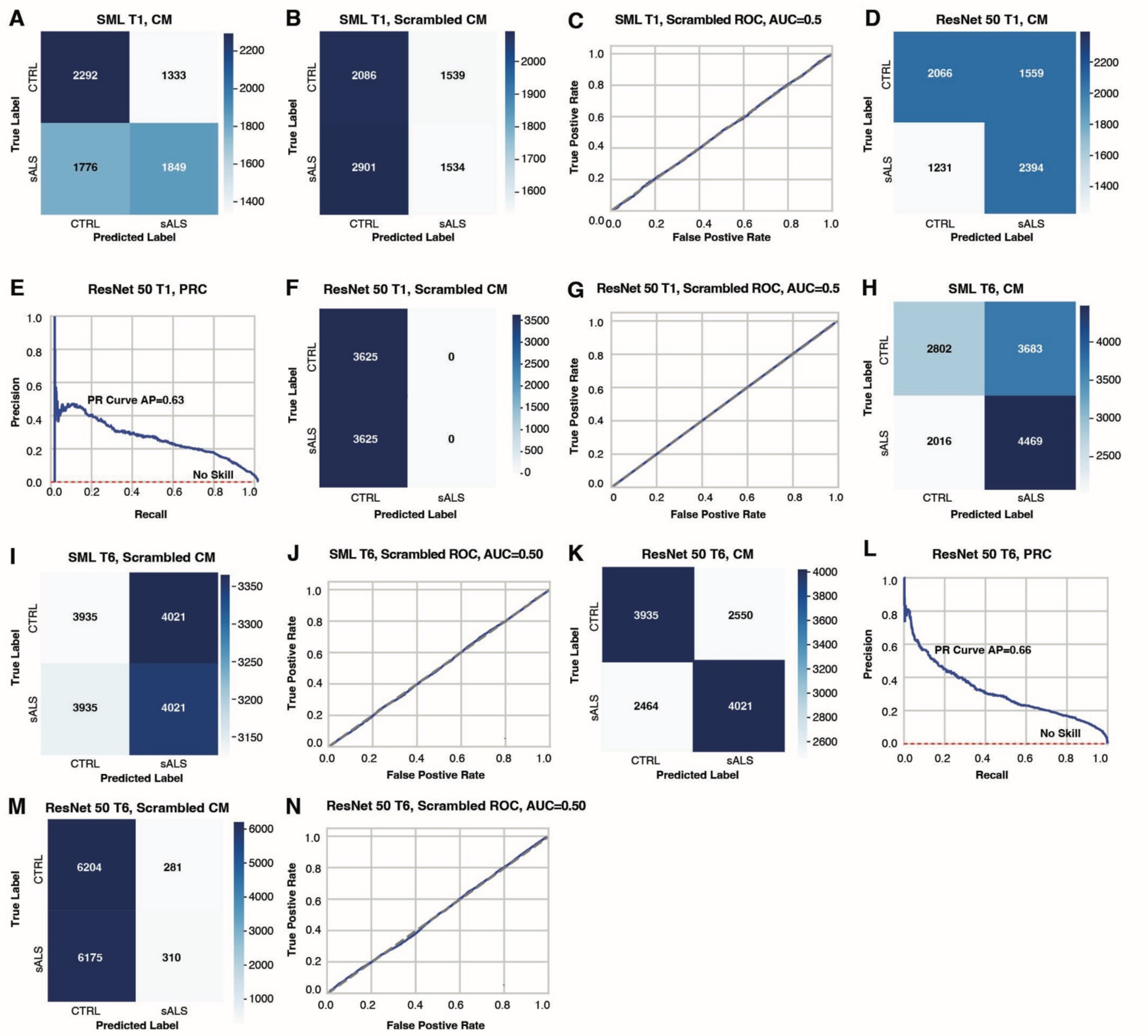


**Supplementary Figure 26. Performance of SML and DNN ResNet classifiers in distinguishing sALS iMNs vs. control iMNs at early (T1) and later (T6) time points. A)** CM showing precision discriminating sALS from control iMNs at T1 using SML. The model shows cross-class misclassification, with many sALS crops predicted as CTRL and vice versa. The model correctly classified 1,849 sALS samples (TPs) and 2,292 CTRL samples (TNs). However, 1,333 CTRL samples were incorrectly predicted as sALS, representing FPs, while 1,776 SALS samples were misclassified as CTRL, representing FNs. **B**) CM showing model predictions on sALS vs CTRL after random permutation of class labels in the SML T1 condition indicate a loss of meaningful class discrimination, supporting chance-level model performance under scrambled-label conditions. Correct and incorrect classifications are distributed approximately evenly across classes, consistent with chance-level performance and indicating the absence of meaningful class-associated signal following label scrambling. **C**). ROC curve corresponding to the SML T1 model trained with randomized labels show an AUC of 0.5, also indicating chance-level discrimination. **D**) CM shows that the ResNet50 classifier performs better than the SML model. The model shows moderate discriminative ability (with 2,394 TPs and 2,066 TNs) but still has substantial cross-class confusion. There were 1,559 FPs and 1,231 FNs, indicating difficulty distinguishing control iMNs while it preforms better discriminating sALS. **E**) PRC for the ResNet50 classifier at T1, demonstrating an AP of ~0.63. The curve shows moderate precision across a broad recall range, reflecting limited threshold-independent discrimination. The dashed line represents the no-skill baseline. **F**) CM summarizing ResNet50 predictions after random permutation of sALS and CTRL labels in the T1 condition. **G**) ROC curve corresponding to the ResNet50 T1 model trained with randomized labels. **H**) CM for the SML T6 model shows improved classification compared to T1. The SML T6 model correctly classified 2,802 CTRL samples and 4,469 sALS samples. Misclassifications were higher with 3683 CTRL samples predicted as sALS as there were less FNs with 2,016 sALS samples predicted as CTRL. **I**) CM illustrating the loss of structured classification after class labels were randomly reassigned in the SML T6 condition. **J**) ROC analysis confirms that, with label scrambling, the SML T6 model performs no better than random guessing. **K**) CM of the ResNet50 model trained on T6 images shows better performance to the T1 ResNet50 model, however misclassifications included 2,550 CTRL samples incorrectly predicted as sALS (FP) and 2,464 sALS samples misclassified as CTRL (FNs). **L**) PRC for the T6 ResNet50 sALS classifier, achieving an AP of ~0.66 with a slightly higher recall than for T1. **M**) CM summarizing ResNet50 predictions after random permutation of class labels in the SML T6 condition. **N**) ROC curve corresponding to the ResNet50 T6 model trained with randomized labels. Color intensity reflects the number of samples per classification outcome, with lighter colors indicating higher counts. A continuous color scale bar to the right denotes sample count magnitude. For all PRCs, the red dashed line represents the expected performance of a classifier making random predictions. Higher precision values at lower recall levels demonstrate strong model performance, with a gradual decline as recall approaches.

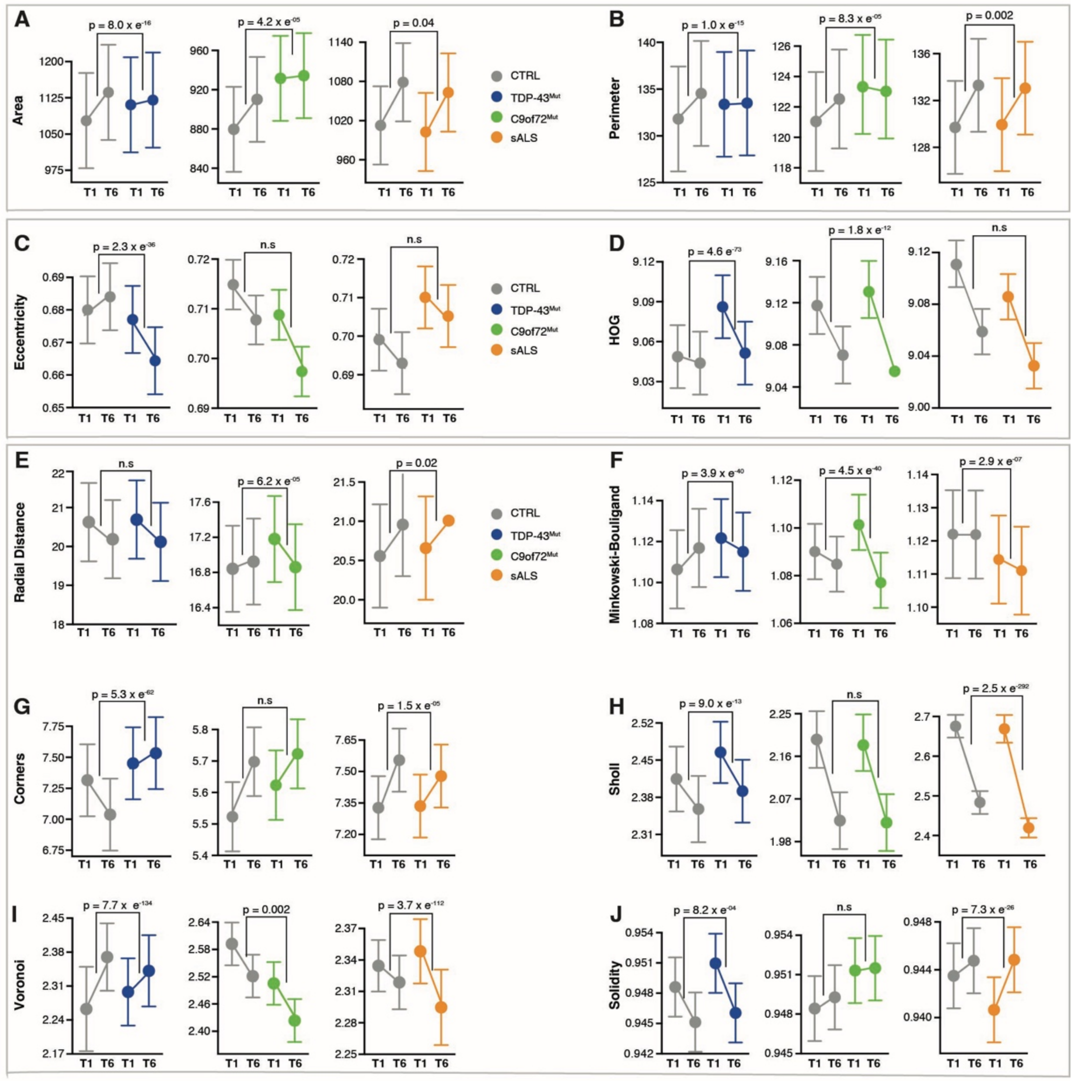


**Supplementary Figure 27. MCOT plots for features capturing size, shape and complexity show robust difference across all ALS groups.** Quantification of morphological feature changes from T1 →T6 (~DIV 28-33) in control (CTRL, grey), TDP-43mut (blue), C9orf72$^{mutant}$ (green), and sALS (orange) iMNs. Each panel shows mean ± SE for the indicated feature at T1 and T6, with brackets denoting the significance of the time-by-disease interaction term derived from a LMM model called RMeDPower2[110]. p values are indicated above brackets and n.s. denotes not significant. Plots here are not log- transformed however, all features were log-transformed before statistical analysis was performed. **A**) Cell area and **B**) perimeter captured size-related changes over time. **C**) Eccentricity, a measure of cell shape that reflects deviation from circularity (higher values indicate more elongated morphologies. **D**) HOG captures local intensity gradients and edge information associated with cellular outlines and neurite structures. **E**) radial distances measuring the average distance from the cell centroid to object boundaries. **F**) Minkowski–Bouligand fractal dimension quantifying cellular complexity that reflects structural richness and branching patterns. **G**) Corners captures regions of high curvature or angularity associated with complex cellular geometries. **H**) Sholl, quantifies the number of intersections between neurites and concentric shells centered on the soma, serving as a proxy for neurite outgrowth and arborization. **I)** Voronoi features, capturing spatial organization and local packing of cellular structures. **J**) solidity, reflecting cellular compactness by comparing object area to its convex hull, with lower values indicating increased fragmentation or irregular boundaries.

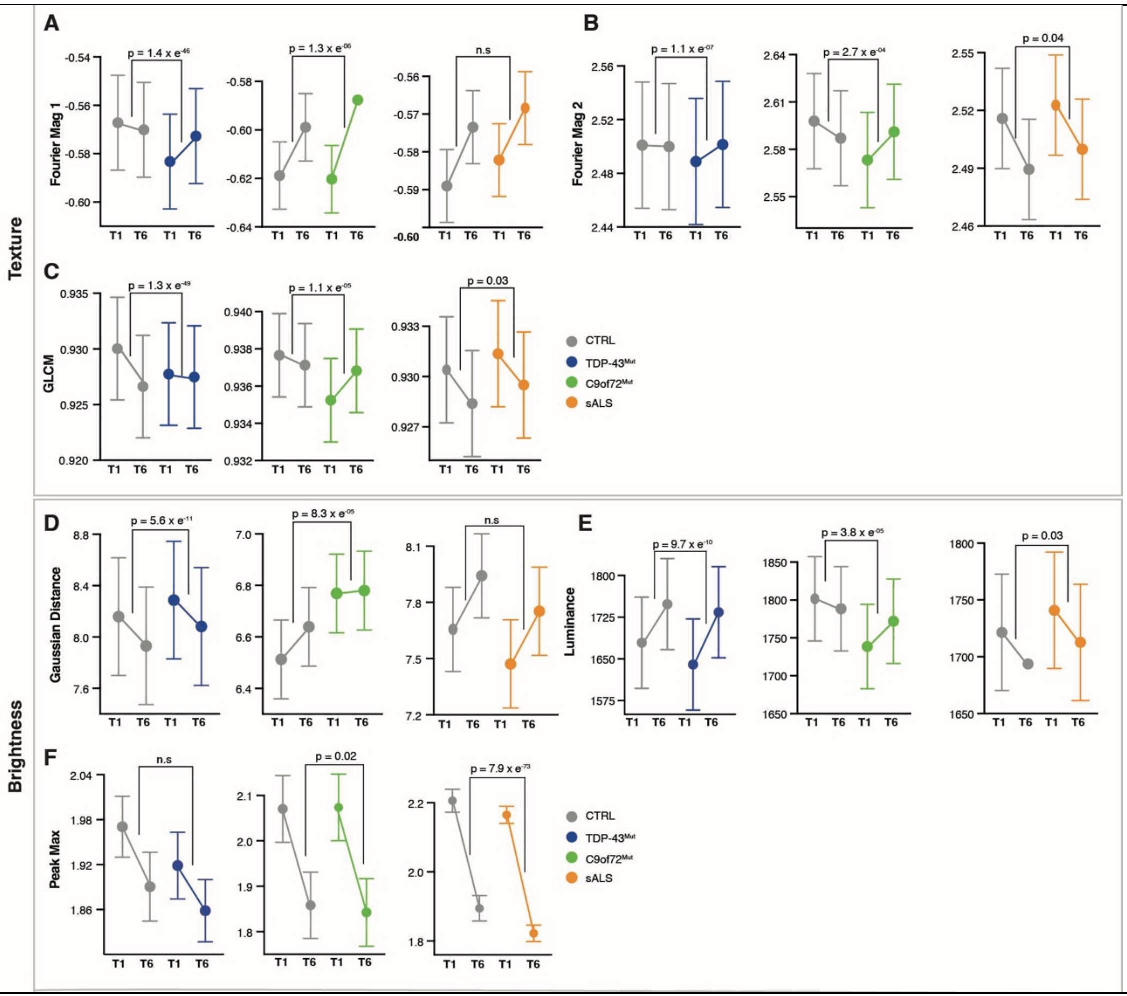


**Supplementary Figure 28. MCOT plots for texture and brightness across all ALS subtypes and controls** Longitudinal analysis of texture- and brightness-related morphological features from T1 →T6 (~DIV 28-33) in control (CTRL, grey), TDP-43mut (blue), C9orf72$^{mutant}$ (green), and sALS (orange) iMNs. Points represent group means ± SD. Statistical significance reflects the time-by-disease-status interaction term derived from linear mixed-effects models LMM model called RMeDPower2[110]. Plots here are not log-transformed however, all features were log- transformed before statistical analysis was performed. **A–B**) Fourier magnitude features (Fourier mag 1 and Fourier mag 2), frequency-domain texture descriptors capturing periodicity and spatial organization of pixel intensity patterns within each crop. These features reflect fine-scale textural changes associated with intracellular organization and structural heterogeneity.

C) Gray-Level Co-occurrence Matrix (GLCM) features, which quantify second-order texture by measuring spatial relationships between neighboring pixel intensities, providing information about local contrast, homogeneity, and pattern regularity. **D**) Gaussian distance, capturing spatial distribution of pixel intensities relative to the object center and reflecting changes in intracellular intensity dispersion. **E**) Luminance, measuring overall brightness intensity of cellular objects and capturing changes in signal intensity over time.

**F**) Peak maximum, representing the highest pixel intensity within each crop and reflecting focal changes in brightness or signal enrichment.

### Supplementary Tables

| iPSC name | iPSC identifier | ALS Mutation | Disease status | Sex | Age At sample acquisition | AO | Source |
|---|---|---|---|---|---|---|---|
| CTRL[1] | 0YX7iCTR | N/A | Unaffected | M | 49 | N/A | Answer ALS/Cedars[93] |
| CTRL[2] | 2AE8iCTR | N/A | Unaffected | F | 50 | N/A | Answer ALS/Cedars |
| CTRL[3] | 5NWTiCTR | N/A | Unaffected | F | 53 | N/A | Answer ALS/Cedars |
| CTRL[4] | 6GNGiCTR | N/A | Unaffected | F | 55 | N/A | Answer ALS/Cedars |
| CTRL[5] | 6ZKZiCTR | N/A | Unaffected | M | 59 | N/A | Answer ALS/Cedars |
| CTRL[6] | 7PFJiCTR | N/A | Unaffected | F | 42 | N/A | Answer ALS/Cedars |
| CTRL[7] | CS2518n2 | N/A | Unaffected | M | 71 | N/A | HD iPSC Consortium[96] |
| CTRL[8] | 9BP3iCTR | N/A | Unaffected | F | 48 | N/A | Answer ALS/Cedars |
| sALS[1] | 0XLRiALS | None | ALS | F | 50 | 49 | Answer ALS/Cedars |
| sALS[2] | 3DE3iALS | None | ALS | F | 45 | 43 | Answer ALS/Cedars |
| sALS[3] | 4HUAiALS | None | ALS | F | 68 | 66 | Answer ALS/Cedars |
| sALS[4] | 5GBQiALS | None | ALS | M | 53 | 52 | Answer ALS/Cedars |
| sALS[5] | 6WKViALS | None | ALS | F | 46 | 46 | Answer ALS/Cedars |
| sALS[6] | 8RBNiALS | None | ALS | M | 41 | 40 | Answer ALS/Cedars |
| sALS[7] | 9DT1iALS | None | ALS | F | 49 | 47 | Answer ALS/Cedars |
| sALS[9] | 5FFPiALS | None | ALS | F | 40 | 25 | Answer ALS/Cedars |
| C9[1] | BUUiALS | *858*[†] | ALS | F | 62 | 60 | Answer ALS/Cedars |
| C9[2] | 6ZLDiALS | Expanded* | ALS | F | 64 | 62 | Answer ALS/Cedars |
| TDP-43[M337V] | GNETARDBPM337V | *TARDBP* M337V | N/A | M | N/A | N/A | PGPC[95] |
| TDP-43[Q331K] | GNETARDBPQ331K | *TARDBP* Q331K | N/A | M | N/A | N/A | PGPC |
| TDP-43[A382T] | NN5320 | *TARDBP* A382T | ALS | F | 62 | NA | NINDS |
| PGP-CTRL[1] | NA | None | N/A | M | N/A | N/A | PGPC |
| PGP-CTRL[2] | NA | None | N/A | M | N/A | N/A | PGPC |
| GC-CTRL[1] | NN5319 | GC control for NN5320 | ALS | F | 62 | NA | NINDS |

**Supplementary Table 1**. Summary of the patient, control and gene-edited lines used. [†] The repeat in *C9ORF72* was estimated by ExpansionHunter[97]. * The *C9ORF72* expansion was confirmed to be expanded by non- CLIAA certified repeat-primed PCR data[98].

| Antibody | Manufacturer | Catalog # | Dilution |
|---|---|---|---|
| MAP2 | ABCAM | ab5392 | 1:1000 |
| Islet-1 | ABCAM | ab109517 | 1:250 |
| NKX6.1 | ABCAM | 221549 | 1:500 |
| FOXG1 | ABCAM | ab18259 | 1:100 |
| SMI/NFH | Biolegend | 801601 | 1:1000 |
| TUJ1 | R&D | MAB1195 | 1:2000 |
| Lamin B1 | ABCAM | ab16048 | 1:2000 |
| SUN2 | ABCAM | ab124916 | 1:150 |
| Calnexin | ABCAM | ab22595 | 1:1000 |
| GM130 | ABCAM | ab52649 | 1:100 |
| NeuN (rabbit) | ABCAM | ab177487 | 1:300 |
| NeuN (mouse) | Millipore | MAB377 | 1:50 |
| TDP-43 | Cell Signaling | E2G6G | 1:100 |
| Goat anti-chicken secondary | Invitrogen | A21449 | 1:1000 |
| Donkey anti-rabbit secondary | Abcam | 150074 | 1:1000 |
| Goat anti-mouse secondary | Invitrogen | A21422 | 1:1000 |
| Goat anti-rat secondary | ABCAM | ab150159 | 1:1000 |

**Supplementary Table 2**. List of all antibodies and dilutions used in this study.
